# Presynaptic filopodia form kinapses and modulate membrane mechanics for synchronous neurotransmission and seizure generation

**DOI:** 10.1101/2024.10.07.616970

**Authors:** Akanksha Jain, Jana Kroll, Jack F. Webster, Jon Moss, Sila K. Ultanir, Alfredo Gonzalez-Sulser, Christian Rosenmund, Michael A. Cousin, Daniela Ivanova

## Abstract

The structural stability of synapses directly contrasts with their functional plasticity. This conceptual dichotomy is explained by the assumption that all synaptic plasticity is generated via either electrical and/or biochemical signaling. Here, we challenge this dogma by revealing an activity-dependent presynaptic response that is physical in nature. We show that dynamic filopodia emerge during action potential discharge and transiently deform synaptic boutons to enhance connectivity. Filopodia generation requires neuronal activity, calcium and actin, and occurs in intact brain circuits and human brain. Mechanistically, their extension preserves synchronous neurotransmitter release by increasing presynaptic membrane tension. However, filopodia generation becomes maladaptive during dysregulated brain activity, exacerbating seizures *in vivo*. Therefore, we provide direct evidence that presynaptic mechanical forces determine the extent and timing of synaptic signals.

## Main Text

Sustained synaptic transmission relies on the recycling of neurotransmitter-filled synaptic vesicles (SVs) (*1, 2*), a process that involves a continuous and often extremely rapid flux of membrane material at the presynapse (*3, 4*). Despite this constant flow of membranes, the prevailing view is that synaptic boutons preserve their structural integrity over long periods of time (days and months) (*5, 6*). The only reported structural plasticity at presynapses involves bouton formation/elimination, and modest size variations (*5, 7*), processes that can be enhanced by synaptic activity and in turn modulate synaptic strength. The limited evidence for activity-dependent structural plasticity within mature synaptic boutons contrasts with the established role of slow, large-scale axonal expansion in circuit refinement during development and in several epilepsy models (*8–15*).

Any structural changes result in a redistribution of forces across the cell membrane and the underlying actin cytoskeleton, thereby influencing membrane tension (*16, 17*). Membrane tension, defined as the membrane’s ability to resist deformations, is emerging as a global coordinator of cell and tissue physiology (*18–21*). Its regulation at the presynapse is particularly pertinent, as tension plays a key mechanical role in the reciprocal control of exocytosis and endocytosis (*22–24*). In this context, elevated membrane tension accelerates exocytosis (*25–29*). Conversely, repeated SV fusion at release sites reduces membrane tension (*30*), weakening the mechanical force driving exocytosis and depressing release rates. SV endocytosis is believed to mitigate this by retrieving SV membranes post-fusion (*30*). However, high-capacity SV endocytosis modes are comparatively slower than exocytosis (*31, 32*). Furthermore, the membrane retrieval efficiency of ultrafast endocytosis (*4*) during repeated fusion, especially at high firing rates, remains uncertain. This indicates a deficit in existing models of presynaptic plasticity that describe the coupling of SV exocytosis and endocytosis, suggesting that crucial physiological mechanisms may remain undetected.

Here, we present data challenging the concept that synaptic boutons are structurally stable elements of the nervous system. Using high-resolution, fast optical imaging of synaptic boutons during neurotransmission, combined with “zap-and-freeze” (*33*) and serial-section transmission electron microscopy (TEM), we reveal that transient presynaptic sprouting is a universal feature of neurotransmitter-releasing synapses. This form of structural plasticity involves localized SV fusion with the tip of dynamically growing and retracting filopodia. Activity-induced filopodia formation is triggered by various neuronal firing rates and enhances neuronal wiring capacity by establishing transient connections, termed kinapses, with postsynaptic structures. Furthermore, functional analyses revealed that filopodia increase presynaptic membrane tension, safeguarding SV fusion rates and the synchronicity of neurotransmitter release during repeated fusion. Finally, we provide *in vitro* and *in vivo* evidence implicating this mechanism in the regulation of seizure activity.

### Presynaptic filopodia – ubiquitous structural remodelling in response to neuronal activity

To assess presynaptic structural changes during synaptic transmission, we labelled the membrane of primary cultures of hippocampal neurons by expressing glycosylphosphatidylinositol-EGFP (GPI-EGFP). For parallel monitoring of SV fusion and retrieval, we co-expressed the red pH-sensitive probe synaptophysin-mOrange2 (SypmOr2), which reports SV fusion as an increase in fluorescence and SV endocytosis as a decrease. High resolution time-lapse imaging revealed that 44% of the presynapses responsive to electrical field stimulation (600 action potentials, AP, at 40Hz), formed extended filopodial protrusions (Movie 1, Fig. 1, A and B). These filopodia were exclusive to active synapses, with an exceptionally high percentage (78%) enclosing SV fusion events within their structure (Fig. 1, A and C).

**Fig. 1.**
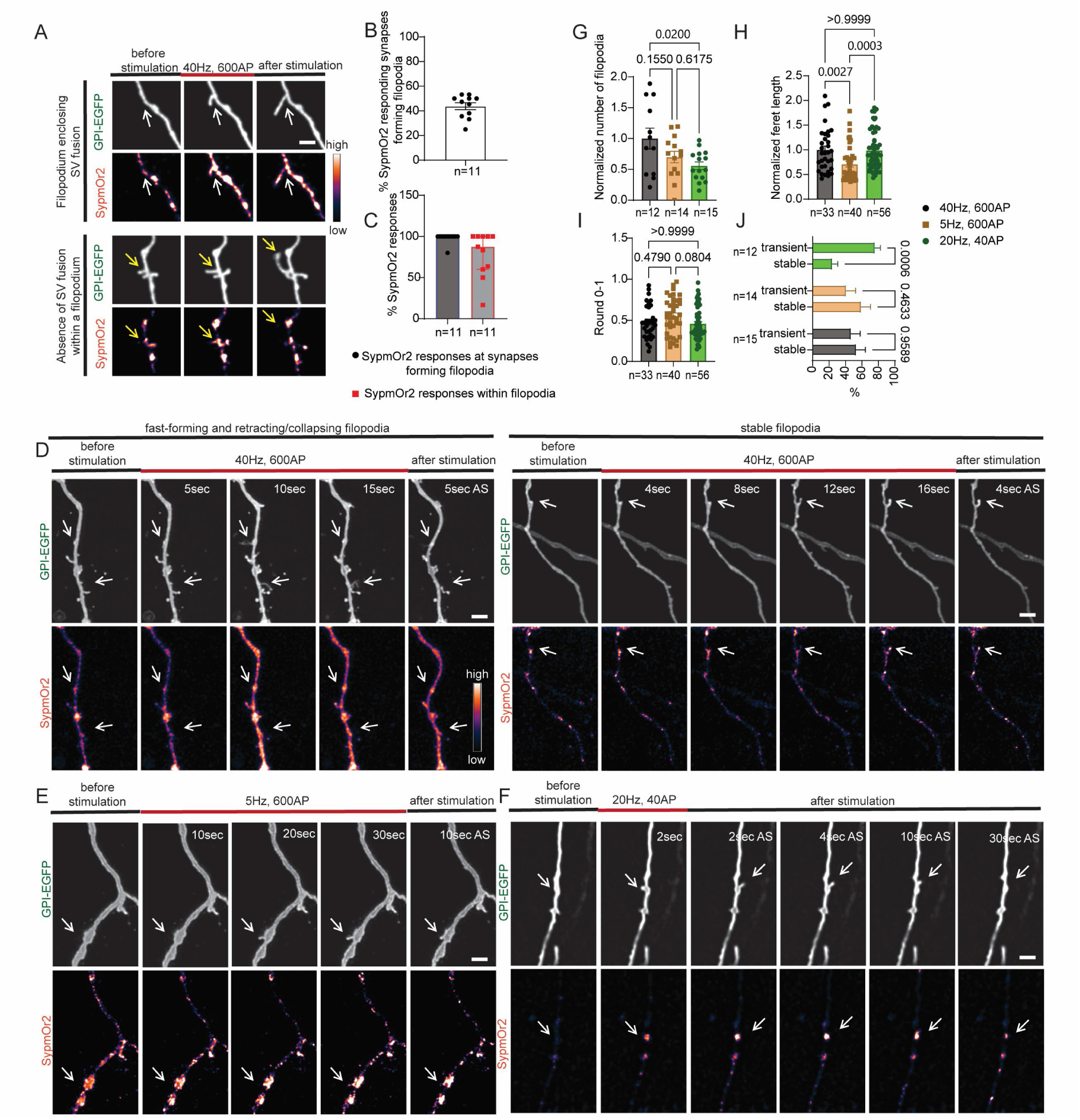
Filopodia form at the presynapse during synaptic activity and enclose SV fusion. (**A-C**) Primary hippocampal neurons were co-transfected with GPI-EGFP and SypmOr2 and stimulated with 600APs (40Hz). (**A**) Representative confocal time-lapse series show filopodia formation at synapses with SypmOr2 fluorescent responses. White arrows, filopodia with SypmOr2 response; yellow arrow indicates filopodia without SypmOr2 response. Scale bar 1 µm. (**B**) Percentage of SypmOr2 responsive synapses forming filopodia. (**C**) Percentage of synapses forming filopodia with Syp-mOr2 responses and percentage of filopodia enclosing Syp-mOr2 responses. (**D-F**) Confocal time-lapse series of synapses forming filopodia in response to (**D**) 600AP at 40Hz, (**E**) 600AP at 5Hz, and (**F**) 40AP at 20Hz. White arrows indicate filopodia-forming synapses. AS, after stimulation. Scale bars 1 µm. (**G**) Normalized filopodia number per 10 µm axon length for 600AP at 40Hz, 600AP at 5Hz, and 40AP at 20Hz. p values, one-way ANOVA with Tukey’s test. (**H**) Normalized filopodia length. p values, Kruskal-Wallis with Dunn’s test. (**I**) Quantification of filopodia shape based on roundness (maximum value of 1 for a circle). p values, Kruskal-Wallis with Dunn’s test. (**J**) Ratio of stable vs transient filopodia. p values, two-way ANOVA with Sidak’s test.

The discovery of these previously undetected alterations in presynaptic structure suggested that these events may be restricted to periods of exceptionally high activity. To determine this, we challenged neurons with two additional stimulation protocols: 600AP, 5Hz, an identical stimulation load but with reduced frequency, and 40AP, 20Hz, which selectively mobilizes the subset of SVs physically attached to the plasma membrane (the readily releasable pool, RRP) (*34*). Both protocols generated filopodia, although with variations in morphology, number, or lifespan (Fig. 1, D to J). At 600AP, 40Hz, 47% of filopodia formed and disappeared during or within 15 seconds (sec) after stimulation, termed transient filopodia (Fig. 1, D and J). This ratio remained consistent at 600AP, 5Hz (Fig. 1, E and J), but substantially increased at 40AP, 20Hz (Fig. 1, F and J), suggesting that prolonged neuronal activity stabilizes presynaptic filopodia. With shorter stimulation trains (40AP, 20Hz), fewer filopodia per axon length were also observed (Fig. 1G). Morphologically, filopodia showed a slight reduction in length at lower stimulation frequencies (600AP, 5Hz) (Fig. 1H), but maintained consistent shape across stimulation paradigms (Fig. 1I). These results suggest that presynaptic filopodia form during repeated SV fusion, irrespective of stimulation load or frequency, with their stability dependent on the duration of neuronal activity.

**Link to Movie 1:** https://doi.org/10.7488/ds/7793

**Movie 1. Activity-induced filopodia form at the presynapse and enclose SV fusion.** Primary cultures of hippocampal neurons were co-transfected with GPI-EGFP and SypmOr2. Neurons were stimulated with 600APs, 40Hz. The movie shows time-lapse series of filopodia formation during stimulation at synapses that display concomitant SypmOr2 fluorescent responses. GPI-EGFP (grey scale) and SypmOr2 (pseudocolour) channels are shown side by side. The periods of stimulation and the time course of the experiment (48 seconds) are indicated. Scale bar 2 µm.

We next evaluated the relationship between neuronal activity and presynaptic calcium on filopodia formation. First, we suppressed neuronal network activity using a combination of the ionotropic glutamate receptor blockers, APV (2-amino-5-phosphonopentanoate) and CNQX (6- cyano-7-nitroquinoxaline-2,3-dione). In parallel, we increased neuronal excitability by treating the neurons with 4-aminopyridine (4-AP), which offers the advantage of inducing stable and spontaneous seizure-like events *in vitro* while also demonstrating robust convulsant properties *in vivo* (*35, 36*).

In these experiments presynaptic membrane dynamics were visualized with the red membrane protein GPI-Apple, with neuronal activity along axons simultaneously monitored via the calcium-sensitive indicator GCaMP6f. No spontaneous calcium spikes or filopodia formation were detected in APV/CNQX-treated neurons (Fig. S1, A and B). In contrast, 4-AP induced propagation of calcium transients that spatially and temporally coincided with a spread of filopodia (Movie 2, Fig. S1, C to G). Importantly, there was a robust temporal coupling between individual calcium spikes and filopodia extension, confirming its activity-dependent nature (Fig. S1, D to F). Furthermore, the comparable ratio of transient versus (vs) stable filopodia in 4-AP-treated neurons (Fig. S1G), with electrical field stimulation confirms this protocol is physiologically relevant.

**Link to Movie 2:** https://doi.org/10.7488/ds/7794

**Movie 2. Temporal coupling of axonal calcium responses and filopodia formation.** Primary cultures of hippocampal neurons were co-transfected with GPI-Apple (red) and GCaMP6f (green) and were incubated with 100 µM 4-AP for 30 min. Live imaging of transfected axons was performed for 5 min in the continuous presence of 4-AP. The movie illustrates the spatio-temporal coupling of filopodia formation and axonal calcium responses as reported by GCaMP6f. The time course of the experiment (5 minutes) is indicated. Scale bar 2 µm.

### Presynaptic filopodia ultrastructure reveals varied morphologies, SV and endosome content

The large dimensions and transient nature of filopodia precludes their ultrastructural visualization through conventional TEM. Therefore, to explore filopodial ultrastructural morphology, we combined “zap-and-freeze” (*33*) of primary hippocampal cultures with serial-section TEM and subsequent 3D reconstructions of axonal segments. The protocol involved electrical field stimulation of primary hippocampal neurons (600AP, 40Hz), followed by high-pressure freezing milliseconds after stimulation. This approach allowed an effective capture of filopodia for ultrastructural characterization.

Eleven presynaptic boutons containing filopodia were 3D reconstructed (Fig. 2). Consistent with our live imaging data, filopodia displayed variable morphology and length, appearing as elongated extensions of the presynapse, each carrying a heterogeneous number of SVs (Fig. 2, D and E). The SVs within the filopodia showed a relatively uniform distribution with only a slightly reduced density near the tip, suggesting possible passive transport from the presynaptic bouton through the filopodia lumen (Fig. 2, D and E). This result supports our observation from live imaging, demonstrating that filopodia carry SVs capable of undergoing fusion within them (Fig. 1, A and C). Interestingly, 73% of the reconstructed filopodia also contained endosomes. Their distribution varied within filopodia (Fig. 2, F and G), with some concentrated at the tip (Fig. 2H, filopodium 1) and others along the filopodial core (Fig. 2H, filopodium 2, 4, 7, 8 and 11) or base (Fig. 2H, filopodium 9 and 10). These individual examples likely represent different stages in the recruitment or generation of endosomes within filopodia. Since endosomes are integral to sorting and directing cellular debris for degradation, we hypothesize that endosomes are a consequence of retrieving the membrane of contracting filopodia.

**Fig. 2.**
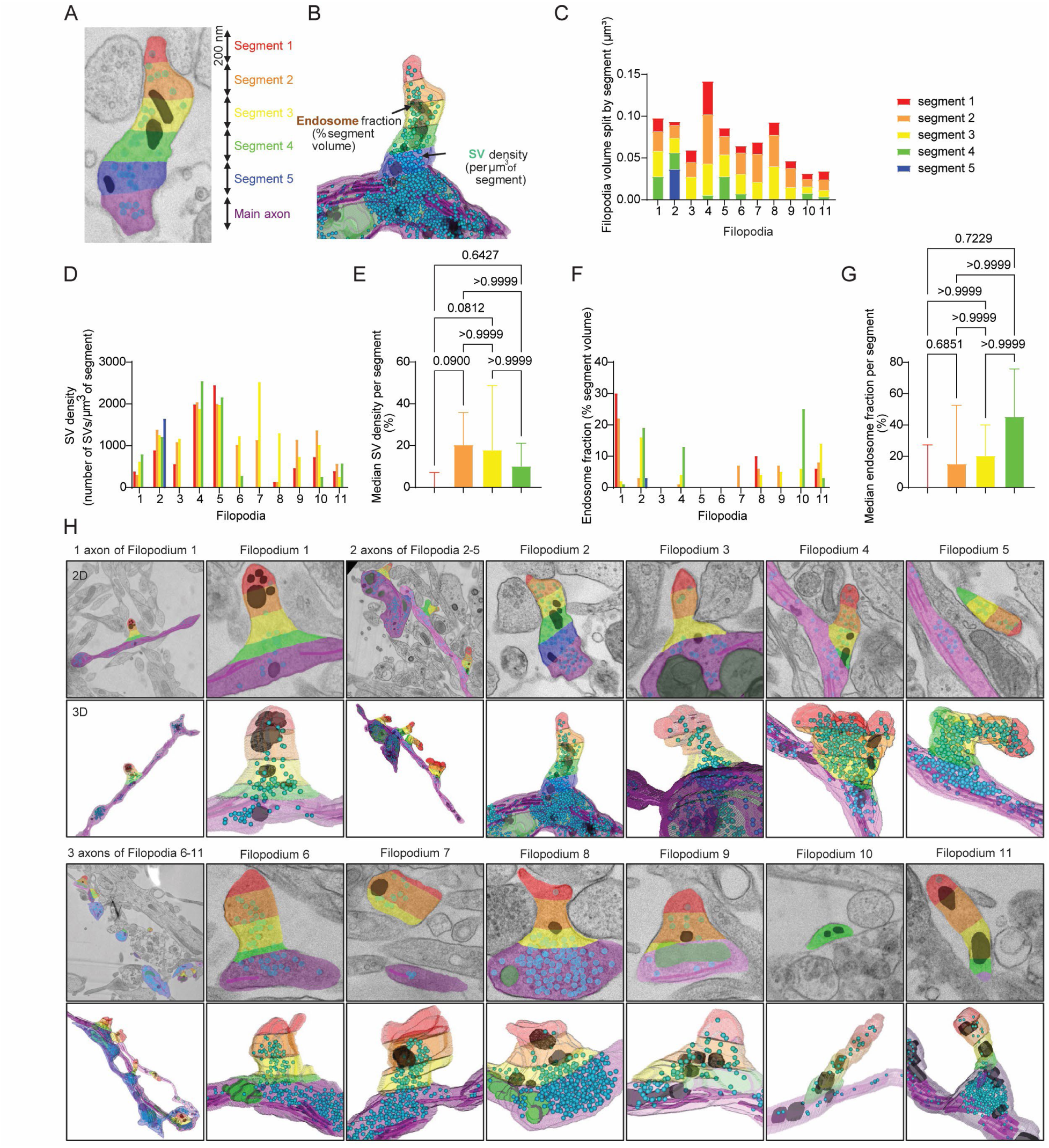
Filopodia show varied morphologies, endosome fractions, and SV densities along their lengths. Primary hippocampal neurons were stimulated with 600AP (40Hz), then subjected to high-pressure freezing and freeze substitution. (**A**) 2D view of a filopodium split into 200 nm segments from tip to main axon. (**B**) 3D reconstruction of a filopodium showing segments, endosomes (brown), SVs (cyan), main axon filaments (magenta) and mitochondria (dark green) from 2D tracings of electron micrographs. (**C**) Segment volumes of eleven filopodia, color-coded as in (**A**). Filopodia under 800 nm lack segment 5; under 600 nm lack segments 4 and 5. (**D**) Segment SV densities (number of SVs per µm³). (**E**) Normalized SV densities in first four segments from tip, averaged across all filopodia. p values, Kruskal-Wallis with Dunn’s test. (**F**) Endosome fraction of filopodia segments (percentage of segment volume). (**G**) Normalized endosome fractions in first four segments from tip, averaged across all filopodia. p values, Kruskal-Wallis with Dunn’s test. (**H**) Representative 2D and 3D images of each filopodium and parent axon from 3 image stacks. Stack 1: one axon with one filopodium (#1); stack 2: two axons (purple, pink) each with two filopodia (#s 2-3, purple; #s 4-5, pink); stack 3: three axons (purple, pink, dark blue) with filopodia #s 6-8, 9-10, and 11 respectively. Segment lengths of 200 nm used for scale, except Filopodium 10 (2D), matched to Filopodium 11.

### Presynaptic filopodia are key in establishing dynamic synaptic connectivity

Filopodia exclusively emerge from synaptic boutons during synaptic activity as indicated by the co-localization with SypmOr2 events (Fig. 1, A and C). To validate their synaptic localization, we investigated their co-localization with two additional presynaptic markers: Bassoon (Bsn), a constituent of the cytomatrix of the active zone, and vGAT, a marker for inhibitory presynapses (Fig. 3, A and B). In parallel, we examined their co-localization with postsynaptic AMPA receptors containing GluA subunits (Fig. 3C).

**Fig. 3.**
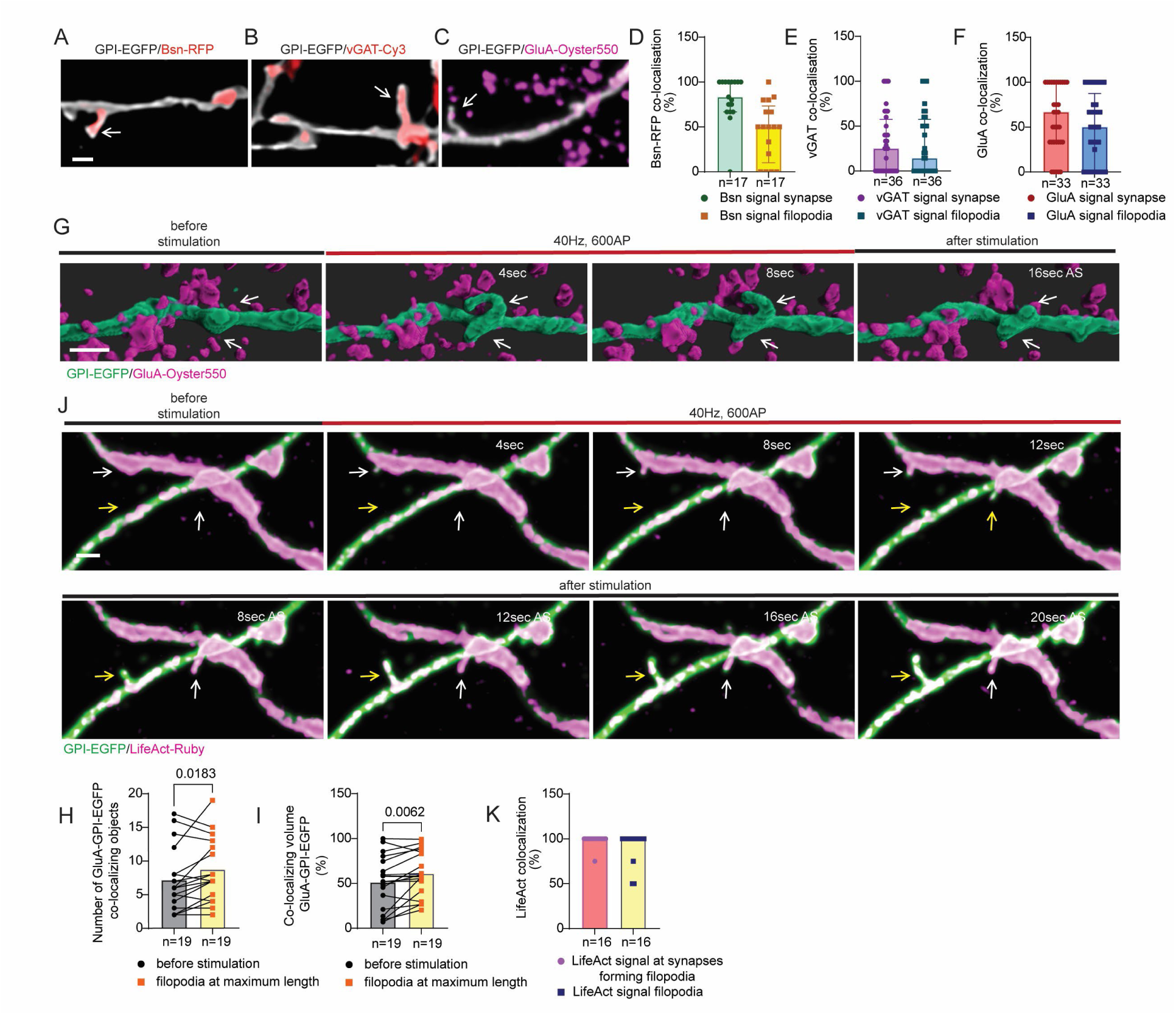
Actin polymerizes in presynaptic filopodia which form transient synaptic contacts (kinapses) with GluA-containing postsynaptic structures. (**A**) Primary hippocampal neurons were co-transfected with GPI-EGFP and Bsn-RFP. (**A-C**) Neurons were transfected with GPI-EGFP, live-labelled with Cy3-conjugated vGAT antibody (**B**) or Oyster550-conjugated GluA antibody (**C**) for 30 min, stimulated with 600APs (40Hz), fixed, and imaged. Representative overlays show filopodia co-localisation with Bsn-RFP (**A**), vGAT-Cy3 (**B**), and GluA-Oyster550 (**C**). White arrows indicate filopodia. Scale bar 1 µm. (**D-F**) Percentage of filopodia-forming synapses and filopodia co-localizing with Bsn-RFP (**D**), vGAT-Cy3 (**E**) and GluA-Oyster550 (**F**). (**G,H,I**) Neurons were transfected with GPI-EGFP and live-labelled with GluA-Oyster550. (**G**) Confocal time-lapse series shows kinapses between filopodia and GluA-labelled structures during 600AP (40Hz) stimulation. White arrows indicate filopodia. Scale bar 1 µm. (**H,I**) Quantification of GluA-GPI-EGFP co-localizing objects at rest and at maximum filopodia extension (**H**) and co-localizing volumes (**I**) p values, paired t-test. (**J**) Neurons were co-transfected with GPI-EGFP and LifeAct-Ruby. Time-lapse series shows actin polymerization in forming filopodia during 600AP (40Hz) stimulation. White arrows indicate filopodia with continuous actin filaments; yellow arrow indicates treadmilling actin filaments. Scale bar 1 µm. (**K**) Percentage of filopodia-forming synapses and filopodia displaying LifeAct-Ruby signal.

First, we performed a post-hoc analysis to assess the co-localization of the synapses from which filopodia emerge and filopodia themselves with the synaptic markers. Analyses revealed that 81% of synapses forming filopodia contained a Bsn signal, with 46% of filopodia showing a Bsn signal along their core (Fig. 3, A and D). A notable portion of synapses forming filopodia were vGAT-positive (30%), and 30% of filopodia contained vGAT along their central axis (Fig. 3, B and E). Furthermore, 59% of synapses forming filopodia were Glu-A positive, and 50% of filopodia overlapped with Glu-A puncta (Fig. 3, C and F). These findings confirm the synaptic origin of filopodia at both excitatory and inhibitory synapses and indicate that they serve as extensions of the presynapse, potentially facilitating connections to postsynaptic structures.

Next, we examined the dynamic 3D co-localization of filopodia with GluA-labelled postsynapses through real-time live imaging during field stimulation (600AP, 40Hz) (Fig. 3, G to I). The assessment of the number of co-localizing GPI-EGFP-GluA objects and volumes showed a statistically significant increase in overlap during filopodia extension (Fig. 3, H and I). This suggests that the formation of filopodia provides the presynapse with the capacity to establish temporary multisynaptic connections. We term these dynamic connections “neuronal kinapses” (a combination of movement (kine-) and fastening (-apse)). Therefore, filopodia formation is a form of short-term structural synaptic plasticity that can transiently modify connectivity in response to AP discharge.

### Filopodia formation requires actin polymerization but is not dependent on SV exocytosis

We have demonstrated that neuronal activity triggers filopodia generation, with SV fusion events detected in most of these structures. However, there were instances of filopodia where no SypmOr2 signal could be detected (Fig. 1, A and C). Therefore, we next assessed whether filopodia formation is dependent on either or both SV fusion and/or calcium influx, via two different strategies that decouple these events. First, we triggered activity- and calcium-independent SV fusion by applying a 5-sec hypertonic sucrose pulse (500mM), which elicits fusion of RRP (*37*). This stimulus triggered a minor enlargement of synapses without the development of elongated protrusions (Fig. S2, A and B), in contrast to RRP mobilization by an electrical field stimulus, where robust filopodia generation was observed (Fig. 1, F to J). Second, we used tetanus toxin (TeTN) treatment to block SV fusion by cleaving synaptobrevin2 (syb2) (Fig. S2, C to H), a key component of the vesicular Soluble N-ethylmaleimide-sensitive factor activating protein receptor complex. Importantly, activity-dependent calcium influx is not impacted by this maneuver. TeTN did not affect filopodia shape but reduced both filopodia number and the average length of individual filopodia (Fig. S2, F to H). Importantly, despite complete blockade of evoked SypmOr2 responses by TeTN, activity-induced filopodia formation persisted unperturbed in some neurons (Fig. S2, E and F). These data demonstrate that SV fusion alone is not sufficient to initiate filopodia formation and point to the critical role of neuronal activity and calcium influx for their emergence. Nevertheless, SV fusion may contribute to filopodia elongation, likely through membrane addition.

The formation of presynaptic protrusions involves extensive plasma membrane deformations, which we reasoned would require a coordinated reorganization of the underlying cytoskeleton. The rearrangement of cortical actin is necessary for the exertion of protrusive forces, that push the membrane forward. To determine if presynaptic filopodia contain a network of actin filaments, we co-expressed LifeAct-Ruby together with GPI-EGFP and visualized actin dynamics during activity-induced filopodia formation. Axons, including putative en-passant synaptic boutons, were filled with filamentous actin (Fig. 3, J and K). LifeAct-Ruby-labelled actin filaments were detected in nearly all filopodia (92%) formed in response to field stimulation (Fig. 3K). Certain actin filaments within the filopodia displayed continuity, seamlessly integrating the filopodial core into the actin meshwork of the synapse (Fig. 3J, white arrows). Conversely, some actin filaments displayed a more intermittent pattern, treadmilling along the filopodial extension trajectory (Fig. 3J, yellow arrows). These results strongly suggest that the elongation of presynaptic filopodia is tightly associated with the *de novo* growth of actin filaments.

To confirm a key role for actin polymerization in filopodia elongation, we disrupted pathways regulating actin dynamics. To this end, we used SMIFH2, an inhibitor of formin-driven actin polymerization which impacts invagination-driven presynaptic clathrin-independent endocytosis (CIE) (*38*). Our array of inhibitors included EIPA and its slightly less potent analogue amiloride, both specifically selected for their ability to target actin-rich membrane ruffles with high selectivity (*39*). To probe the dynamin-dependence of filopodia assembly, we also used dynasore, a potent inhibitor of this GTPase (*40*).

The number of filopodia formed in response to field stimulation served as an indicator of inhibition efficacy (Fig. S3A). Our analyses revealed that SMIFH2 moderately reduced filopodia numbers, whereas EIPA and amiloride virtually abolished their formation (Fig. S3, A and B).

SMIFH2 reduced filopodia length but filopodial shape remained unaffected by the inhibitors (Fig. S3, C and D). Dynasore showed no significant impact on the number or morphology of activity-induced filopodia (Fig. S3, A to D). The interventions also had no significant effect on the ratio of stable-to-transient filopodia (Fig. S3E). Altogether, these data suggest that targeting actin remodelling effectively inhibits presynaptic filopodia formation, with EIPA and amiloride serving as potent eliminators of this process.

EIPA and amiloride inhibit formation of membrane protrusions by blocking most known NHE (sodium-hydrogen antiporter) isoforms (*39, 41, 42*). We tested the extent to which we could replicate their effect by reducing the expression of just one NHE isoform -NHE1 (Fig. S3, F to L). Using an shRNA to selectively reduce NHE1 expression in hippocampal neurons (Fig. S3, F and G) led to a marginal decrease in filopodia formation during synaptic activity (Fig. S3, H and I), with no impact on filopodial length (Fig. S3J), morphology (Fig. S3K), or the ratio of transient-to-stable filopodia (Fig. S3L). This suggests a subtle involvement of NHE1 in regulating membrane protrusions at the presynapse, which contrasted with the broader inhibitory effects observed with compounds like EIPA and amiloride.

To examine synaptic ultrastructure upon treatment with these agents, we performed “zap-and-freeze” (600AP, 40Hz) and prepared samples for conventional TEM. We focused on key presynaptic nanoarchitecture parameters, including SV number and distribution, docked vesicle numbers, SV size, and endosome number (Fig. S3, M to Q). No significant changes in overall synaptic morphology were observed with any inhibitor.

SV cluster density and distribution relative to the active zone as well as SV size distribution remained unchanged between control and treated cells (Fig. S3, N and Q). There was a slight increase in synaptic profiles with more docked SVs under SMIFH, EIPA, and amiloride conditions, which may indicate delayed SV fusion (Fig. S3O). A reduced frequency of synaptic profiles with multiple endosomes was scored, under dynasore (more than three endosomes) and SMIFH, EIPA, and amiloride (more than four endosomes) treatments (Fig. S3P). These findings indicate a possible involvement of filopodia in presynaptic membrane recycling.

### Filopodia formation modulates membrane tension and promotes synchronous synaptic transmission

We next explored the synaptic function of filopodia, exploiting the fact that EIPA eliminated their activity-induced formation (Fig. S3, A and B). We first tested if presynaptic filopodia contribute to acute changes in tension at the presynapse during synaptic activity, as structural deformations are expected to generate membrane tension gradients. While slightly less than half of active synapses form filopodia (Fig. 1B), evidence suggests tension can propagate over several microns in axons (*30*). Thus, we hypothesized that inhibiting filopodia formation could produce detectable changes in membrane tension through this long-range coupling.

To monitor the dynamics of presynaptic tension in real time, we used the molecular probe Flipper-TR which reports changes in membrane tension through alterations of its fluorescence lifetime (Fig. 4 and S4) (*43*). Membrane tension was measured at active synapses (live-labelled with an antibody against synaptotagmin1 (syt1-ATTO647N)) before and during stimulation using fluorescence lifetime imaging microscopy (FLIM) (Fig. 4, A and B). As a positive control, we exposed neurons to a mild hypertonic shock (500 mM sucrose, 5 sec) (Fig. 4, B to D) which decreases membrane tension (*43, 44*) and triggers SV fusion (Fig. S2A). As predicted, the lifetime of synaptic Flipper-TR decreased significantly following the hypertonic pulse (Fig. 4, B to D). However, it showed bidirectional changes in response to electrical field stimulation (Fig. 4A and S4A), resulting in only a slight average variation (Fig. 4C). This observation suggests the existence of robust physiological mechanisms that safeguard membrane tension during neuronal activity.

**Fig. 4.**
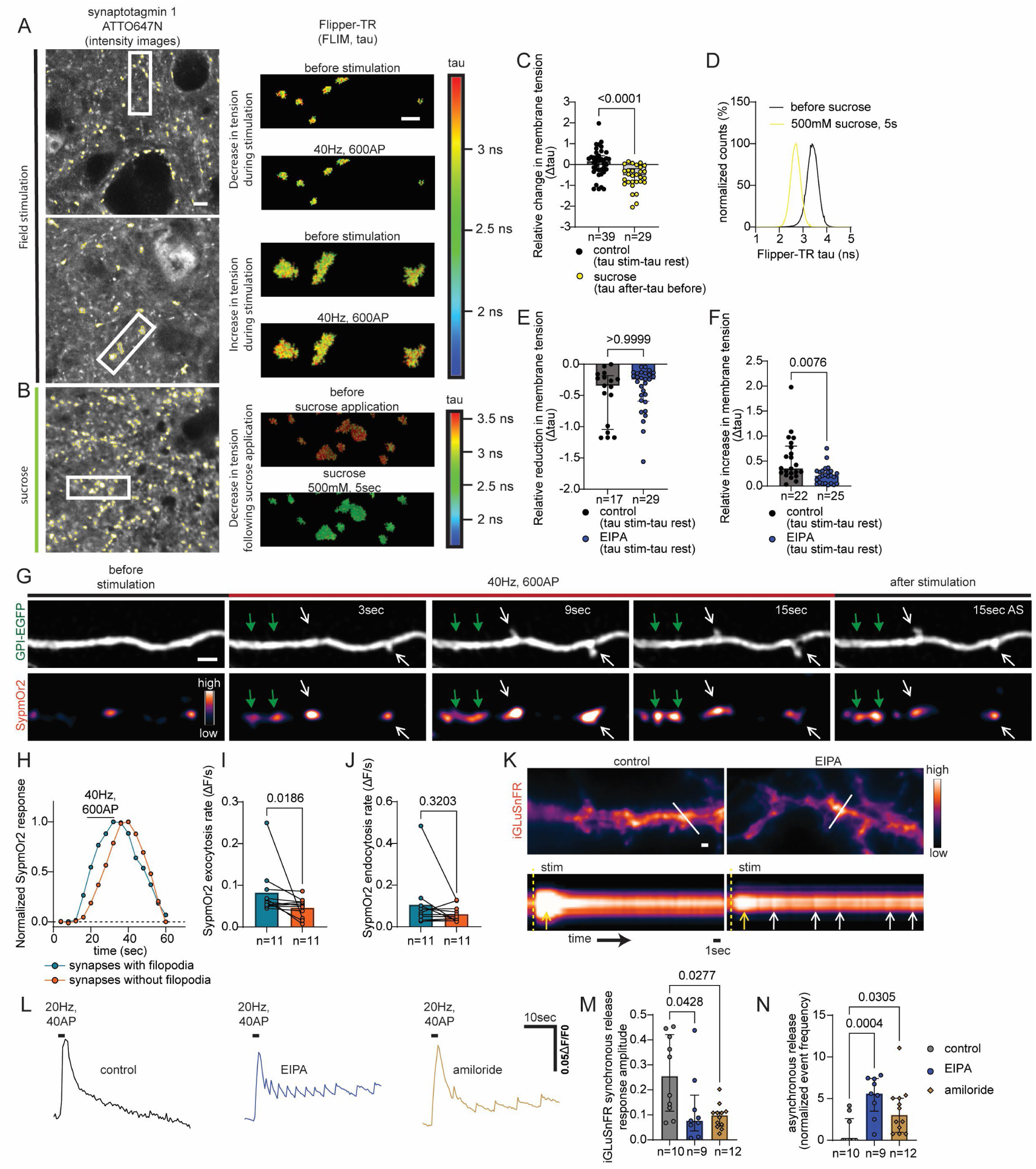
Presynaptic filopodia formation triggers an increase in membrane tension and promotes synchronous SV fusion. Synaptic boutons in primary hippocampal cultures were live labelled with an ATTO 647N-conjugated synaptotagmin1 antibody for 20-30 min. Neurons were incubated with Flipper-TR, with its fluorescence lifetime (tau) monitored using FLIM imaging. (**A, B**) Left: Representative images of synaptotagmin1 fluorescence intensity showing Flipper-TR tau ROIs at rest and during stimulation with either 600APs (40Hz) (**A**) or 500 mM sucrose (**B**). Right: Enlarged Flipper-TR tau ROIs (pseudocolour) at rest and during stimulation. Example images display a reduction (Top) or increase (Middle) of Flipper-TR tau during AP stimulation and reduction (Bottom) after sucrose application. Scale bar: left 5 µm, right 2 µm. (**C**) Average change in Flipper-TR tau (Δtau) before and during stimulation (control cells) or before and after sucrose application. p values, Mann-Whitney U test. (**D**) Distribution of Flipper-TR tau showing the shift from high to low tension after 500 mM sucrose treatment. Analysis from cell shown in (**B**). (**E, F**) Neurons treated and stimulated as in (**A**) were incubated with EIPA (50 µM) or vehicle control for 30 min. Δtau shows average reduction (**E**) or increase (**F**) in Flipper-TR tau during stimulation in control and EIPA-treated cells. p values, Mann-Whitney U test. (**G-J**) Hippocampal neurons co-transfected with SypmOr2 and GPI-EGFP were stimulated with 600AP, 40Hz. SypmOr2 exocytosis and endocytosis kinetics were compared at synapses with and without filopodia. (**G**) Representative images demonstrating the difference in SypmOr2 fusion rates between synapses that form filopodia (white arrows) and those that do not (green arrows). Scale bar 1 µm. (**H**) Peak-normalised mean SypmOr2 responses from a single cell. (**I,J**) SypmOr2 exocytosis (**I**) and endocytosis (**J**) rates (ΔF/s); p values: Wilcoxon test. (**K-N**) Hippocampal neurons transfected with iGluSnFR184S were treated with 50 µM EIPA or 1 mM amiloride for 30 min before stimulation with 40AP (20Hz) during live imaging. (**K**) Example iGluSnFR184S images of dendritic segments. Scale bar 1 µm. Kymographs show synchronous (yellow arrows) and asynchronous (white arrows) iGluSnFR184S responses. (**L**) Example iGluSnFR184S fluorescence responses following stimulation with and without EIPA and amiloride. (**M**) Analysis of iGluSnFR184S synchronous response amplitude. (**N**) Analysis of asynchronous event frequency. (**M, N**) p values, Kruskal-Wallis with Dunn’s test.

Simultaneous increases and decreases in membrane tension during synaptic activity imply a reciprocal link between the two. Indeed, elevated membrane tension promotes SV fusion, while SV fusion, in turn, reduces membrane tension (*25–29*). Importantly, further analysis of these bidirectional changes revealed that EIPA significantly decreased the fraction of synapses that experienced an increase in membrane tension during synaptic activity, without impacting the fraction of synapses at which membrane tension decreased (Fig. 4, E and F; Fig. S4B). Therefore, our data strongly suggest that the increased tension observed in response to neuronal activity is due to filopodia-generated mechanical forces, which may contribute to sustained exocytosis rates during repeated SV fusion.

Both *in silico* and experimental data consistently show that increased membrane tension enhances the efficiency of membrane merging (*26, 28, 29*). To determine the influence of filopodia-generated membrane tension on SV fusion, we directly compared the SypmOr2 responses evoked by electrical field stimulation at synapses with filopodia to those without (Fig. 4, G to J). Synapses displaying filopodia formation showed a significant acceleration in exocytosis compared to those lacking filopodia (Fig. 4, G to I). This is consistent with the hypothesis that membrane protrusions elevate membrane tension to provide mechanical support for SV fusion. In contrast, the rate of endocytosis remained unchanged between synapses with or without filopodia (Fig. 4, H and J).

To support this observation, we next determined whether inhibiting filopodia formation with either EIPA or amiloride could reduce SV fusion rates. This was achieved using the pH-sensitive reporter synaptophysin-pHluorin (SypHy) combined with bafilomycin A1 (*45*) to prevent vesicle reacidification, ensuring that changes in fluorescence exclusively reflected SV exocytosis (Fig. S5A). Neurons were challenged with two consecutive trains of stimuli (40AP and 900AP, 20Hz) that target the RRP and all releasable SVs respectively (Fig. S5, A to C). In control cells, SypHy fluorescence increased synchronously with the stimulation and plateaued, while in treated neurons, the plateau was delayed, more prominently after the second stimulus (Fig. S5, A to C). These results suggest that filopodia inhibition does not block SV fusion, but instead significantly retards and desynchronizes it, an effect that is exacerbated during prolonged neuronal activity.

To directly assess if reduced membrane tension was responsible in delaying SV exocytosis, we perfused neurons with 350 mM sucrose while stimulating with 900AP, 20Hz (Fig. S5, D and E). The mild hypertonic solution triggered minimal SV fusion, as indicated by a small SypHy fluorescence response (Fig. S5D, purple arrow). However, when paired with electrical field stimulation, it effectively mimicked the effect of EIPA and amiloride, slowing the upstroke of the stimulus-evoked SypHy response (Fig. S5, D and E). Therefore, artificially reducing membrane tension via application of hypertonic sucrose phenocopies the impact of filopodia inhibition on SV fusion (Fig. S5, D and E), confirming that the SV fusion rate is highly sensitive to membrane tension and slows under reduced tension.

To evaluate the impact of inhibiting filopodia formation on wider synaptic performance, we monitored both evoked neurotransmitter release and intracellular calcium transients (Fig. 4, K to N, and Fig. S5, F and G). Glutamate release and axonal calcium responses in cultured neurons were measured using the high affinity iGluSnFRA184S variant (Fig. 5, K to N) and GCaMP6f respectively (Fig. S5, F and G) in response to 40AP, 20Hz stimulation. The stimulation train triggered a robust synchronous iGluSnFRA184S response in control neurons, only occasionally followed by asynchronous release (Fig. 4, K to N). In contrast, in neurons treated with EIPA and amiloride, the amplitude of the collective synchronous response to 40AP was significantly reduced (Fig. 4, L and M). Importantly, this was followed by a period of asynchronous release, sometimes lasting many seconds after stimulation (Fig. 4, L and N), in agreement with the SypHy experiments (Fig. S5, A to C). In contrast, inhibition of filopodia formation with EIPA and amiloride did not significantly affect the amplitude of the stimulus-evoked calcium responses, nor did it lead to desynchronization of the calcium transients in axons (Fig. S5, F and G). Taken together, these results reveal that filopodia formation supports synchronous neurotransmitter release, specifically at the level of SV fusion, without affecting the preceding step of calcium influx.

**Fig. 5.**
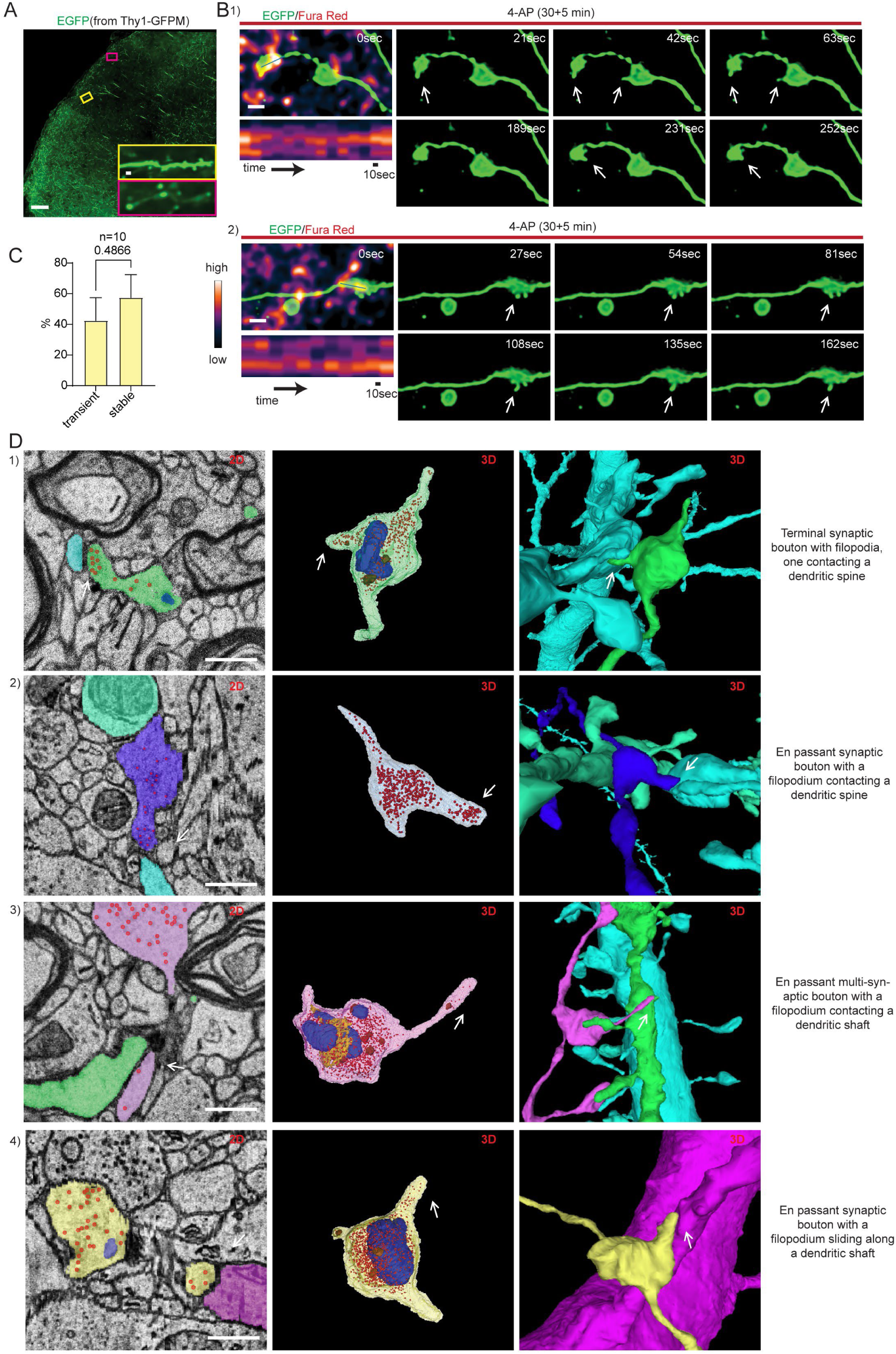
Presynaptic filopodia form rapidly in intact brain tissue and enhance connectivity. (**A-C**) Time-lapse imaging of EGFP-labelled neurons in acute brain slices from Thy1-GFPM mice, loaded with Fura Red and treated with 100 µM 4-AP for 30 min. (**A**) Example image showing cortical region with sparse EGFP expression. Putative axons (purple box) and dendrites (yellow box) identified by morphology. Scale bars: large image 100 µm, inset 2 µm. (**B**) Confocal time-lapse series showing filopodia formation at putative synaptic boutons. Kymographs display Fura Red calcium transients (white arrows, filopodia). Scale bar 1 µm. (**C**) Ratio of stable to transient filopodia. p values, unpaired t-test. (**D**) Representative 2D views (left) and 3D (middle and right) reconstructions of synaptic boutons with putative filopodia from human cerebral cortex. White arrows, filopodia. (Middle) 3D reconstructions of synaptic boutons with filopodia showing SVs (red), endosomes (brown), mitochondria (blue), and endoplasmic reticulum (yellow) from 2D tracings of electron micrographs. (Right) 3D reconstructions of the same filopodia forming contacts with dendritic spines (1 and 2) or dendritic shafts (3 and 4) and thereby enhancing connectivity to a single (1 and 4) or multiple neurons (2 and 3). Scale bars 500nm.

### Presynaptic filopodia form rapidly in intact brain tissue and enhance connectivity

To our knowledge, rapid (occurring within seconds) activity-dependent microscopic-level alterations of presynaptic structure, have not been previously reported in intact brain tissue. Therefore, we used high-resolution time-lapse imaging to visualize the dynamics of EGFP-labelled excitatory synaptic boutons in the superficial cortical layers of acute coronal sections from the Thy1-GFPM mouse line (Fig. 5, A to C). Importantly, the sparse EGFP expression in these regions allowed clear delineation of the outlines of putative en passant and terminal boutons (Fig. 5B). By preloading slices with the calcium indicator dye Fura Red (Fig. 5B), we monitored coarse changes in activity in the imaged boutons. We analyzed the dynamic changes in the shape of synaptic boutons from slices treated with either 4-AP or APV+CNQX (both for 30 min, with an additional 5 min during imaging) (Fig. 5B and S6A). No presynaptic structural changes and no extensive fluctuations in the Fura Red fluorescence signals were observed in slices that had their electrical activity silenced by APV+CNQX (Fig. S6A). In contrast, in 53% of the slices treated with 4-AP, we identified synaptic boutons with dynamic filopodia-like extensions (Fig. 5B), which had a slightly shorter average length than in dissociated neuronal cultures (Fig. S6B). This was usually associated with transient decreases in Fura Red fluorescence, indicating its binding to calcium (Fig. 5B). Thus, rapid presynaptic filopodia formation and retraction in response to changes in activity appear to be a universal phenomenon that also occurs in intact brain tissue.

To ensure presynaptic sprouting was conserved across species and in intact brain, we evaluated the role of filopodia in enhancing neuronal connectivity in the human cerebral cortex. We manually identified and reconstructed the subsynaptic architecture of synaptic boutons with putative filopodia from a recently released resource-a sample of human temporal cortex obtained during surgery to access a hippocampal lesion in an epilepsy patient (*46*). The sample, imaged via serial section EM, contained numerous automatically reconstructed neurons and their synaptic connections. Synaptic boutons with putative filopodia were readily identifiable in the human cortex (Fig. 5D), typically appearing as short extensions (≤500 nm, Fig. 5D, 1), 2) and 4)) deforming the synaptic boutons, while others were longer, likely initiating collateral branches extending from the presynapse (Fig. 5D, 3)). Identical to dissociated neuronal cultures, all putative filopodia contained varying densities of SVs (Fig. 5D, 1) to 4)) and occasionally endosomes (Fig. 5D, 1) and 3)). The putative filopodia established non-classical contacts, with both spiny (Fig. 5D, 1) and 2)) and non-spiny (Fig. 5D, 3) and 4)) postsynaptic targets, in some cases supporting multi-neuron connectivity (Fig. 5D, 3)). Therefore, presynaptic filopodia formation is a conserved mechanism in both mouse and human brain tissue, presumably playing an essential role in the rapid modulation of neuronal connectivity.

### Presynaptic filopodia formation supports synchronous synaptic transmission and exacerbates seizure severity *in vitro* and *in vivo*

To determine if the delay in SV fusion resulting from inhibiting filopodia formation translates into changes in neurotransmission, we investigated how EIPA treatment affected synaptic transmission at CA3-CA1 hippocampal synapses using whole-cell patch-clamp recordings (Fig. 6, A to G). Quantification of excitatory postsynaptic current (EPSC) amplitudes at several stimulus intensities showed no difference between control and EIPA-treated slices (Fig. 6B), indicating that the ability of presynaptic stimuli to evoke postsynaptic responses is not affected by the treatment. Although basal synaptic transmission was preserved (Fig. 6B), we observed a faster decay of the EPSC amplitudes in EIPA-treated slices under high-frequency stimulation (600AP, 40Hz), (Fig. 6, C and D). These results are consistent with the hypothesis that the reduced rate of repeated SV fusion due to filopodia inhibition (Fig. S5, A to C), leads to an accelerated rundown of synchronous synaptic transmission. The high frequency stimulation protocol also triggered asynchronous release (Fig. 6E), which persisted after the presynaptic firing had ceased. While asynchronous release events were detected in control slices, their frequency was markedly increased in EIPA-treated slices (Fig. 6, E and F), whereas their amplitude remained unchanged (Fig. 6, E and G). Altogether, these results provide compelling evidence that during repeated SV fusion, filopodia formation promotes rapid neurotransmitter release in synchrony with APs. Furthermore, we identified a clear inverse correlation between synchronous and asynchronous glutamate release on inhibition of filopodia formation (Fig. 4, L to N and Fig. 6, C to G).

**Fig. 6.**
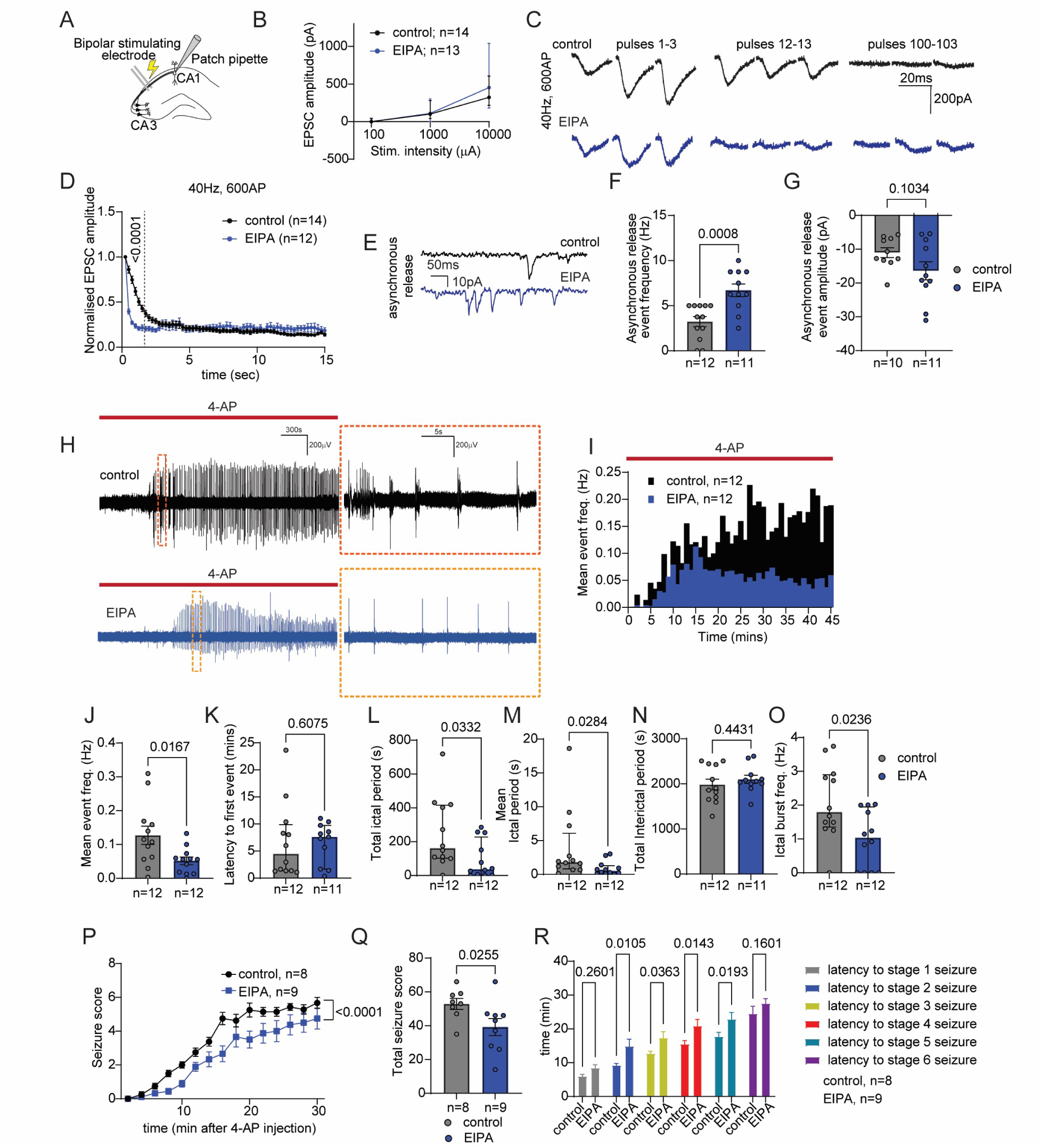
Inhibition of filopodia formation differentially regulates synchronous and asynchronous neurotransmitter release with anti-seizure effects *in vitro* and *in vivo*. (**A-G**) Synaptic transmission at CA3-CA1 synapses assessed by whole-cell patch-clamp recordings in wild-type mouse brain slices with (blue) or without (black) 50 µM EIPA for 30 min. (**A**) Schematic of recording configuration. (**B**) Input-output plot of stimulation intensity vs. EPSC amplitude. (**C,D**) Example EPSC amplitudes (**C**) and synaptic rundown (**D**) during 600AP, 40Hz stimulation. **D**, p values, two-way ANOVA with Sidak’s test. (**E**) Example traces of asynchronous release. (**F, G**) Analysis of asynchronous release frequency (**F**) and amplitude (**G**). p values, unpaired t-test. (**H-O**) Brain slices were treated with 50 µM EIPA for 45 min and epileptiform activity elicited by 100 µM 4-AP was recorded in CA1 pyramidal layer. (**H**) Example traces of interictal and ictal-like discharges. (**I**) Time-frequency histograms of epileptiform discharges during 4-AP wash-in. (**J-O**) Analysis of epileptiform activity: frequency (**J**), latency to first event (**K**), total (**L**) and mean (**M**) duration of ictal-like activity, interictal-like activity duration (**N**), and burst frequency during ictal-like events (**O**). p values: unpaired t-test (**J, N**), Mann-Whitney U test (**K, L, M, O**). (**P-R**) EIPA (5 mg/kg, blue) or vehicle (black) were administered via intraperitoneal injection to 8-10-week-old wild-type mice 30 min before 4-AP (6.65 mg/kg) to induce seizures. (**P**) Time progression of seizure severity post-4-AP injection. p values, two-way ANOVA with Sidak’s test. (**Q**) Total seizure score analysis. p values, unpaired t-test. (**R**) Latency to different seizure stages. p values, two-way ANOVA with Fisher’s LSD.

It can be easily inferred that the timing of synaptic signals and the extent of physical connectivity play a critical role in driving neuronal network synchronization, which is essential for normal brain function (*47*). However, excessive synchronization is observed in several neurological disorders, most prominently in epilepsy (*48*). Therefore, we hypothesized that inhibiting filopodia formation may counteract seizure propagation, in two complementary ways. The first is by limiting connectivity during AP discharge (Fig. 3, G to I; Fig. 5D) and hence, reducing neurotransmitter spread. The second is by increasing asynchronous release, which can limit the extent of seizures by creating periods of reduced synchronous activity.

To test this hypothesis, we first used the 4-AP *in vitro* epilepsy model (Fig. 6, H to O), which replicates the electroencephalographic activity seen in patients with partial epilepsy and successfully emulates the antiepileptic effectiveness of standard anti-seizure medications (ASMs) (*35*). We recorded field potential discharges within the CA1 region of acute brain slices following exposure to 4-AP (Fig. 6H). Two different categories of field potential discharges induced by the drug were detected: ictal-like discharges, within which the spike burst frequency peaked between 0.8 and 3.7Hz, and interictal-like events, encompassing the duration within the recording not classed as an ictal period.

At a concentration that effectively blocked filopodia formation, EIPA was perfused alongside 4-AP and this resulted in a significant reduction in the overall frequency of epileptiform discharges, without affecting their onset (Fig. 6, H to K). A more detailed analysis uncovered that EIPA effectively reduced the total (Fig. 6I) and average ictal period (Fig. 6M) as well as the ictal burst frequency (Fig. 6O), while leaving the total interictal period unperturbed (Fig. 6N). In contrast, the conventional ASM carbamazepine, which we used as a control (Fig. S7, A to H), caused a significant delay in the onset but had no impact on the overall frequency of epileptiform discharges once they commenced (Fig. S7, A to D). Carbamazepine did not alter the total (Fig. S7E) or mean ictal period (Fig. S7F), nor did it affect the total interictal period (Fig. S7G), but it significantly reduced the ictal burst frequency (Fig. S7H).

4-AP is also a highly effective seizurogenic agent in mice (*36*), causing increased activity, clonic limb movements, and sometimes tonic hindlimb extension, along with episodes of wild running and jumping. We evaluated EIPA’s ability to provide protection against acute seizures induced by 4-AP in mice, using an adapted Racine scale (Fig. S7I). Furthermore, we compared its efficacy to that of carbamazepine, which has previously shown protective effects in this model of acute seizures (*36*).

Behavioral seizure responses were scored for 30 min following the administration of 4-AP via a systemic subcutaneous injection. We observed a significant difference in behavioral seizure scores between animals preinjected with EIPA and those receiving a vehicle injection (Fig. 6, P to R). Animals injected with EIPA showed a substantial reduction in seizure intensity throughout the 30-min period following the 4-AP injection (Fig. 6P), and as evidenced by the total seizure score (Fig. 6Q). This diminished sensitivity to 4AP-induced seizures was also evident in the latency to reach stages 2, 3, 4 and 5 seizures (Fig. 6R). Furthermore, EIPA’s protective efficacy was found to be comparable to that of carbamazepine (Fig. S7, J to L).

We conclude that inhibiting presynaptic filopodia formation demonstrates promising anti-seizure potential in both 4-AP *in vitro* and *in vivo* seizure models. While producing similar behavioral outcomes to the classic ASM carbamazepine, EIPA showed a unique effect on a neuronal population level. This finding is significant, given the prevalence of pharmacoresistant epilepsy and the urgent demand for novel approaches targeting diverse mechanisms.

## Discussion

In this study we demonstrate that rapid, transient structural rearrangements of synaptic boutons are a common phenomenon driven by neuronal activity. These structural changes arise from activity-induced reorganization of the actin cytoskeleton, leading to the formation of dynamically expanding, retracting, and collapsing filopodia. SV fusion occurs within filopodia, which also establish connections with postsynaptic structures, thereby enhancing the neuronal wiring plasticity during AP discharge. As a result of these observations, we have termed these transient contacts kinetic synapses - kinapses. Furthermore, we show that these protrusions elevate membrane tension at the presynapse, a critical mechanical factor that regulates the kinetics of exocytosis. In agreement, inhibition of filopodia formation depressed SV fusion rates and increased asynchronous neurotransmitter release at high stimulation frequencies. Finally, blocking filopodia formation effectively reduced seizure severity in both *in vitro* and *in vivo* models of acute seizures, likely by regulating dynamic wiring and averting over-synchronization during excessive AP discharge. Thus, our study spotlights a novel synaptic plasticity mechanism and reveals its significant physiological and pathophysiological effects at synaptic and circuit levels.

### Influence of filopodia-generated membrane tension on SV exocytosis rates

The mechanical characteristics of the membrane, such as membrane tension, are unfolding as key regulators of membrane trafficking (*23*). Mathematical models and molecular dynamics simulations have provided proof of concept, demonstrating that increased membrane tension drives exocytosis by regulating fusion pore expansion and enhancing membrane merging rates (*22, 25, 26, 28*). Experimental evidence corroborates this, affirming that membrane tension, fuelled by actin dynamics, accelerates the fusion of vesicles with the plasma membrane (*27, 29, 49*). Membrane protrusions such as filopodia can increase membrane tension by pulling on the surrounding membrane and causing localized stretching. They also recruit molecules to their edges and exert contractile forces via the actin cytoskeleton that can be propagated over long distances (*18, 44*).

Our data support a model suggesting that the kinetics of SV exocytosis is heavily influenced by the mechanical properties of the release sites. Repeated SV fusion decreases local membrane tension, diminishing the force driving fusion. The emergence of filopodia counteracts this by raising membrane tension, thereby preventing a decline in SV exocytosis rates, and promoting synchronous neurotransmitter release. Prior research demonstrated rapid tension propagation along axons in developing neurons (*50*), but this process is slower in specialized presynaptic terminals, covering a few micrometers in a few seconds (*30*). Our data suggest that filopodia have a strong local effect on membrane tension and SV fusion rates. Although tension spreads more slowly from the terminal to the axon (*30*), synapses with filopodia may still maintain sufficient membrane tension to support SV fusion in nearby boutons during prolonged neuronal activity.

### Role of presynaptic filopodia in establishing dynamic synaptic connectivity. Implications for seizure-induced circuit remodelling

Filopodia are crucial in neuronal development (*51–59*). During synaptogenesis, the stochastic exploration of dendritic and axonal filopodia contributes to synaptic partner selection, with their stabilization possibly acting as a rate-limiting step for synaptogenesis (*60, 61*). Emerging evidence suggests that filopodia serve as a key reservoir for plasticity even in the adult brain (*62*).

Importantly, we demonstrate a direct link between presynaptic filopodia and neuronal activity in mature neurons and established neuronal circuits. Therefore, our study shifts the concept of synapse stability towards that of metastability. During synaptic transmission, the development of filopodia enables the presynapse to transiently extend from the axonal shaft and produce more widespread excitation and inhibition. Multi-synaptic boutons, where one synaptic bouton interfaces with multiple postsynaptic spines across different neurons, are common throughout the nervous system (*63–66*). Here we report a novel mode of synaptic connectivity underpinned by the transient contacts of the sprouting nerve terminal with multiple postsynaptic targets. We name this dynamic and temporary interaction between pre- and postsynaptic neurons a “neuronal kinapse”, drawing an analogy to the brief contacts between T-cells and antigen-presenting cells in the immune system (*67*). Furthermore, presynaptic filopodia displayed diverse dynamics, with approximately half disappearing during or shortly after stimulation, while the other half stabilized. These persistent filopodia could potentially establish stable synaptic connections or promote collateral axon branching, thereby contributing to adult morphogenesis. This concept provides a new framework for interpreting synaptic plasticity in the context of both physiological and pathological conditions.

Seizures trigger significant structural and functional changes in neuronal circuits. A prominent example is mossy fiber sprouting in the hippocampus (*13, 68*), which results in the formation of recurrent excitatory circuits that promote epileptogenesis (*69, 70*). Presynaptic filopodia formation is essentially a transient small-scale sprouting that can also contribute to the maladaptive remodelling of neuronal circuits. In support, inhibiting filopodia formation with EIPA resulted in increased asynchronous glutamate release, which in turn was associated with a decrease in ictal-like discharges in a 4-AP *in vitro* model of acute seizures, and a reduction of seizure severity *in vivo*. We propose that presynaptic filopodia can increase excitability through two complementary mechanisms. Firstly, they transiently increase synaptic connectivity.

Secondly, they modify the mechanical properties of the release sites, maintaining high SV fusion rates during excessive AP discharge. Both mechanisms promote a hypersynchronous state, which is assumed to underlie the abnormal coupling of hyperactive neuronal populations, leading to the onset of seizures (*71–73*), although the relationship is not linear (*48*). In contrast, desynchronization leads to the functional isolation of these populations and restricts epileptic activity (*74, 75*). An increase in asynchronous neurotransmitter release upon inhibition of filopodia can lead to desynchronization of neuronal circuit activity by lowering the signal-to-noise ratio of synaptic signals and distorting their temporal integration within the network. These results suggest that presynaptic filopodia can establish positive feedback loops within neuronal circuits. Their emergence promotes excessive excitation, triggering further filopodia growth, which ultimately results in hyperexcitability. This process could serve as one of the foundational explanations for the “seizures beget seizures” theory (*76*). Therefore, determining the precise mechanisms underlying seizure-induced filopodia formation is crucial, as it may provide new insights and therapeutic opportunities for intractable epilepsies.

In summary, these results significantly expand our current knowledge of synapse function and dysfunction. They highlight the limitations of relying solely on static maps of neuronal connectivity as an approach to understand how brain circuits are orchestrated and how network dysfunctions develop. Rather, they reveal synaptic connectivity as a dynamic feature, subject to rapid changes during AP discharge that have fundamental implications for synaptic physiology and pathophysiology.

## Acknowledgments

We thank Dr. Karen Smillie for assistance in grant administration. We are grateful to Nicole Lai and Suzanne Claxton for their excellent technical assistance in preparing mouse and rat hippocampal cultures, respectively. We also thank the CDBS IMPACT imaging facility, the Crick Advanced Light Microscopy facility (CALM) and the Edinburgh Super-Resolution Imaging Consortium (ESRIC) for their assistance with imaging and analysis. We extend our gratitude to Dr. Vincent Magloire and Dr. Kathrine Dobson for their input on imaging intact brain tissue. For the 3D-EM project design, sample preparation, transmission electron microscopy, analysis, and figure creation, we thank The Roslin Institute 3D Volume Electron Microscopy Capability within the Proteomics and Metabolomics Facility (funded by a BBSRC Core Capability Grant BB/CCG2270/1). We would like to thank Prof. Tara-Spires Jones, Prof. David Lyons (both, University of Edinburgh) and Prof. Gabriele Lignani (UCL) for their critical review and constructive feedback on this manuscript.

For the purpose of open access, the authors have applied a CC-BY public copyright license to any author-accepted manuscript version arising from this submission.

## Funding

EASTBIO BBSRC PhD studentship (AJ)

Walter Benjamin position, project number 458275811, of the DFG (JK)

Francis Crick Institute core funding from Cancer Research UK (CC2037); the UK Medical Research Council (CC2037), and the Wellcome Trust (CC2037) (SU)

Carnegie Research Incentive Grant RIG012496 (AGS) Royal Society Research Grant RGS\R1\221112 (AGS)

Reinhart Koselleck project, project number 399894546, of the DFG (CR)

Wellcome Trust Investigator Award 204954/Z/16/Z (MAC)

Simons Foundation 529508 (MAC)

Epilepsy Research Institute UK Endeavour project grant P2204 (DI and MAC)

FEBS short-term fellowship (DI)

## Author contributions

Conceptualization: DI

Methodology: AJ, JK, JW, JM, AGS, DI

Investigation: AJ, JK, JW, JM, DI

Visualization: AJ, JM, DI

Funding acquisition: SU, AGS, CR, MAC, DI

Project administration: DI

Supervision: AGS, CR, MAC, DI

Writing – original draft: DI

Writing – review & editing: all authors

## Data and materials availability

All data are available in the main text or the supplementary materials.

## Supplementary Materials

Materials and Methods

Figs. S1 to S7

Fiji code “Find responding synapses and set ROIs”

Tables S1 and S2

References (*77-86*)

## Materials and Methods

### Experimental design

Sample sizes were not determined in advance using power analysis because they were not chosen based on a predetermined effect size. Instead, we performed several independent experiments with biological replicates. Detailed descriptions of sample sizes and statistical analyses used to assess normality and calculate P values are provided in the “Statistical analysis” section, figures, figure legends, and tables S1 and S2. Data collection and analysis were carried out by multiple researchers, who were blind to the conditions whenever feasible. The experiment-specific inclusion criteria used for time-series analysis are outlined in their respective sections. No outliers were excluded from our analyses.

The objective of this research was to assess synaptic boutons for any structural alterations during synaptic transmission and elucidate their functional significance. We hypothesized that the structural reorganization of presynaptic terminals, induced by AP discharge, influences the mechanical properties of the plasma membrane, thereby regulating SV fusion and neurotransmitter release. We also hypothesized that these changes modulate network excitability, providing new insights into the mechanisms underlying epileptogenesis. All experiments were conducted within a controlled laboratory environment.

### Animals

Neuronal cells and brain tissue were obtained from Thy1-GFPM, wild-type C57BL/6J mice, and wild-type Sprague-Dawley rats of both sexes. C57BL/6J mice were sourced from Charles River Laboratories, while Thy1-GFPM mice were obtained from the Jackson Laboratory. Homozygous and hemizygous Thy1-GFPM show EGFP expression in sparse subsets of neurons (10% of cortical neurons). Both homo- and hemizygous animals were bred to maintain the mouse colony. Sprague-Dawley rats were sourced from an in-house colony at the Francis Crick Institute. Mouse colonies were housed in individually ventilated cages, whereas rat colonies were housed in standard open-top cages, all maintained on a 14-hour light/dark cycle (light from 07:00 to 21:00). Breeding animals were fed RM1 chow, and stock animals were maintained on RM3 chow. All animal procedures adhered to the U.K. Animal (Scientific Procedures) Act 1986, under Project and Personal License authorization, and received approval from the Animal Welfare and Ethical Review Body at either the University of Edinburgh (Home Office project licenses to M.A.C.: PP5745138 and A.G.S.: PP1538548) or the Francis Crick Institute (Home Office project license to S.K.U.: 70/7771). All euthanasia procedures were performed using schedule 1 protocols in accordance with U.K. Home Office Guidelines. Euthanasia of adult animals was performed through either cervical dislocation or exposure to rising CO_2_ levels followed by decapitation, while embryos were euthanized via decapitation followed by brain destruction. All procedures for the use and care of mice for electron microscopy experiments were authorized by the Animal Welfare Committee of Charité Medical University and the Berlin State Government.

### Materials

In this study, the following materials were used: dl-2-amino-5-phosphonopentanoic acid (AP-5; ab120271), 6-cyano-7-nitroquinoxaline-2,3-dione (CNQX; ab120044), 4-aminopyridine (4-AP; ab120122), amiloride hydrochloride (ab120281) and SMIFH2 (ab218296) from Abcam; bafilomycin A1 (A8627) from Stratech; papain (LK003176) from Worthington; EIPA (B7389) from ApexBio; dynasore (2897) from Tocris; tetanus toxin (TeTX) from Calbiochem (582243-25UG); Flipper-TR (251SC020) from Spirochrome; carbamazepine (18467) from Cayman Chemical; Neurobasal (21103-049), l-glutamine (25030-024), B27 supplement (17504-044), penicillin-streptomycin (15140-122), Fura Red, AM, cell permeant (F3021), Pluronic F-127 (P6867), Alexa Fluor 647 (10000 MW, Anionic, Fixable, D22914) and Lipofectamine 2000 (11668-019) from Thermo Fisher Scientific; fetal calf serum (S1810-500) from Biosera; FluorSave reagent (345789) from Merck Millipore; poly-d-lysine hydrobromide (P7886), d-laminin (L2020), 3,3′-diaminobenzidine (D8001), glutaraldehyde grade I (G5882), and protease inhibitor cocktail (P8849) from Sigma-Aldrich; osmium tetroxide, crystalline (E19132), and acetone (RT100016) from Electron Microscopy Services.

### Oligonucleotides

The following oligonucleotides were used to genotype Thy1-GFPM mice: ACA GAC ACA CAC CCA GGA CA (transgene forward); CGG TGG TGC AGA TGA ACT T (transgene reverse); CTA GGC CAC AGA ATT GAA AGA TCT (positive control forward); GTA GGT GGA AAT TCT AGC ATC ATC C (positive control reverse).

### DNA constructs

The following expression vectors were used in this study and were obtained from the indicated sources: sypHy, L. Lagnado (University of Sussex); syp-mOr2, S. Takamori (Doshisha University, Kyoto, Japan); red fluorescent protein (RFP)–Bassoon, C. Garner (German Centre for Neurodegenerative Diseases, Berlin); pCAG:GPI-GFP (Addgene plasmid # 32601 ; http://n2t.net/addgene:32601 ; RRID:Addgene_32601) (*77*); GPI-mApple (Addgene plasmid # 182868 ; http://n2t.net/addgene:182868 ; RRID:Addgene_182868) (*78*); mRuby-Lifeact-7 (Addgene plasmid # 54560 ; http://n2t.net/addgene:54560 ; RRID:Addgene_54560); SF-iGluSnFR.A184S (Addgene plasmid # 106174 ; http://n2t.net/addgene:106174 ; RRID:Addgene_106174)(*79*); GCaMP6f (Addgene plasmid #40755; RRID: Addgene_40755)(*80*); an shRNA targeting mouse NHE1 (mSlc9a1), target sequence CCACAATTTGACCAACTTAAT, was cloned into a lentiviral vector, pLV[shRNA]-EGFP-U6>mSlc9a1[shRNA#2], VB900142-8600taz, by VectorBuilder. The scrambled control plasmid (sequence GACTTTACTGCCCCTTACT) was from (*81*).

### Antibodies

The following primary mouse antibodies were used in this study: synaptobrevin 2 (WB, 1:2000; Synaptic Systems, 104211, RRID:AB_2619757); synaptotagmin 1 ATTO647N (ICC, 1:200; Synaptic Systems, 105311AT1, RRID:AB_993036); GluA Oyster550 (ICC, 1:200; Synaptic Systems, 182411C3, RRID:AB_2619877); The following rabbit antibodies were used: synaptophysin 1 (WB, 1:1000; Synaptic Systems, 101 002, RRID:AB_1961591); SV2A (ICC, 1:1000; Abcam, ab32942, RRID:AB_778192); vGAT Sulfocyanine 3 (vGAT-Cy3) (ICC, 1:100; Synaptic Systems, 131103C3, RRID:AB_2189809); NHE1 (ICC, 1:200, Thermo Fisher Scientific, PA520444, RRID:AB_11153959).

The following secondary antibodies were used in this study: donkey anti-rabbit Alexa Fluor 568 (ICC, 1:1000; Thermo Fisher Scientific, A10042, RRID:AB_2534017); goat anti-rabbit Alexa Fluor 488 (ICC, 1:1000; Thermo Fisher Scientific, A11008, RRID:AB_143165); goat anti-rabbit Alexa Fluor 647 (ICC, 1:1000; Thermo Fisher Scientific, A21244, RRID:AB_2535812); donkey anti-rabbit immunoglobulin G (IgG) IRDye 800 CW (WB, 1:10,000; LI-COR, 92532213, RRID:AB_2715510); goat anti-mouse IgG IRDye 800 CW (WB, 1:10,000; LI-COR, 92632210, RRID:AB_621842).

### Tissue culture and transfection

Primary dissociated hippocampal cultures were prepared from mouse (C56BL/6J, E17.5) and rat (E18.5) embryos. The mouse hippocampi were treated with papain (10U/ml) in Dulbecco’s phosphate-buffered saline (PBS) and then mechanically triturated into a single-cell suspension after rinsing with Minimal Essential Medium containing 10% fetal bovine serum (FBS). The rat hippocampal neurons were dissociated using trypsin and mechanically triturated. For both species, the cells were plated on 25 mm in diameter coverslips coated with poly-D-lysine and laminin.

The mouse neuronal cultures were grown in Neurobasal medium supplemented with 2% B-27, 0.5 mM L-glutamine, and antibiotics (penicillin and streptomycin at 1% v/v), with 1mM cytosine β-D-arabinofuranoside added at 3 days *in vitro* (DIV) to suppress glial growth. Rat neurons were initially cultured in modified Eagle’s medium supplemented with 10% FBS, 0.45% dextrose, sodium pyruvate (0.11 mg/ml), 2 mM glutamine, penicillin (100 U/ml), and streptomycin (100 mg/ml), and then switched to maintenance media containing 2% B27, 12.5 μM glutamate, and the same concentrations of penicillin, streptomycin, and glutamine. Media changes for rat neurons occurred every 4 days.

At DIV 7 and 8, the hippocampal neurons were transfected or co-transfected using Lipofectamine 2000 (Thermo Fisher Scientific, 11668-019), following manufacturer’s instructions. Imaging of these neurons occurred from DIV14 to 21.

For electron microscopy, primary hippocampal neurons from P0-P2 C57BL/6J mice of either sex were cultured on astrocyte feeding layers aged 1-3 weeks, which were grown on sapphire discs. To inhibit glial proliferation, 8 mM 5-fluoro-2-deoxyuridine and 20 mM uridine were added before neuron culturing. Neurons were maintained in Neurobasal-A medium supplemented with B-27 and Glutamax (Invitrogen, Germany) at 37°C for 15-21 days before analysis.

### Protein biochemistry

For quantifying protein expression via Western blotting (WB), cell cultures treated with 2nM TeTN or a vehicle for 24 hours were lysed in 2x SDS sample buffer. The buffer contained 67 mM tris (pH 7.4), 2 mM EGTA, 9.3% glycerol, 12% β-mercaptoethanol, bromophenol blue, and 67 mM SDS. The samples were boiled for 10 min before being separated using SDS-polyacrylamide gel electrophoresis (PAGE).

Proteins were resolved on SDS-PAGE and transferred to nitrocellulose membranes. The membranes were incubated with primary antibodies overnight at 4°C, followed by secondary antibodies for 1 h at room temperature. Imaging was performed using an Odyssey 9120 Infrared Imaging System (LI-COR Biosciences) with LI-COR Image Studio Lite software. Analysis was done using ImageJ (*82*), measuring the integrated density of signals within identical rectangular regions of interest (ROIs) set around the protein bands.

### Immunocytochemistry (ICC)

For ICC, neurons were fixed in 4% PFA in PBS for 4 to 5 min at room temperature. Samples were blocked and permeabilized with 10% FBS, 0.1% glycine, and 0.3% Triton X-100 in PBS for 20 min, then incubated with primary antibodies overnight at 4°C. Secondary antibodies were applied for 1 h at room temperature. Coverslips were mounted on microscope slides using FluorSave reagent (Merck Millipore). Immunofluorescence intensity was quantified in circular ROIs with a diameter of 1μm, placed over nerve terminals in ImageJ.

Live labelling with synaptotagmin1 (syt1-ATTO647N), GluA-Oyster550, or vGAT-Cy3 antibodies was performed by incubating neurons with fluorescently labelled antibodies dissolved in culture medium for 20-30 min at 37°C. For quantitative analyses, the same antibody solutions were applied to all coverslips in the corresponding experiments.

### Live imaging and analysis of SypHy and iGluSnFR reporters

All live imaging of primary neurons was performed at 37°C using an imaging buffer containing 136 mM NaCl, 2.5 mM KCl, 2 mM CaCl_2_, 1.3 mM MgCl_2_, 10 mM glucose, and 10 mM Hepes (pH 7.4). For electrical field stimulation, cells were mounted in an imaging chamber (Warner Instruments, USA, RC-21BRFS) equipped with parallel platinum wires spaced 0.6 cm apart. The neurons were subjected to several stimulation protocols, including 600AP (100mA and 1-ms pulse width) at 40Hz; 400APs at 40Hz; 600APs at 5Hz; 40APs at 20Hz; and 900APs at 20Hz. Imaging of SypHy and iGluSnFRA184S reporters was performed using an inverted Zeiss Axio Observer Z1 microscope, with an EC Plan-Neofluar 40x oil immersion objective (numerical aperture, NA 1.3), and a Colibri 7 light-emitting diode light source (Zeiss). The microscope was equipped with a Zeiss AxioCam 506 camera and was controlled by ZEISS ZEN 2 software. Transfected neurons were imaged using a GFP filter (exciter range 450 to 490nm; beam splitter at 495nm; emitter range 500 to 550nm), and images were acquired at 2Hz for SypHy and 2.5Hz for iGluSnFRA184S.

Quantitative analysis of SV fusion kinetics and SypHy response amplitudes was performed in the presence of 1 μM bafilomycin A1 (Stratech). Quantitative analysis of SV fusion kinetics upon reduction of membrane tension was performed during continuous perfusion of imaging buffer containing 350 mM sucrose and simultaneous stimulation with 900AP at 20Hz. Data from the SypHy time-lapse series were analysed using the Time Series Analyzer plugin for ImageJ (https://imagej.nih.gov/ij/plugins/time-series.html). ROIs were set at active SypHy synaptic terminals, and the average change in fluorescence intensity was tracked over time. The threshold for responsive terminals was set at an average fluorescence intensity increase of ≥2% above baseline upon AP stimulation. Any necessary adjustments for photobleaching were made using a correction factor obtained from inactive terminals on the same coverslip. The fluorescence response amplitudes elicited by field stimulation were normalized to the baseline (0) and the second response amplitude (1). Rate of exocytosis was measured from a linear fit. Statistical analyses were performed using Microsoft Excel and GraphPad Prism 10.

To enhance the detection of asynchronous glutamate release events, the neurons were transfected with iGluSnFRA184S, a high-affinity glutamate sensor displaying increased peak amplitudes, which enables better detection of glutamate release (*79*). Stimulation of sparsely transfected hippocampal neurons was performed with 40AP, 20Hz. The sensor’s slower-off rate, combined with the chosen image acquisition rate (2.5 Hz) enabled measurement of the collective synchronous glutamate response amplitude. However, individual release events in response to each AP were not detected.

To detect individual glutamate release sites on dendrites, we applied a Gaussian filter (Sigma, 2.0) in ImageJ to enhance the iGluSnFR.A184S signal and reduce background fluorescence. This preprocessing improved the accuracy of automatic release site detection. Glutamate release site positions were automatically identified using custom ImageJ macro employing the Find Maxima algorithm, which sets circular ROIs (∼1 x 1µm) centered on the detected maxima. The detection threshold was set to eliminate false-positive sites arising from background fluorescence. The synchronous glutamate response amplitude was measured and represented as ΔF/F_0_. Neuronal activity was monitored for 70 sec post-stimulation to capture potential asynchronous release events. Automatic detection of asynchronous events was performed using the Peak Analyzer tool in Origin (10.1.0.178), by applying the following detection criteria: the baseline was created using the XPS-Shirley model, peak identification was performed using the local maximum method, and peak filtering was based on a predefined threshold of peak height (20%). The frequency of asynchronous release events, expressed as the number of detected peaks per second, was plotted for analysis.

### Live imaging of filopodia formation and co-localisation analysis with SypmOr2, LifeAct, Bsn and GluA

High-resolution live imaging of synaptic membrane architecture during SV fusion was performed using a Nikon A1R confocal microscope, controlled by NIS Elements software. The microscope was equipped with a resonant scanner for high-speed imaging and a live cell imaging chamber with an environmental control unit, ensuring precise regulation of temperature and CO_2_ levels.

Primary hippocampal neurons were co-transfected with GPI-EGFP to label the membrane and SypmOr2 as a reporter for SV recycling, resulting in sparse labelling suitable for imaging individual synaptic boutons. To monitor actin dynamics, neurons were co-transfected with GPI-EGFP and LifeAct-Ruby. For co-localisation analysis with Bsn, neurons were co-transfected with GPI-EGFP and Bsn-RFP. For co-localization analysis with postsynaptic structures and inhibitory presynapses, GPI-EGFP-transfected hippocampal neurons were live-labelled with either GluA-Oyster550 or vGAT-Cy3 antibodies for 20-30 min before imaging.

Imaging was performed at 35-37°C. Z-stacks with a Z-step of 0.125 µm were acquired every 3 to 4 sec for 1.5 or 3.5 min. Typically, imaging began with a baseline period, followed by either 15 sec (600AP, 40Hz) or 120 sec (600AP, 5Hz) of stimulation, and then an additional 1 minute of imaging post stimulation. To assess the calcium and activity dependence of filopodia formation, neurons were imaged before, during and after a short pulse of 500mM sucrose (5 sec).

A 60x Plan Apo VC/NA 1.4 oil objective was used for imaging. Confocal settings were configured with a display resolution of 512x512 pixels, a 12-bit dynamic range, and an optical zoom of 8.0, resulting in a final pixel size of 0.05μm. Excitation of EGFP was achieved at 488nm, while emission light was collected at 525/50nm. Excitation for SypmOr2, LifeAct-Ruby, and GluA-Oyster550 was performed at 561nm, with fluorescence signal detection at 595/50nm. All time-lapse series of image stacks were deconvolved with Huygens Professional version 23.04 (Scientific Volume Imaging, The Netherlands, http://svi.nl), using the CMLE algorithm, with SNR:15, 100 iterations, and automatically estimated background value.

Imaging of filopodia formation during SV fusion triggered by short stimulation (40AP, 20Hz) was performed on an inverted Nikon Ti2 microscope. This microscope was equipped with Photometrics Kinetix back-illuminated sCMOS camera and “Nikon D-LEDI” 4 LED Illumination and controlled by NIS Elements software. The integrated live cell imaging chamber and a control unit allowed maintenance of a constant 37°C environment.

Primary hippocampal neurons were co-transfected with GPI-EGFP for membrane labelling and SypmOr2 to report SV recycling. Imaging duration was 1.5 minutes, including a baseline measurement, a 2-second stimulation period (40AP, 20Hz), and approximately 75 seconds of post-stimulation observation. A 60x Plan Apo /NA 1.42 oil objective was used for imaging.

Images were acquired at a frequency of 0.5Hz with a 12-bit depth. EGFP was excited at 475 nm and its emission detected at 520/30 nm, while SypmOr2 excitation occurred at 550 nm with emission detection at 641/75 nm. Automatic 2D deconvolution was performed on all time-lapse series using the built-in wizard in NIS Elements software.

The morphometric analysis of filopodia was performed on maximum intensity Z-projections obtained from time-lapse series of image stacks, with the exception of the dynamic 3D colocalization analysis of GluA and filopodia. To sharpen the cell boundaries, an unsharp mask filter (with a radius sigma of 1 pixel and a mask weight of 0.6) was applied in ImageJ, before segmentation using the Otsu thresholding algorithm.

Morphological changes over time were extracted using the Image Calculator tool in ImageJ. This involved subtracting the binary image of a pre-stimulation time point from subsequent time points to highlight regions of change and generate filopodial masks. The time point at which filopodia reached their maximum extension length was identified and quantitative data on morphological features such as ferret length and roundness were computed.

To analyze the ratio of transient vs stable filopodia, the following criteria were established. For filopodia induced by electrical field stimulation, transient filopodia were defined as those that emerged and retracted either during the stimulation period or within 15 sec post-stimulation. For filopodia induced by 4-AP treatment (both in primary cultures and acute slices), transient filopodia were identified as those that persisted for no more than 40 sec following their initial emergence.

For the co-localization analysis with synaptic markers, the segmentation of presynaptic filopodia at their maximum length was performed as described above, with the segmented structures then being converted into binary masks. These masks were subsequently applied to the fluorescence channel associated with synaptic markers to evaluate their co-localisation with filopodia.

Following background subtraction, the fluorescence intensity of each synaptic marker (Bsn, vGAT, GluA, and LifeAct) was quantified within the filopodial masks. Co-localisation was determined after binarization, using predefined intensity thresholds. The presence of synaptic markers at the synapses from which the filopodia branched was also assessed. This was done by measuring the marker intensity within a circular ROI (∼ 1 x 1 µm) centered on the axonal shaft from which the filopodia extended. The intensity threshold was selected to discriminate true signals from background noise, thereby reducing the incidence of false-positive results.

DiAna ImageJ plugin was used to perform dynamic 3D co-localisation analysis of GluA and GPI-EGFP-labelled axonal segments during filopodia formation (*83*). Rectangular ROIs measuring 6.1x6.1µm were cropped and centered around axonal segments where filopodia were observed. On deconvolved image stacks, 3D segmentation was performed using DiAna’s built-in segmentation algorithm with a uniform global intensity threshold applied to the pre-stimulation image stack (a single time point) and the stack at maximum filopodia length. This segmentation allowed assessment of the 3D spatial volume occupied by GluA and GPI-EGFP-labelled objects. The computation of the number of co-localised GluA-labelled objects and GPI-EGFP, along with the calculation of co-localised voxels between the two, was achieved through post-thresholding analysis of the respective labelled images.

The examples of 3D time-lapse series in Fig. 3G were generated using the shadow projection function in Imaris 10.1.0.

### Imaging and analysis of the coupling between filopodia formation and axonal calcium spikes

Imaging spontaneous filopodia formation and axonal calcium dynamics was performed with Nikon Ti2 microscope using the same settings as described above. Primary hippocampal neurons were co-transfected with GPI-Apple to label the membrane and GCaMP6f as a calcium sensor. Prior to imaging, neurons were treated with either 100 µM 4-AP or a combination of 50 µM APV and 10 µM CNQX and incubated for 30 min. Time-lapse imaging was performed for 5 min (0.5 Hz) in the presence of these inhibitors.

Analysis was performed in ImageJ. Circular ROIs (∼1 x 1µm) were set around putative synapses showing filopodia formation. Within these ROIs calcium transients, reported by GCaMP6f fluorescence, were monitored over the 5 min period, concurrently with filopodia dynamics. To optimize the signal-to-noise ratio and facilitate peak identification, a pre-processing step was performed on the obtained calcium traces using second-order smoothing in GraphPad Prism 10. Positive peaks in the data traces were detected automatically using the Peak Analyzer tool in Origin (10.1.0.178). The following criteria were used for the analysis: the baseline was established using median constant baseline, local maximum method was used for peak finding, and peak filtering was executed based on a predefined threshold of peak height (20%). The temporal dynamics of calcium transients were quantitatively characterized by extracting peak positions, which served as a measure for assessing the timing of the calcium events. These data were juxtaposed with the corresponding times of filopodia extension branching from the same ROIs (maximum length). The resulting graphical representation, (Fig. S1, D and F), displays the temporal coupling between calcium events and filopodial extension dynamics.

### FLIM imaging

Synaptic boutons in primary hippocampal cultures were live labelled with syt1-ATTO647N antibody for 20-30 min prior to incubation with either 50 µM EIPA or a vehicle for a period of 30 min. Within the final 15 min of the drug incubation, 1.5 µM Flipper-TR was introduced. Imaging was performed in an imaging buffer supplemented with 10 μM CNQX and 50 μM APV.

FLIM of Flipper-TR and intensity imaging of syt1-ATTO647N were performed using a Leica SP8 FALCON system, a fully integrated FLIM confocal platform controlled by LAS X 3.5.2.18963 software, equipped with a pulsed White Light Laser and HyD single molecule detection detectors (HyD SMD). Imaging was performed using an HC PL APO CS2 63x/NA 1.40 oil objective.

Confocal settings for image acquisition were configured as follows: a display resolution of 512 by 512 pixels, an 8-bit dynamic range, an optical zoom of 3.0, resulting in a final pixel size of 0.12 μm. A three-time frame averaging was applied to frames, with a scan speed of 400 Hz. Excitation of ATTO647N was achieved at 647 nm, while emission light was collected within the range of 681 to 771 nm. Flipper-TR excitation was performed at 480 nm, with fluorescence signal detection occurring in the range of 500 to 616 nm via HyD SMD.

An area of interest containing multiple syt1-ATTO647N-labelled synaptic terminals was selected, and a single FLIM image was acquired before field stimulation (or before 500 mM sucrose pulse). A train of 600AP, 40Hz was triggered offline. A second FLIM image of the same area of interest was captured during the last 5 sec of stimulation (or during the 5 sec of 500 mM sucrose application). Fluorescence decay data, obtained from full images, were then fitted to a double-exponential model using the LAS X software. Quality assessment of the fit was established based on the proximity of the χ2 to 1. For assessment of membrane tension changes at synapses, ROIs were set around syt1-ATTO647N labelled synaptic boutons using the LAS X software and applied to the FLIM Flipper-TR image. The same ROIs were applied to the FLIM images acquired at rest and during field stimulation. The relative change in membrane tension between rest and stimulation (or before and during sucrose application) was computed as: Δtau = Flipper-TRtau_rest (before sucrose) − Flipper-TRtau _stimulation (+sucrose). The pseudo-colour FLIM images (Fig. 4, A and B) were produced using LAS X, with the Flipper-TRtau display range specified in the figures.

### Acute slice preparation, Fura Red AM dye loading and live imaging of EGFP-expressing presynaptic boutons in coronal brain sections from Thy1-GFPM mice

Coronal brain slices (350 μm) were prepared from Thy1-GFPM mice (8-10 weeks old of either sex). The brains were quickly transferred to chilled (2° to 5°C) carbogenated sucrose-modified artificial cerebrospinal fluid (saCSF; 86 mM NaCl, 1.2 mM sodium phosphate buffer, 2.5 mM KCl, 25 mM NaHCO_3_, 25 mM glucose, 50 mM sucrose, 0.5 mM CaCl_2_, and 7 mM MgCl_2_) for 2 min. Brains were subsequently sectioned in the same solution using a vibrating microtome (Leica, VT1200S). The brain sections were left to recover for 30 min at 37°C in carbongenated standard aCSF, containing 126 mM NaCl, 3 mM KCl, 1.2 mM sodium phosphate buffer, 25 mM NaHCO_3_, 15 mM glucose, 2 mM CaCl_2_, and 2 mM MgCl_2_.

Neuron loading with Fura Red AM was performed as previously described (*84*). A fresh stock solution of 20% Pluronic F-127 in DMSO was prepared using sonication. The calcium loading solution was prepared by mixing Fura Red AM (30 µM) and Pluronic F-127 (0.065%) in carbongenated standard aCSF. The brain slices were covered with the calcium loading solution and incubated at 37°C/5% CO₂ for 45-60 min in a cell culture incubator. After incubation, the dye solution was replaced with fresh aCSF to prevent dye overload. The slices were then incubated with either 100 µM 4-AP or a combination of 50 µM APV and 10 µM CNQX for additional 30 min prior to live imaging for 5 minutes in the presence of the inhibitors.

Brain slices loaded with Fura Red AM were placed in an imaging chamber mounted on a Nikon A1R confocal microscope. Superficial cortical layers with sparsely EGFP-labelled axons and putative synaptic boutons were selected for imaging. Time-lapse imaging was performed at 35- 37°C with 5% CO₂, using a resonant scanner over a 5 min period. A 40x Plan Fluor/NA 1.3 oil objective was used for imaging. Image stacks (typically, 50-70 sections, spaced 0.125µm apart) were acquired at a speed of ∼0.07Hz. Confocal settings were adjusted to a 1024 x 1024-pixel resolution, 12-bit dynamic range, and 4.0 optical zoom, achieving a final pixel size of 0.04 μm. Dual excitation of EGFP and Fura Red at 488 nm was implemented, with emissions collected at 525/50 nm for EGFP and 595/50 nm for Fura Red. The interaction of Fura Red with cytoplasmic calcium reduced its excitation efficiency at 488 nm, decreasing its emission at ∼650 nm.

All time-lapse series of image stacks were deconvolved with Huygens Professional version 23.04 (Scientific Volume Imaging, The Netherlands, http://svi.nl). A Gaussian filter with (sigma: 4.0) was applied to the Fura Red channel in ImageJ to enhance the signal and minimize background noise. Synaptic boutons marked by Fura Red were inspected for fluorescence decreases, serving as a proxy for calcium dynamics through kymographs in ImageJ. The data used to generate kymographs were scaled multiple times to enhance visual clarity.

### High-pressure freezing and freeze substitution

High-pressure freezing (HPF) and freeze substitution (FS) were performed as described in (*85*). For HPF, neurons (DIV15-21) grown on 6 mm carbon-coated sapphire glass disks were treated with specified compounds (DMSO as a vehicle, 50 µM EIPA, 1 mM amiloride, and 30 µM SMIH2 for 30 min, and 80 µM dynasore for 3-5 min) in an extracellular solution containing 140 mM NaCl, 2.4 mM KCl, 10 mM HEPES, 2 mM CaCl_2_, 4 mM MgCl_2_, and 10 mM glucose (pH adjusted to 7.3 with NaOH, 300 mOsm) in an incubator at 37°C. For cryofixation, sandwiches consisting of the cultured sapphire glass disk, a 100 nm spacer ring, another sapphire glass disk, and a 400 nm spacer ring, and filled up with extracellular solution and the respective compound, were assembled. They were electrically stimulated with 600AP, 40Hz at 37°C and cryofixed within 5 ms following stimulation using a Leica ICE high-pressure freezer.

The sapphire disks were then placed in cryovials with a freeze substitution solution consisting of 1% (w/v) osmium tetroxide, 1% (v/v) glutaraldehyde, and 1% ddH_2_O in anhydrous acetone, pre-cooled to −90°C. FS was performed using a Leica Microsystems AFS2 device with the following temperature protocol: −90°C for 5 h, ramping from −90°C to −20°C over 14 h (5°C/h), holding at −20°C for 12 h, and then ramping from −20°C to 20°C over 4 h (10°C/h).

The samples were then washed four times for 15 min each with anhydrous acetone, stained with 0.1% (w/v) uranyl acetate in acetone for 1 h, and washed again four times for 15 min each with anhydrous acetone, all at room temperature. Next, the samples were infiltrated with epoxy resin (42.8% (w/w) Epon, 31.2% (w/w) DDSA, 26% (w/w) MNA; Fluka) by soaking for 2 h in 30% Epon/acetone, 2 h in 70% Epon/acetone, and overnight in 90% Epon/acetone. After a day, the sapphire disks were placed in embedding molds with a mixture of Epon and Araldite (26.3% (w/w) Epon, 18.7% (w/w) Araldite, 51.9% (w/w) DDSA, 2.98% (w/w) BDMA; EMS) for polymerization at 60°C for 48 h.

### Transmission electron microscopy and analysis of 2D and 3D electron microscopy

For 2D EM, ultrathin sections (50 nm thick) were cut using an Ultracut UCT ultramicrotome (Leica) and collected on formvar-coated 200-mesh nickel or copper grids. Prior to imaging, the samples were post-stained with 2% (w/v) uranyl acetate in ddH_2_O and Reynold’s lead citrate for enhanced contrast. The sections were then imaged using a JEOL JEM 1400plus transmission electron microscope equipped with a OneView camera. Images were captured at 60 kV with a pixel size 0.28x0.28 nm. All imaging was performed blind to the experimental conditions to prevent bias.

Images from all groups per culture were pooled and randomized into stacks using a custom ImageJ macro. Between 70 and 100 images were acquired per group.

Structural features were labelled using line and selection tools in ImageJ. Data analysis was performed using custom ImageJ macros and Python scripts (*85*). The AZ was defined as the presynaptic membrane facing PSD. Docked SVs were those whose outer membrane contacted the AZ membrane. The distribution of cytosolic SVs was measured by calculating the shortest distance to the AZ membrane, and SV diameters were determined by fitting a circle to the outer membrane. Endosome-like structures were identified as spherical (diameter >60 nm) or tubular structures with a single outer membrane and a clear lumen.

For 3D EM, brightfield images were captured of stimulated (600AP, 40Hz) HPF/FS samples and candidate regions of fine axons selected for TEM (Zeiss AxioImager Z2). Samples were then glued to a resin block, trimmed with blades and glass knives, and ribbons of serial 70 nm-thick ultrathin sections cut (Leica Ultracut 7 ultramicrotome) and collected onto formvar/carbon-coated single slot copper grids. Grids were contrasted with Uranyless and 3% lead citrate before imaging. TEM images were acquired using a 12-sample carousel and annular scanning TEM (aSTEM) detector housed inside a Zeiss Gemini 360 field emission scanning electron microscope. Serial images from 3 samples were used to find 11 filopodia on 5 axons and then to trace filopodia (split into 200 nm segments from the tip), parent axons, endosomes, vesicles, mitochondria and filaments using ImageJ (TrakEM2 plugin). These tracings were used to create 2D images, 3D reconstructions and to measure segment volumes of filopodia and endosome volumes to calculate endosome fractions (% of segment volume). SVs were counted for each segment then expressed as a density of SVs per µm^3^ volume of each segment.

The raw EM data presented in Fig. 5D was obtained from the publicly available EM dataset: H01 (*46*). The Neuroglancer platform was used for visual exploration of the dataset, the EM ultrastructure and segmentations. For 3D visualization (right in Fig.5D), only axons containing synaptic boutons with putative filopodia, and their postsynaptic targets were rendered. Serial EM images from the same sample were analyzed to trace filopodia and synaptic organelles (SVs, endosomes, mitochondria, and endoplasmic reticulum), using the ImageJ TrakEM2 plugin, as described above. These tracings were then used to generate the 2D images and 3D reconstructions (middle) displayed in Fig. 5D.

### Acute brain slice preparation for *in vitro* electrophysiology

4–8-week-old C57BL/6 mice were used for all *in vitro* electrophysiology experiments. Mice were humanely euthanized via cervical dislocation with death confirmed by decapitation. Brains were rapidly dissected and transferred to ice cold sucrose-based aCSF bubbled with carbogen (95% O_2_, 5% CO_2_). This consisted of (in mM) sucrose 75, NaCl 87, KCl 2.5, NaHCO_3_ 25, NaH_2_PO_4_ 1.25, Glucose 25, MgCl_2_ 7, CaCl_2_ 0.5. Acute horizontal brain sections were then cut at 350µm on a Leica VT1200s vibrating microtome. Slices were then allowed to recover at 37°C in a chamber filled with sucrose-based aCSF bubbled with carbogen for at least 30 minutes, before being transferred to a chamber containing aCSF consisting of (in mM) NaCl 125, KCl 2.5, NaHCO_3_ 25, NaH_2_PO_4_ 1.25, Glucose 25, MgCl_2_ 1, CaCl_2_ at room temperature where they maintained until required for experimentation.

### Electrophysiological recordings and data analysis

Prior to recording, slices were preincubated for 30 min with 0.2% DMSO (used as a vehicle); or either 50 µM EIPA or 80 µM carbamazepine dissolved in 0.2% DMSO in aCSF. Slices were then transferred to a recording chamber and continually perfused with carbogenated aCSF at a flow rate of 2-3 min.

*In vitro* seizure experiments were recorded as extracellular field recordings. For these experiments, a broken glass micropipette (resistance 0.5-2 MΩ) filled with aCSF was implanted into the CA1 pyramidal layer. All recordings were acquired in current clamp. Following a baseline recording period, a 45 min free-recording period was commenced in which 100 µM 4-AP dissolved in aCSF was washed in via the perfusion system to induce seizure-like events within the slice.

For whole-cell experiments, a bipolar stimulating electrode made from twisted nickel-chromium wire (80/20 ratio; 0.066 mm width) was implanted into stratum radiatum at the boundary between CA1 and CA2. A glass micropipette (3-6 MΩ) was then filled with an internal recording solution consisting of (in mM) K-gluconate 142, KCl 4, EGTA 0.5, HEPES 10, MgCl_2_ 2,

Na_2_ATP 2, Na_2_GTP 0.3, Na_2_-phosphocreatine 1 and biocytin 2.7. Whole-cell configuration was achieved within a CA1 pyramidal neuron by use of negative pressure to rupture the cell membrane. Synaptic properties were tested in voltage clamp as previously described (*86*).

Briefly, the amplitude of excitatory postsynaptic currents (EPSCs) was first tested in response to a series of single 50 µs stimulation pulses (100 – 10000 µA). The stimulation intensity was then set to give an EPSC of approximate 200 pA. Presynaptic properties were then examined through a tetanic stimulation protocol of 600 pulses at 40 Hz. This protocol was used to assess the plasticity of synchronous neurotransmitter release during repeated stimulation.

All recordings were sampled at 20 kHz and acquired using Clampex v10.7 with an Axon Instruments Multiclamp 700B amplifier and Digidata 1550B digitizer.

All data were analysed offline using Axograph X v1.8.0 or Clampfit v10.7. For synaptic properties, peak EPSC amplitude was measured for each stimulation pulse and normalised to the first pulse in the train. Asynchronous release frequency values were calculated by utilizing the template matching feature in Clampfit to detect all spontaneous excitatory currents in the first 0.8 s after the termination of the 40Hz train.

For *in vitro* seizure experiments, a bandpass filter (lowpass 2 Hz; highpass 2 kHz) was first applied offline to raw traces in Clampfit. Individual seizure-like events were then detected using a custom template taken from an average of multiple events within an example control recording. To calculate metrics from these experiments, a representative single event template was generated by averaging multiple events from a control DMSO recording. Events were classed as a single continuous ictal period where the interevent interval between two consecutive events was less than 5 s. Total ictal period was calculated as the summation of the duration all ictal periods, and interictal period encompassed the duration within the recording not classed as an ictal period following the onset of the first detected event. Ictal burst frequency was calculated as the mean event frequency of all events within the ictal period in each recording.

### Behavioral assessment of 4-AP-induced acute seizures

Female C57BL/6J mice, 8-10 weeks old, were used for the experiments. 4-AP was dissolved in 0.9% saline and administered subcutaneously at 6.65mg/kg. To assess the protective effects of EIPA (5mg/kg) and carbamazepine (20mg/kg) against the behavioral effects induced by 4-AP, these drugs were injected intraperitoneally 30 min before 4-AP. DMSO served as a vehicle control. All drugs were prepared in 0.9% saline. Mice were placed in a testing arena, and their behavioral responses were monitored for 30 minutes post 4-AP injection with both top and side video cameras. Seizure intensity was evaluated using a modified Racine scale: 1. sudden behavioral arrest; 2. head nodding and facial jerking; 3. clonus of one forelimb (sitting); 4. clonus of both forelimbs (sitting); 5. clonic, tonic-clonic seizure (lying on belly); 6. clonic, tonic-clonic seizure with wild running and jumping; 7. tonic extensions, possibly resulting in respiratory arrest and death. The total seizure score was determined by adding together the scores from every two-minute interval. Behavioral evaluations were performed blindly to experimental conditions.

### Statistical analysis

Statistical analysis was performed using GraphPad Prism 10. To determine the appropriate statistical tests, the distribution of the datasets was assessed using the D’Agostino-Pearson normality test. The points in bar graphs represent the average value from each individual experiment. For datasets that followed a normal distribution, results are expressed as means or means ± SEM. For non-normally distributed data, results are given as median ± interquartile range. Comparisons of non-Gaussian datasets were made using Mann-Whitney (two-tailed), Wilcoxon matched-pairs signed-rank (two-tailed), and Kruskal-Wallis tests with Dunn’s post hoc analysis. For normally distributed datasets, comparisons were made using Student’s t test (two-tailed) and analyses of variance (ANOVAs) followed by Dunnett’s or Tukey’s post hoc tests.

Two-way ANOVA was used to assess the effects and interactions in multigroup comparisons when applicable. Details on sample sizes, the statistical tests used to compute *p* values, and the numerical results are provided in the figures, figure legends, and Tables S1 and S2.

---

**Fig. S1.**
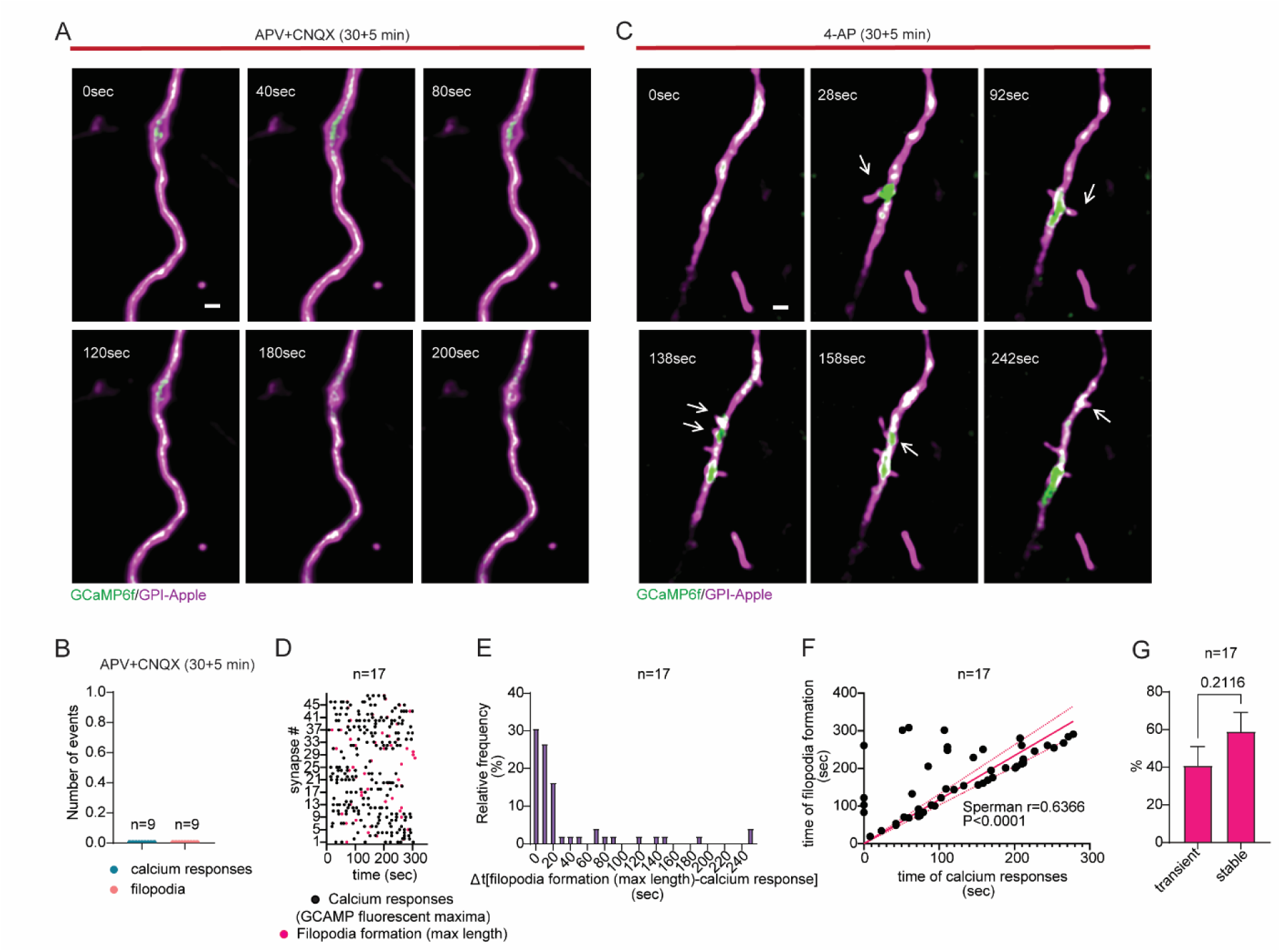
Temporal coupling of axonal calcium responses and filopodia formation. Primary hippocampal neurons were co-transfected with GPI-Apple and GCaMP6f, then treated with 50 µM APV and 10 µM CNQX (**A,B**) or 100 µM 4-AP for 30 min (**C-G**). Live imaging was performed for 5 min with inhibitors present. (**A**) Time-lapse series of APV+CNQX treated neuron showing no GCaMP6f fluorescence responses or filopodia formation. Scale bar 1 µm. (**B**) Number of GCaMP6f responses (blue) and filopodia formed (pink) in APV+CNQX treated cultures. (**C**) Time-lapse series showing spatio-temporal coupling of filopodia formation and axonal calcium responses (white arrows). Scale bar 1 µm. (**D**) Time histogram of simultaneous GCaMP6f responses and filopodia formation. Each row represents a synaptic bouton; symbols indicate GCaMP6f peak responses and filopodia reaching maximum length. (**E**) Frequency histogram of time differences between GCaMP6f peak responses and filopodia formation. (**F**) Correlation between GCaMP6f peak responses and filopodia formation with linear regression (pink line) and 95% confidence bands (dotted lines). (**E, F**) GCaMP6f peak fluorescence responses occurring immediately before filopodia formation were included in the analysis. (**G**) Ratio of stable vs transient filopodia. p values, unpaired t-test.

**Fig. S2.**
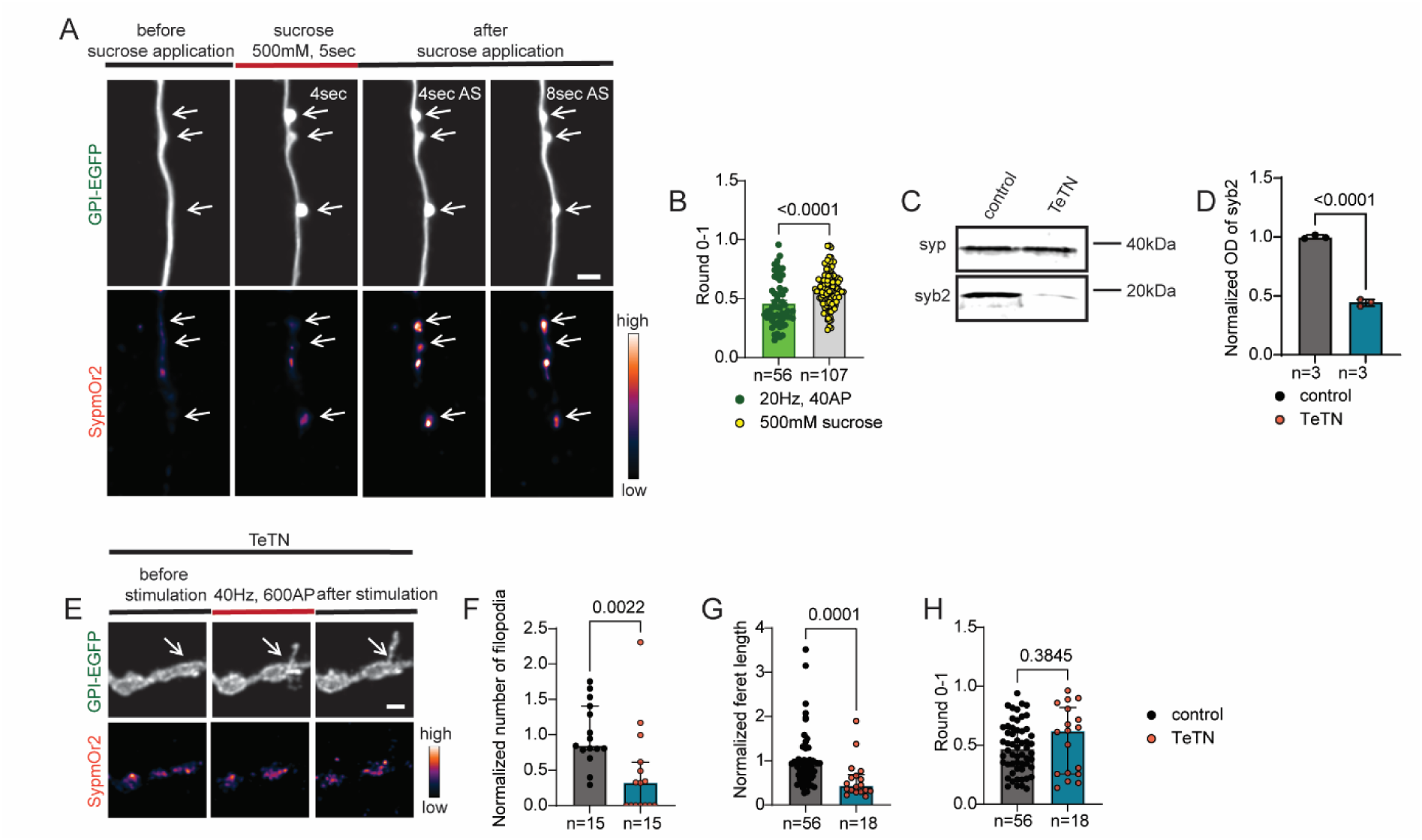
SV fusion is not an absolute requirement for activity-induced filopodia formation. (**A**) Primary hippocampal neurons were co-transfected with GPI-EGFP and SypmOr2. Neurons received a 5 sec hypertonic sucrose pulse (500 mM). Time-lapse series shows sucrose-triggered SypmOr2 fluorescence responses and structural changes at synapses (white arrows indicate presynaptic enlargements and SypmOr2 responses). Scale bar 1 µm. (**B**) Quantification of presynaptic structural alterations (roundness) after 40AP at 20Hz or sucrose pulse, both mobilizing RRP. 20Hz, 40AP, the same values as in Fig. 1I. p values, unpaired t-test. (**C**) Neurons treated with 2 nM tetanus toxin (TeTN) or vehicle control for 24 h, then lysed. Representative western blots (WBs) with synaptobrevin2 (syb2) and synaptophysin (syp) antibodies. (**D**) WB analysis of syb2 expression levels normalized to syp. p values, unpaired t-test. (**E**) Neurons treated with 2 nM TeTN for 24 h. Time-lapse series shows filopodia formation despite lack of SypmOr2 responses (white arrows indicate filopodia). Scale bar 1 µm. (**F**) Normalized number of filopodia per 10 µm axon length in cultures with and without 2 nM TeTN for 24 h (stimulation: 600AP, 40Hz). (**G**) Normalized filopodia length in cultures with and without 2 nM TeTN for 24 h (stimulation: 600AP, 40Hz). (**H**) Quantification of filopodia shape (roundness) in cultures with and without 2 nM TeTN for 24 h (stimulation: 600AP, 40Hz). (**F-H**) p values, Mann-Whitney U test.

**Fig. S3.**
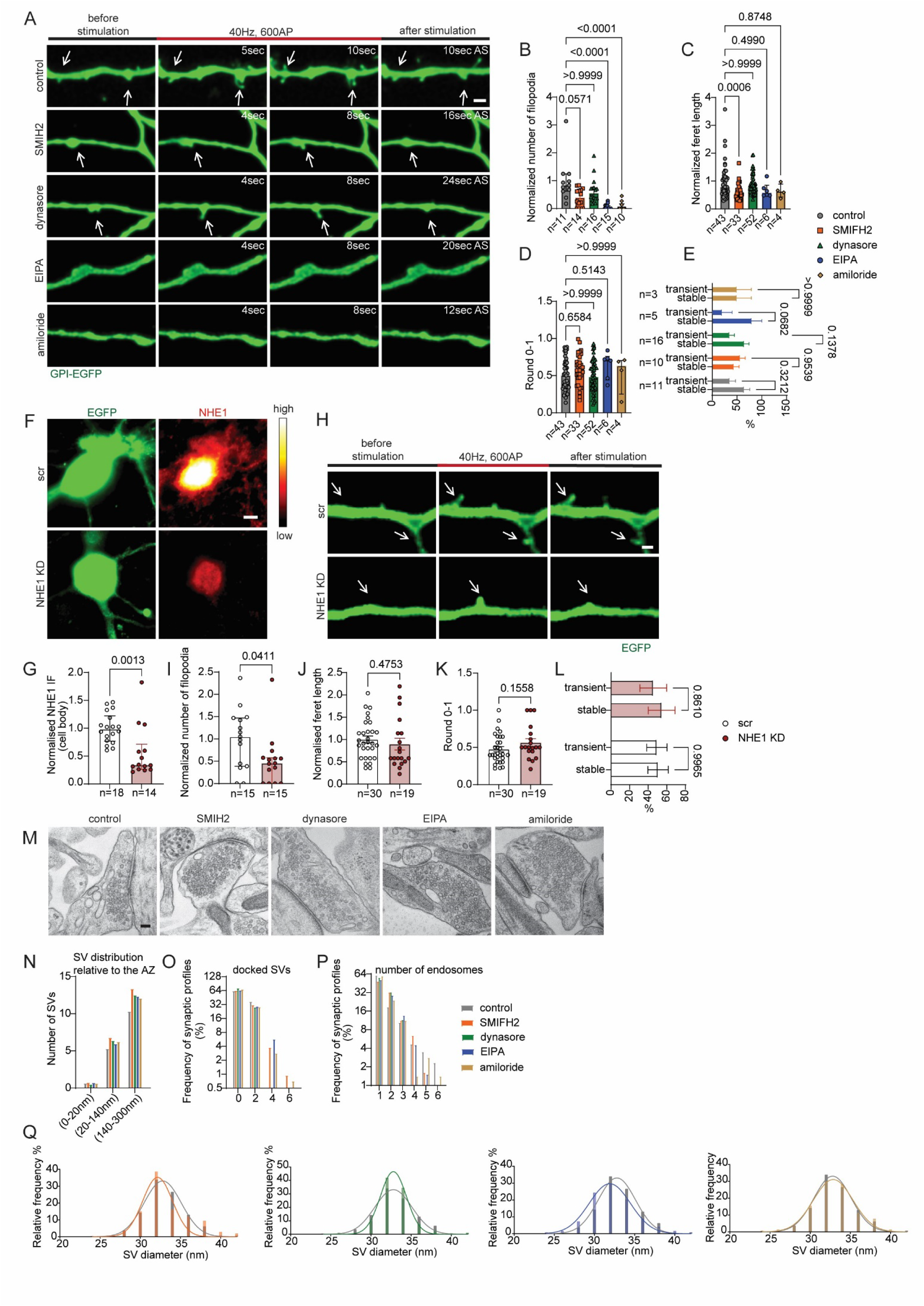
Membrane ruffling inhibitors EIPA and amiloride eliminate activity-induced filopodia formation at the presynapse. (**A-E**) Hippocampal neurons transfected with GPI-EGFP were treated for 30 min with vehicle, 30 µM SMIH2, 50 µM EIPA, 1 mM amiloride, or acutely with 80 µM dynasore for 2-4 min. Neurons were stimulated with 600APs (40Hz) and filopodia formation was monitored using time-lapse imaging. (**A**) Confocal time-lapse series showing filopodia formation (white arrows) with and without inhibitors. Scale bar 1 µm. (**B**) Normalized number of filopodia per 10 µm axon length. p values, Kruskal-Wallis with Dunn’s test. (**C**) Normalized filopodia length. p values, Kruskal-Wallis with Dunn’s test. (**D**) Quantification of filopodia shape. p values, Kruskal-Wallis with Dunn’s test. (**E**) Ratio of stable to transient filopodia. p values, two-way ANOVA with Sidak’s test. (**F-L**) Neurons transfected with shRNA targeting NHE1 or scrambled control, with EGFP as a reporter. (**F**) Images showing EGFP and NHE1 expression in neurons transfected with scrambled and NHE1KD plasmids. Scale bar 5 µm. (**G**) Quantification of NHE1 expression in cell bodies. p values, Mann-Whitney U test. (**H**) Time-lapse series showing filopodia formation in neurons transfected with scrambled or NHE1KD plasmids (white arrows). Scale bar 1 µm. (**I**) Normalized number of filopodia per 10 µm axon length. p values, Mann-Whitney U test. (**J**) Normalized filopodia length. p values, unpaired t-test. (**K**) Quantification of filopodia shape. p values, unpaired t-test. (**L**) Ratio of stable to transient filopodia. p values, two-way ANOVA with Sidak’s test. (**M**) Electron microscopy images of synaptic boutons treated with 30 µM SMIH2, 50 µM EIPA, 80 µM dynasore, and 1 mM amiloride, stimulated with 600AP at 40Hz. Scale bar 100 nm. (**N**) Number of SVs in different regions from the AZ. (**O**) Percentage of synaptic profiles with different numbers of docked SVs. (**P**) Percentage of synaptic profiles with different numbers of endosomes. (**Q**) Distribution of SV sizes in different treatment groups, with control group plotted alongside each treatment.

**Fig. S4.**
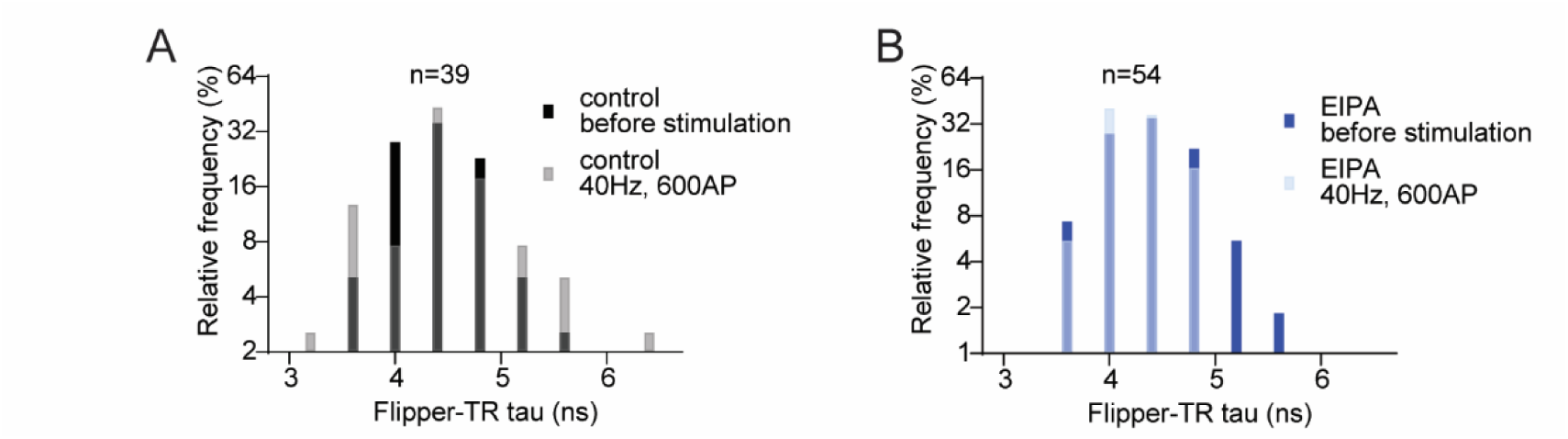
Distributions of Flipper-TR tau in control and EIPA treated neurons before and during stimulation. (**A,B**) Synaptic boutons in primary hippocampal cultures were live labelled with an ATTO 647N-conjugated antibody against the luminal domain of synaptotagmin1. Neurons were incubated with Flipper-TR and its fluorescence lifetime (tau) was monitored using FLIM imaging. (**A**) A frequency histogram showing the distribution of Flipper-TR tau in control neurons before and during stimulation with 600APs (40Hz). Note the widening of the histogram of Flipper-TR tau during stimulation. (**B**) A frequency histogram showing the distribution of Flipper-TR tau in EIPA (50 µM, 30 min) treated neurons before and during stimulation with 600APs (40Hz).

**Fig. S5.**
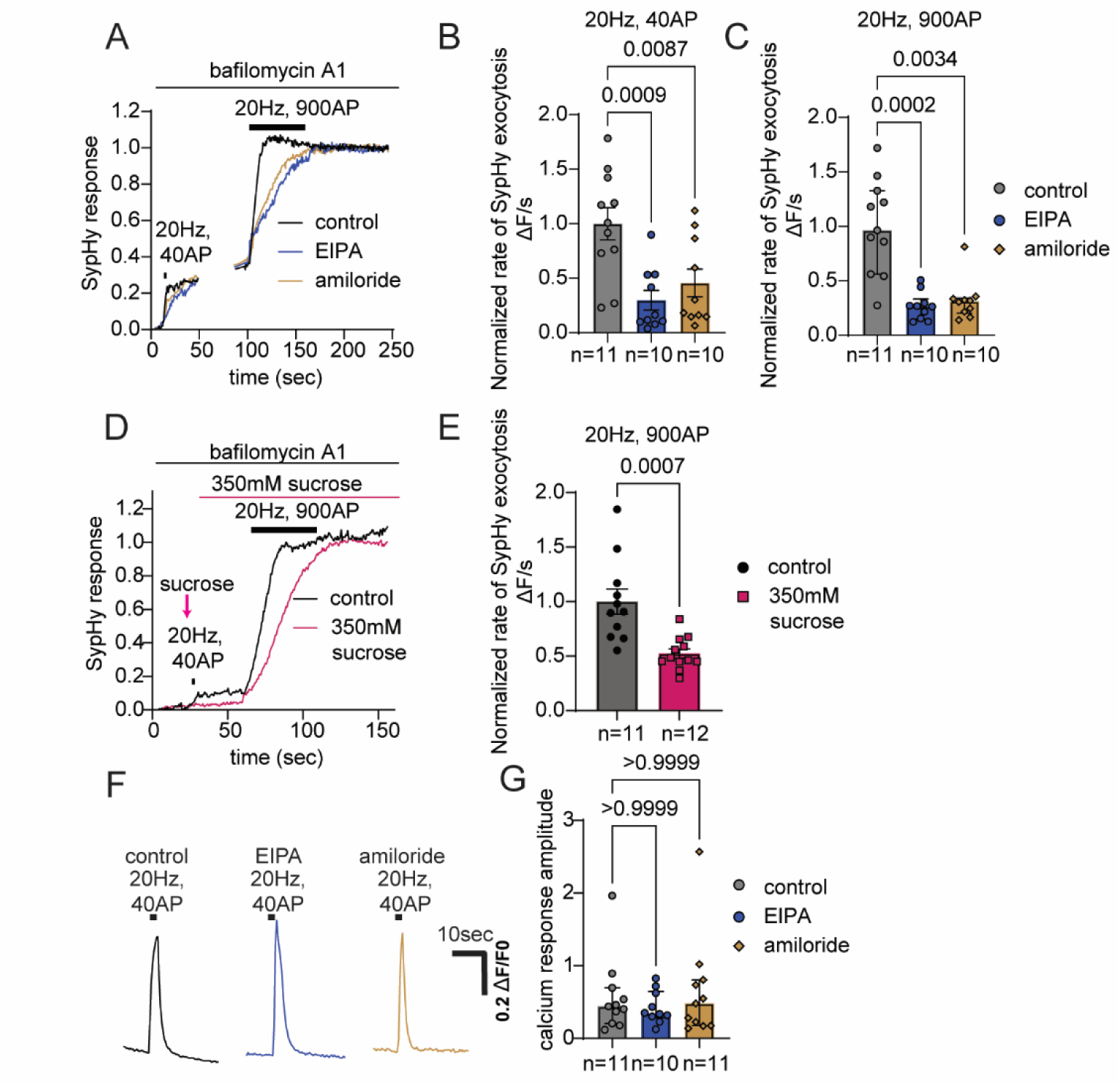
Blocking presynaptic filopodia mimics reduced membrane tension, slowing SV fusion. (**A-C**) Hippocampal neurons transfected with SypHy were treated with vehicle, 50 µM EIPA, or 1 mM amiloride for 30 min before live imaging. Neurons were stimulated with trains of 40 and 900AP (20Hz) in the presence of the drugs and 1 μM bafilomycin A1. (**A**) Average SypHy response traces (ΔF/F_0_) normalized to the second response amplitude. (**B,C**) Normalized SypHy exocytosis rate (ΔF/s) following the first (**B**) or second (**C**) stimulus. p values, **B**, one-way ANOVA with Dunnett’s test, **C**, Kruskal-Wallis with Dunn’s test. (**D,E**) Primary cultures of hippocampal neurons were transfected with SypHy. The neurons were either perfused with a mild hypertonic solution containing 350 mM sucrose or stimulated with 40AP, 20Hz (indicated by bar), both in the presence of 1 μM bafilomycin A1. After this, both control and sucrose-treated neurons were stimulated with 900AP, 20Hz. (**D**) Example traces of SypHy response normalized to the second response amplitude. (**E**) SypHy exocytosis rates (ΔF/s); p values, unpaired t-test. (**F,G**) Neurons transfected with GCaMP6f were incubated with 50 µM EIPA or 1 mM amiloride, then stimulated with 40AP, 20Hz. (**F**) Example GCaMP6f responses (ΔF/F_0_). (**G**) GCaMP6f response amplitudes; p values: Kruskal-Wallis test.

**Fig. S6.**
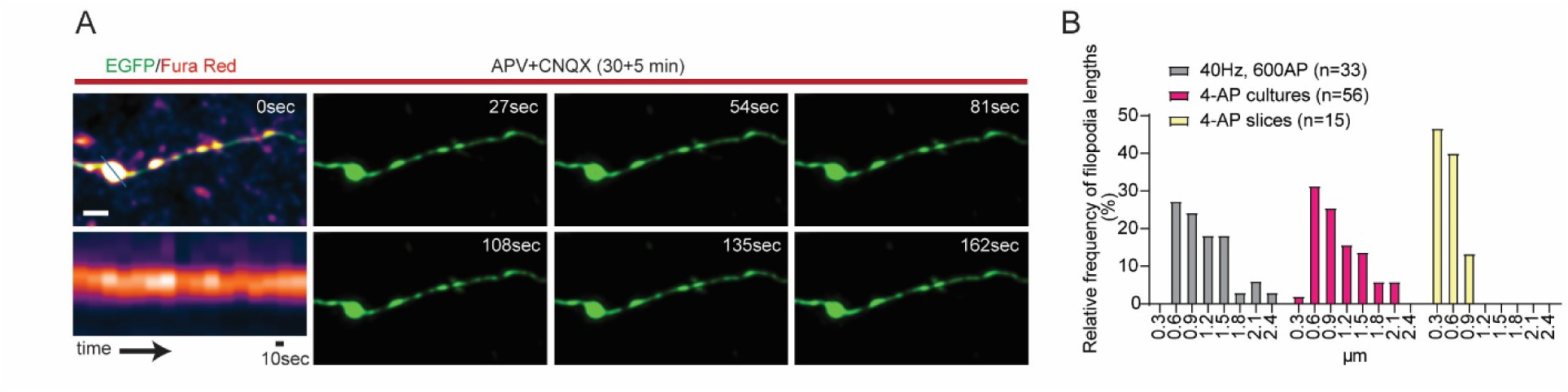
Neuronal activity is required for presynaptic filopodia formation in intact brain tissue. (**A**) Time-lapse imaging of EGFP-labelled neurons in acute brain slices from a Thy1-GFPM mouse. Brain slices were loaded with Fura Red, incubated with 50 µM APV and 10 µM CNQX for 30 min. Confocal imaging shows no filopodia formation at putative synaptic boutons. Kymographs show uniform Fura Red calcium signals in regions marked by blue lines. Scale bar 1 µm. (**B**) Frequency histogram showing the length (µm) of presynaptic filopodia induced by electrical field stimulation or 4-AP in dissociated neuronal cultures, and by 4-AP in acute brain slices.

**Fig. S7.**
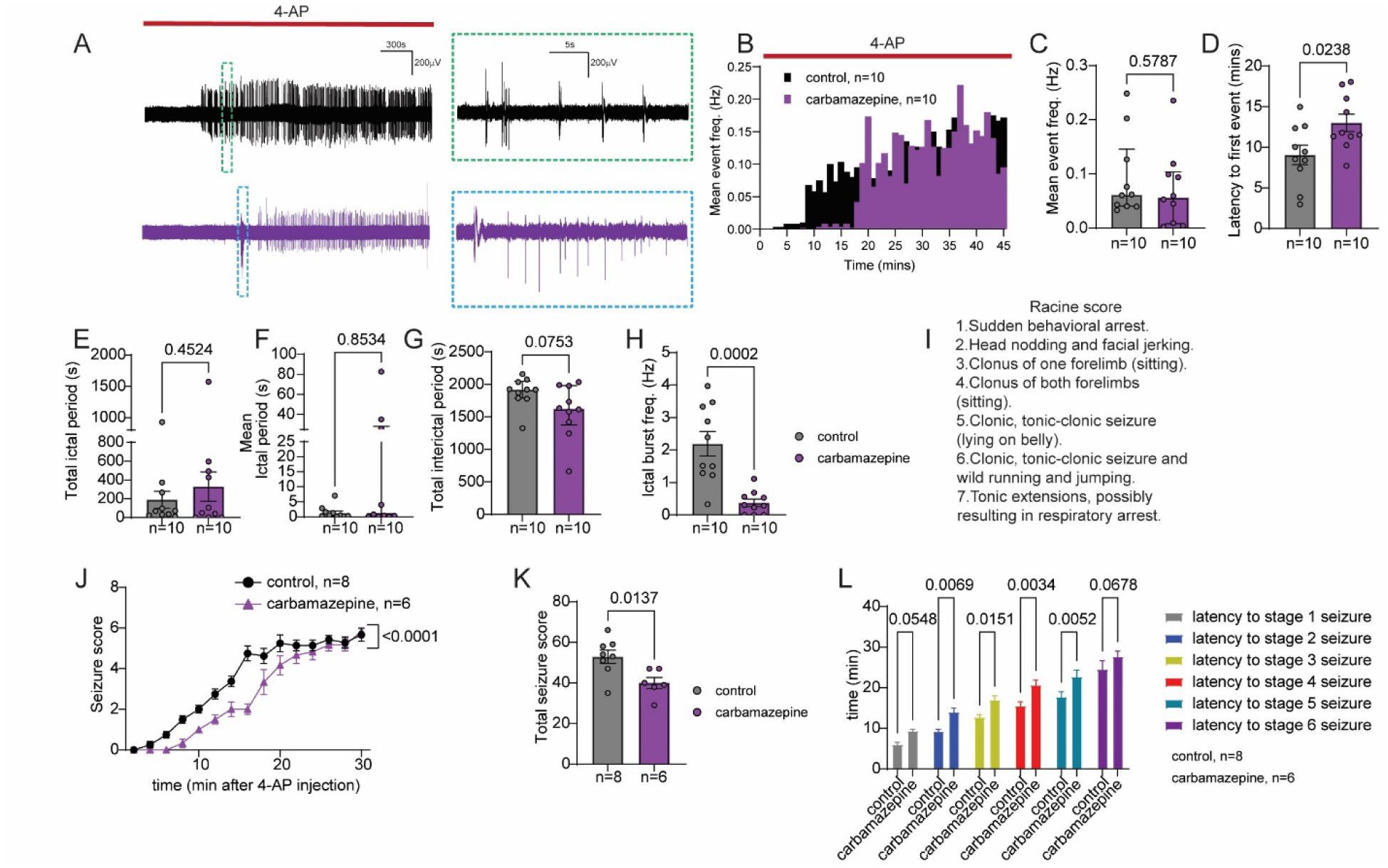
Effect of the ASM carbamazepine on 4-AP-induced seizures *in vitro* and *in vivo.* (**A-H**) Slices incubated with and without 80 µM carbamazepine for 45 min. Epileptiform activity was elicited by 100 µM 4-AP and recorded in the CA1 pyramidal layer. (**A**) Example traces of interictal and ictal-like discharges in control and carbamazepine-treated slices. (**B**) Time-frequency histograms of epileptiform discharges during 4-AP wash-in. (**C-H**) Analysis of epileptiform event frequency (**C**), latency to first event (**D**), total (**E**) and mean (**F**) duration of ictal-like activity, total duration of interictal-like activity (**G**) and burst frequency during ictal-like events (**H**). **C, E, F, G,** p values, Mann-Whitney U test. **D, H,** p values, unpaired t-test. (**I-L**) Carbamazepine (20 mg/kg) or vehicle was injected into 8-10-week-old wild-type mice 30 min before 4-AP (6.65 mg/kg) to induce behavioral seizures. Data for vehicle control as in **Fig. 6P-R**. (**I**) Modified Racine scale for scoring seizure severity. (**J**) Progression of average seizure severity after 4-AP injection in treated and untreated mice. p values, two-way ANOVA with Sidak’s test. (**K**) Total seizure score. p values, unpaired t-test. (**L**) Latency to different seizure stages. p values, two-way ANOVA with Fisher’s LSD.

## Fiji code “Find responding synapses and set ROIs”

run(“Clear Results”);

//run(”Close All”);

//open(“”);

run(“Remove Slice Labels”);

rename(“timelapse”);

run(“Set Scale…”, “distance=0 known=0 pixel=1 unit=pixel”);

run(“Z Project…”, “start=26 stop=30 projection=[Average Intensity]”);

rename(“stim.tif”);

selectWindow(“timelapse”);

run(“Z Project…”, “start=11 stop=15 projection=[Average Intensity]”);

rename(“baseline.tif”);

imageCalculator(“Subtract create”, “stim.tif”,“baseline.tif”);

run(“Find Maxima…”, “noise=400 output=[Point Selection] exclude”);

run(“Measure”);

a=newArray(0);

b=newArray(0);

for(i=0;i<nResults;i=i+1) {

x=getResult(“X”,i);

y=getResult(“Y”,i);

a = Array.concat(a,x);

b= Array.concat(b,y);

}

for (i=0;i<a.length;i=i+1) {

makeOval(a[i]-5,b[i]-5,10,10);

if (i<a.length) {

roiManager(“Add”);

}

}

---

**Table S1.**
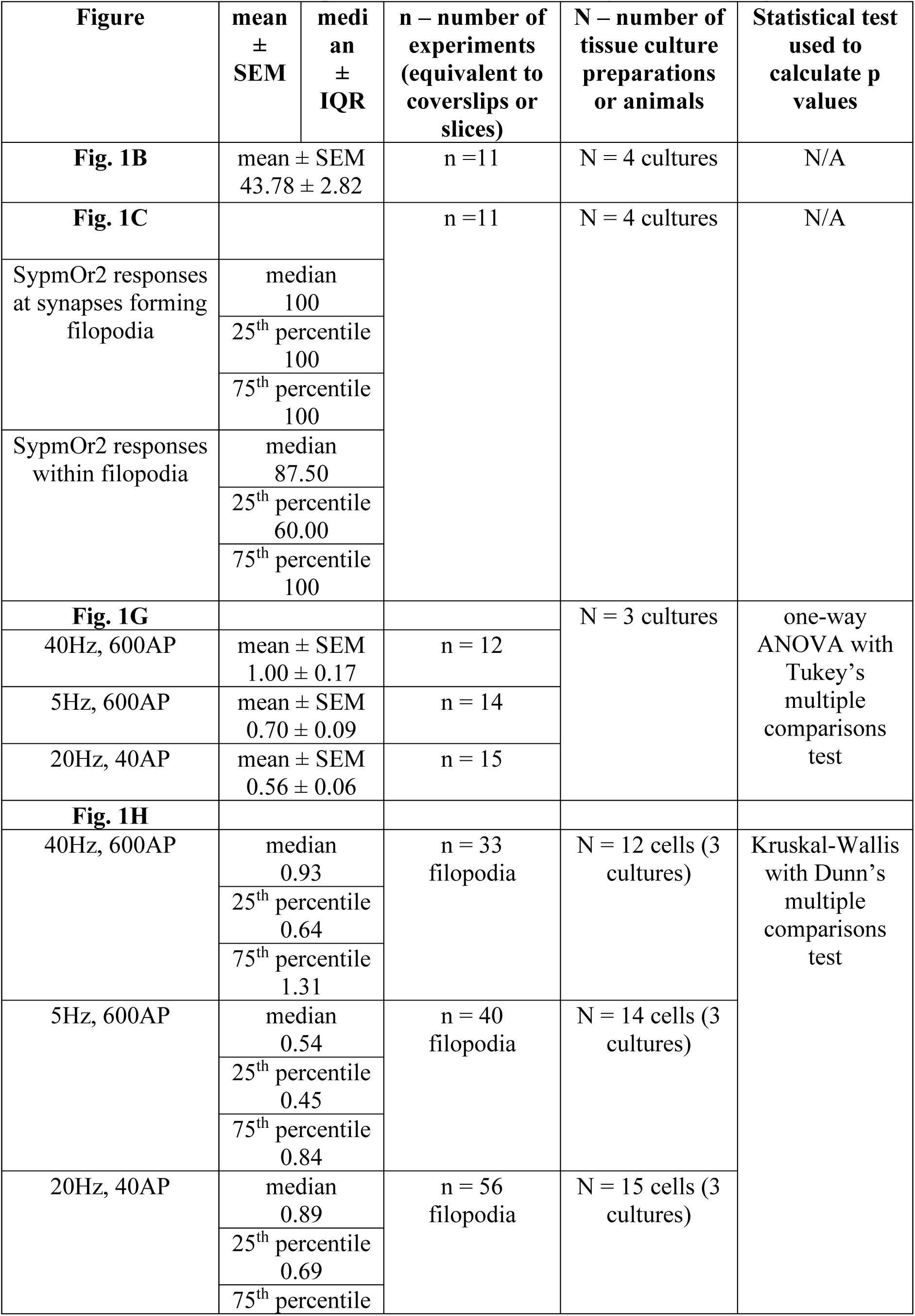
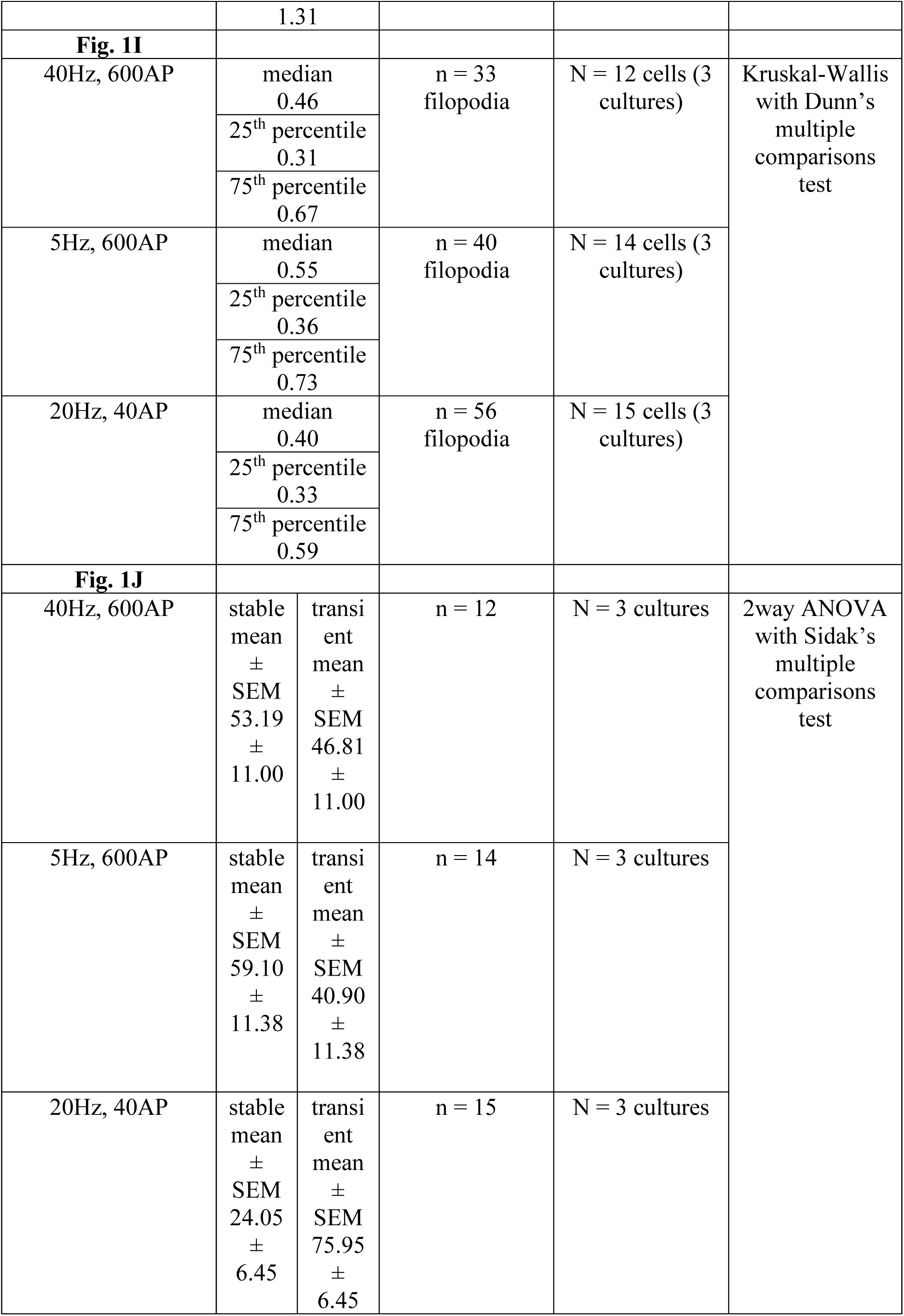
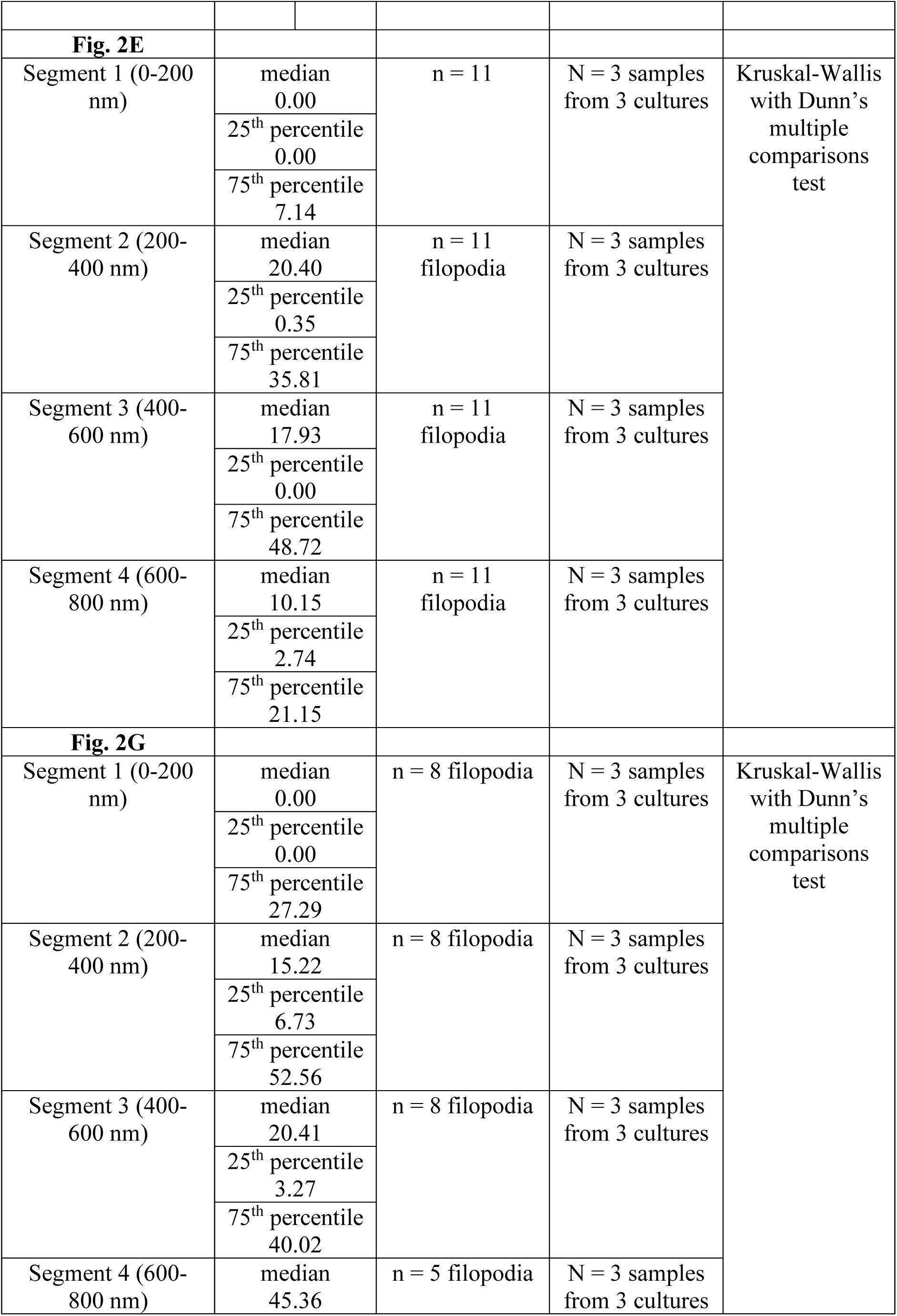
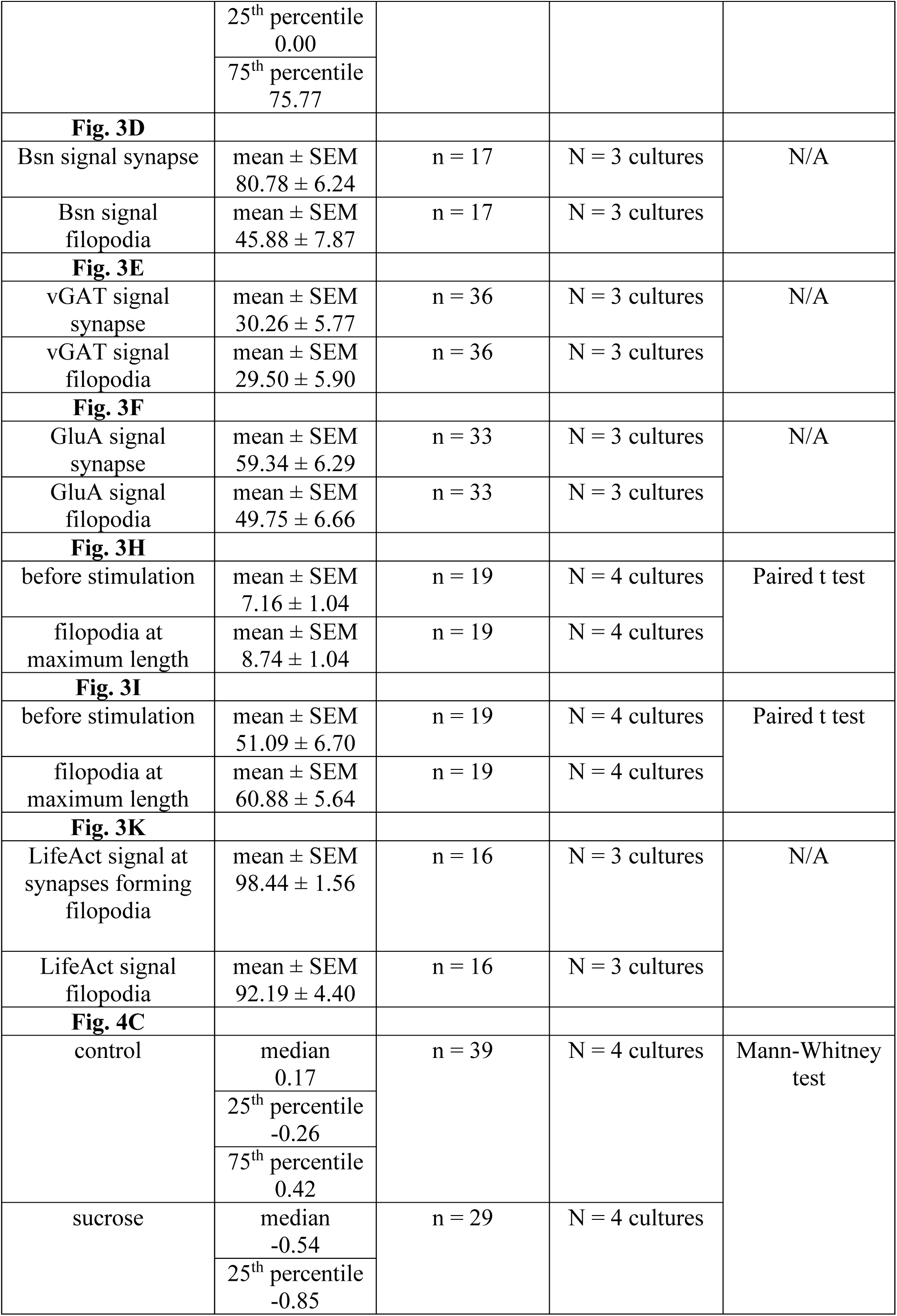
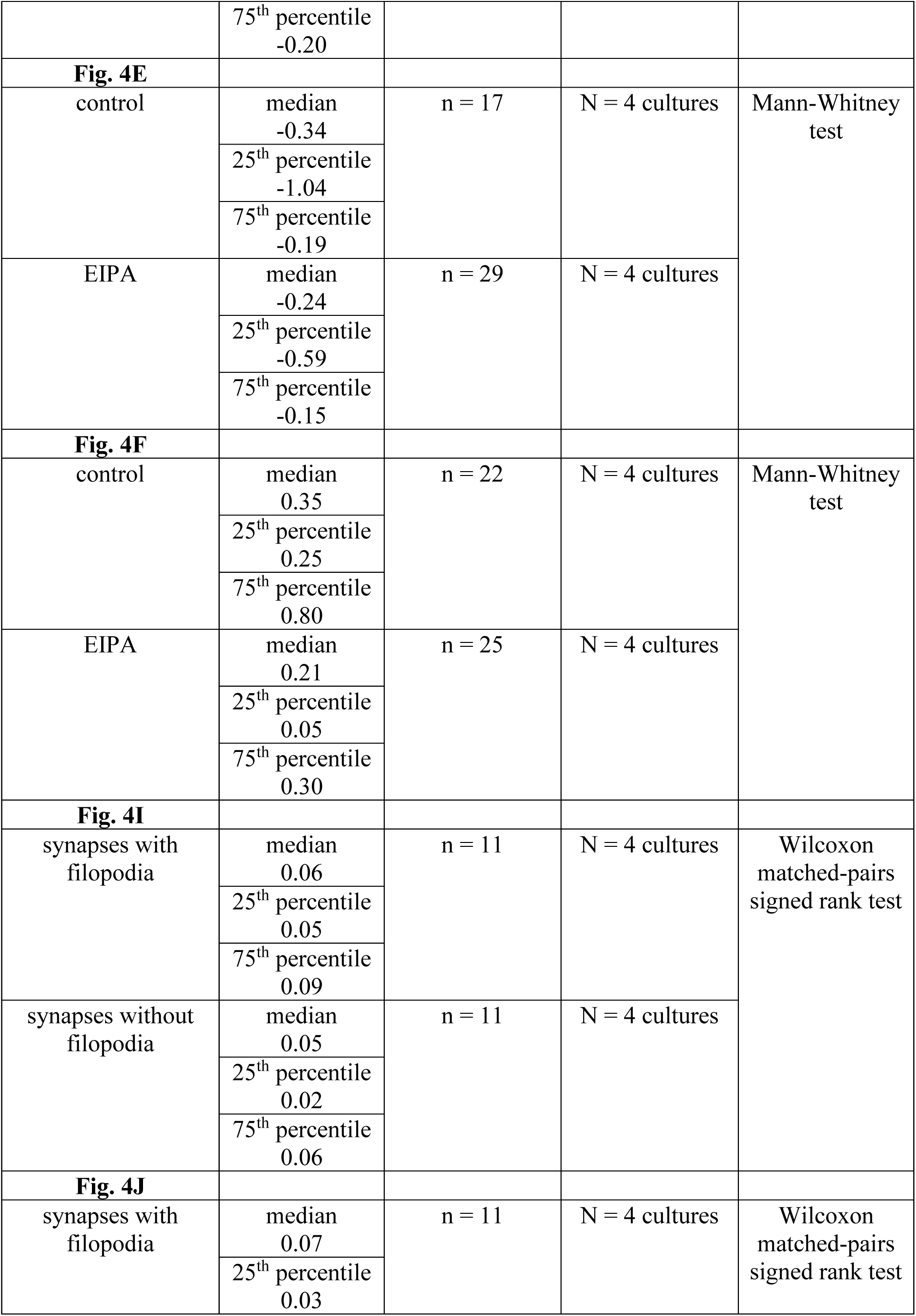
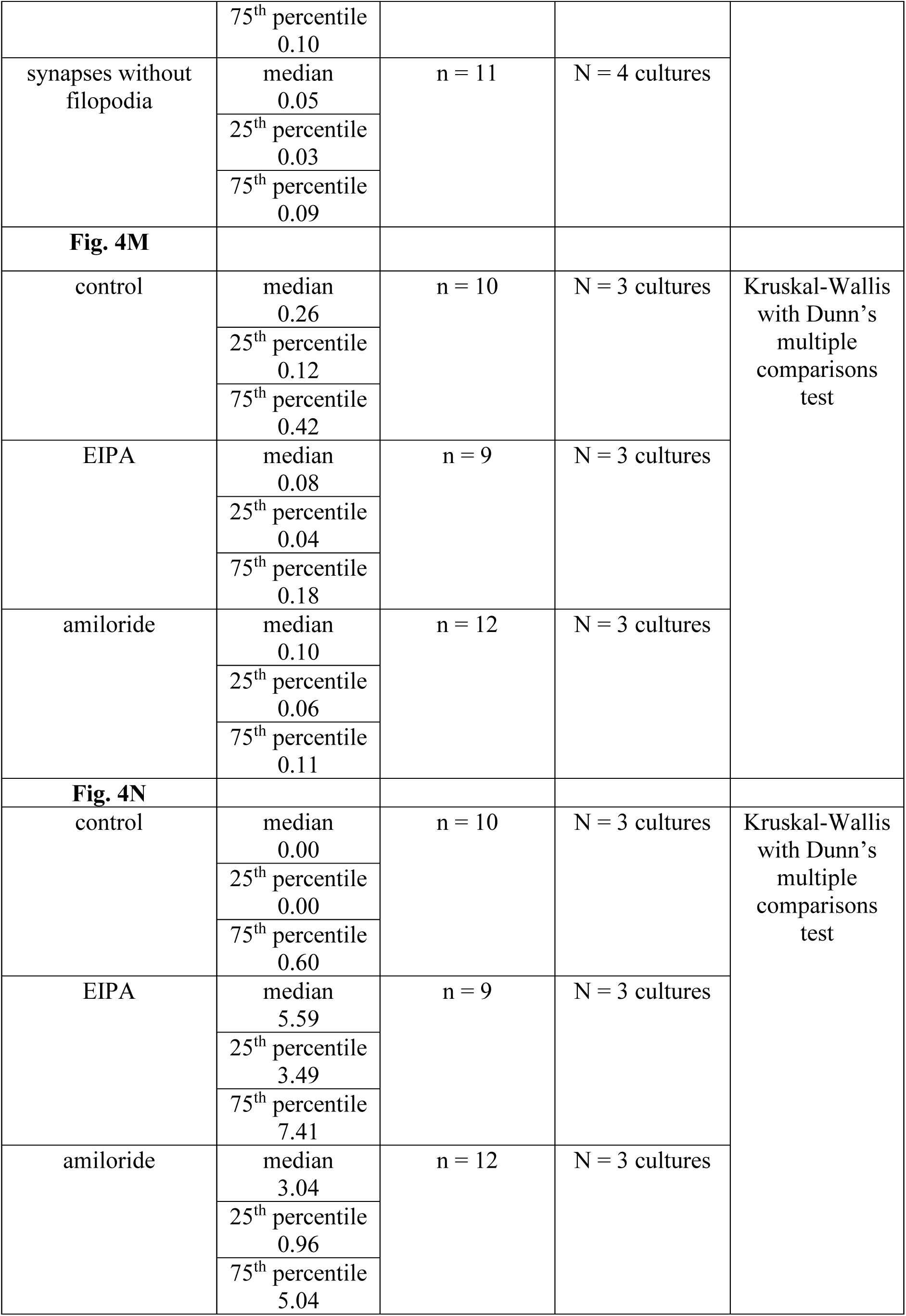
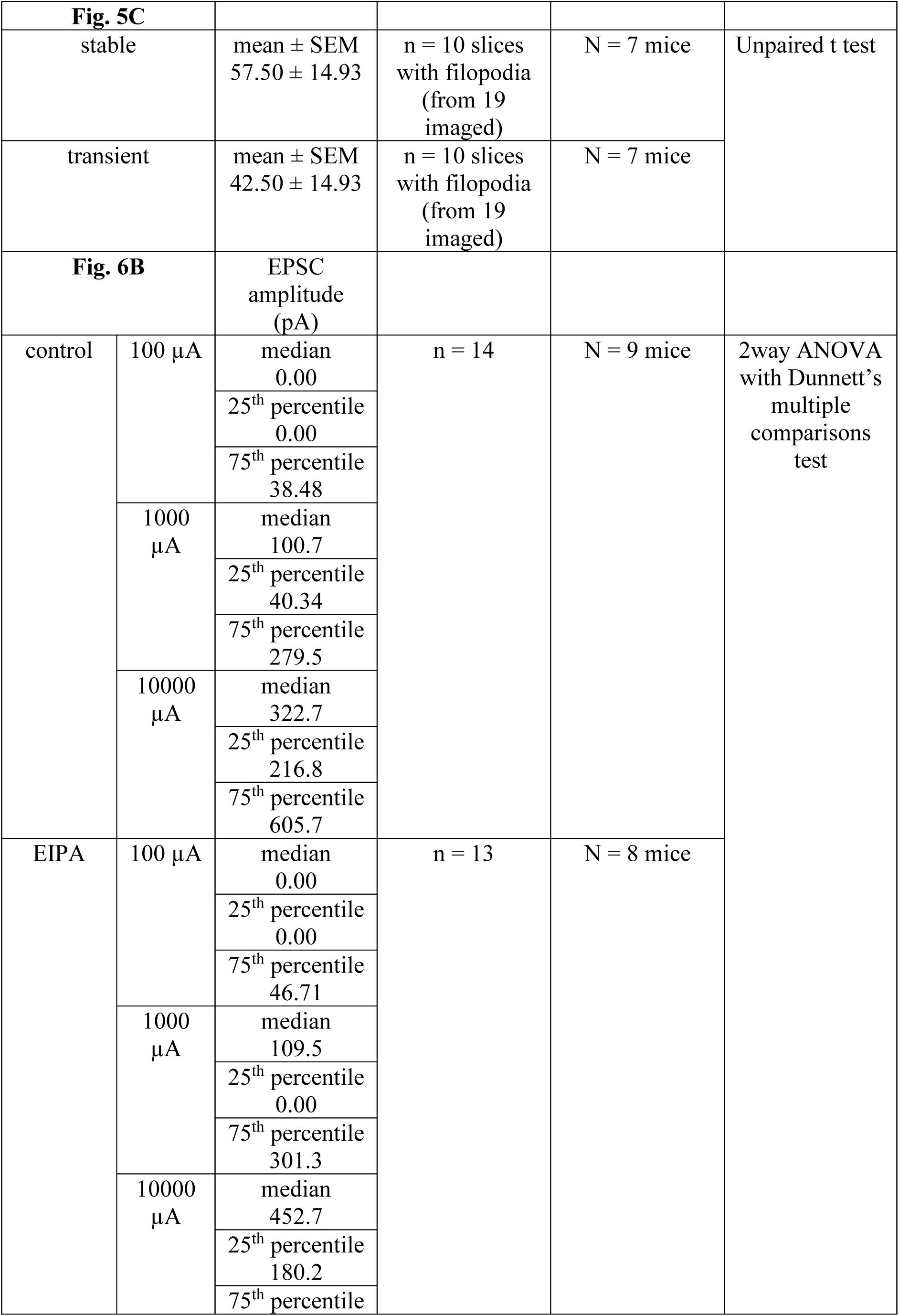
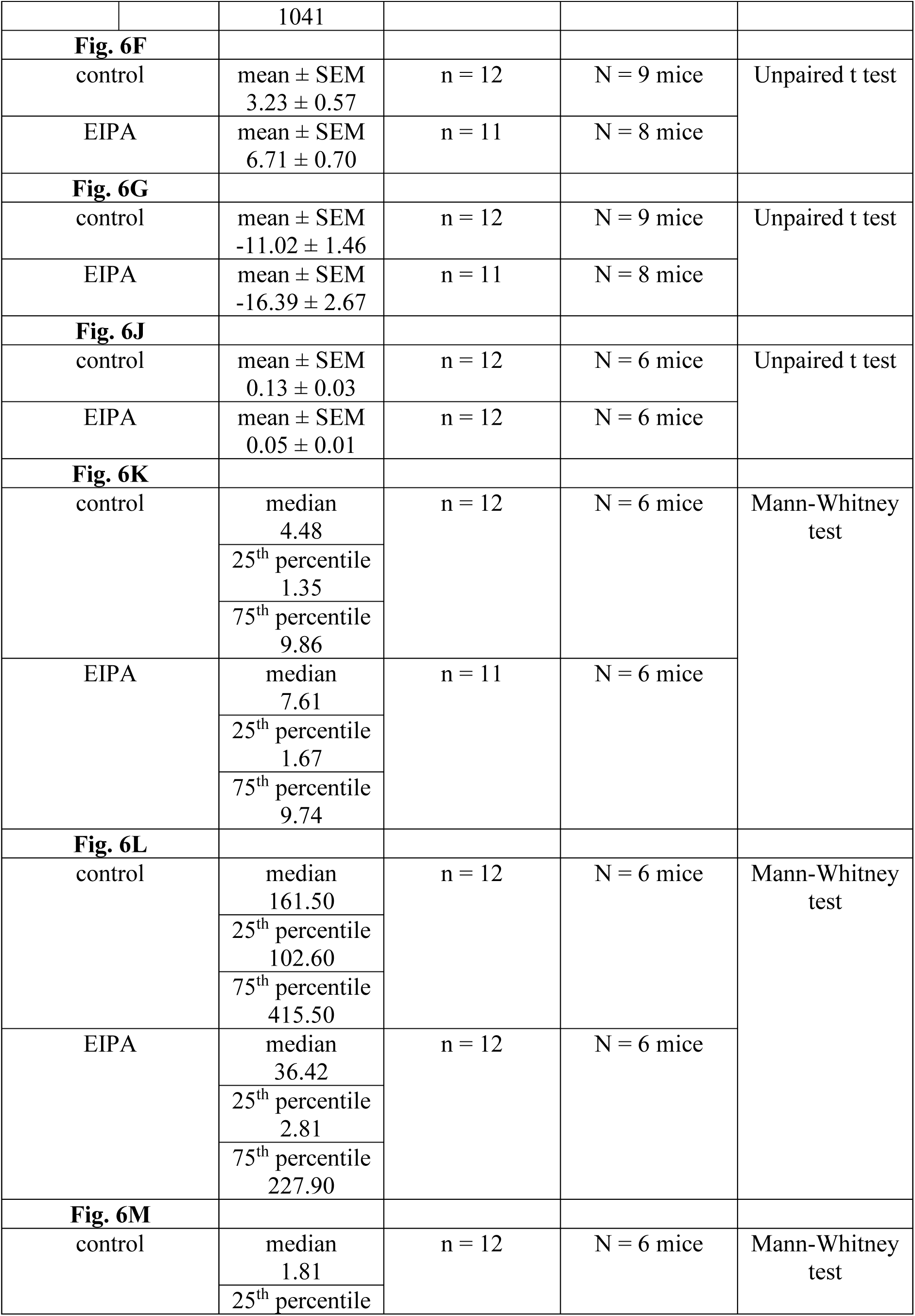
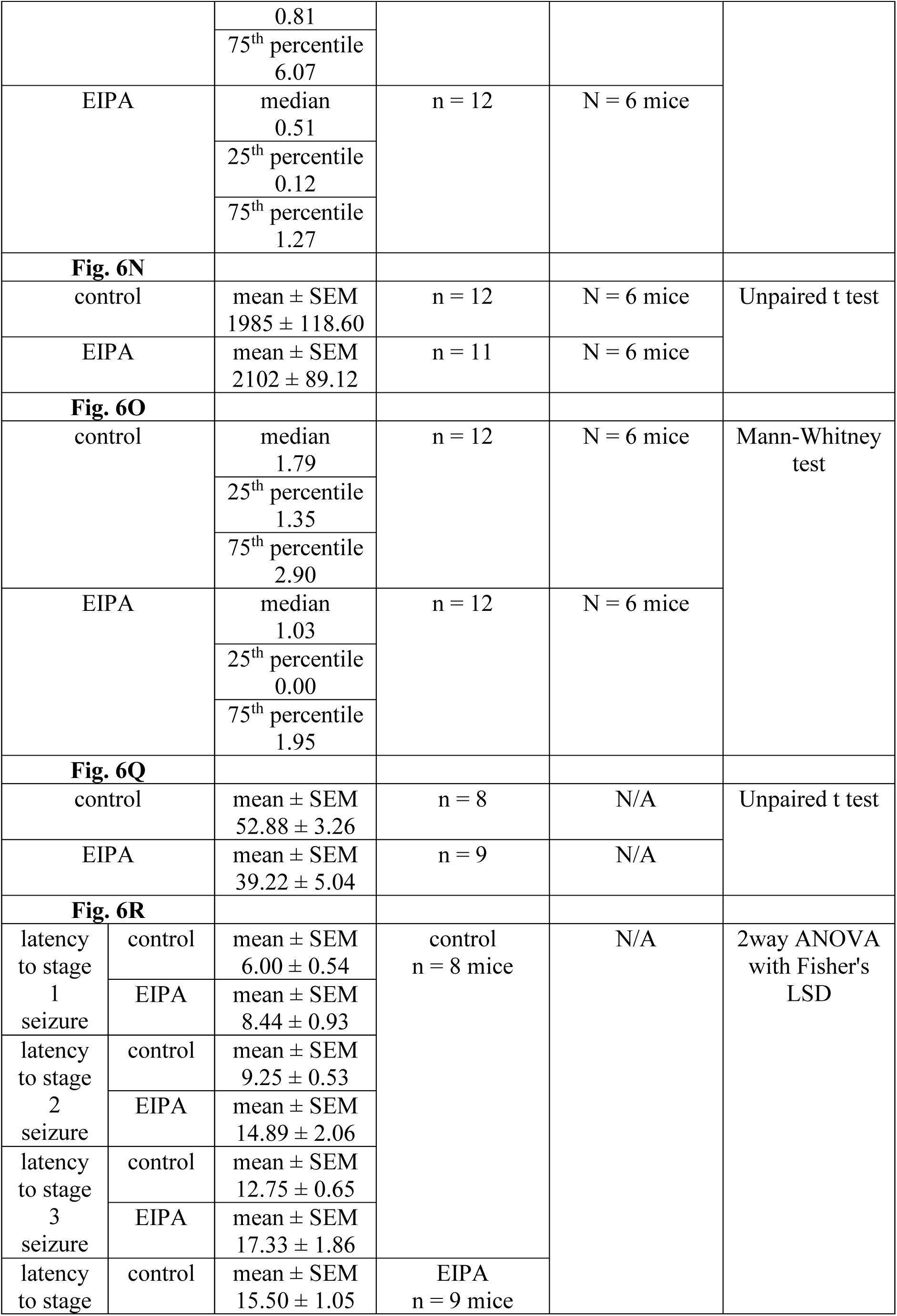
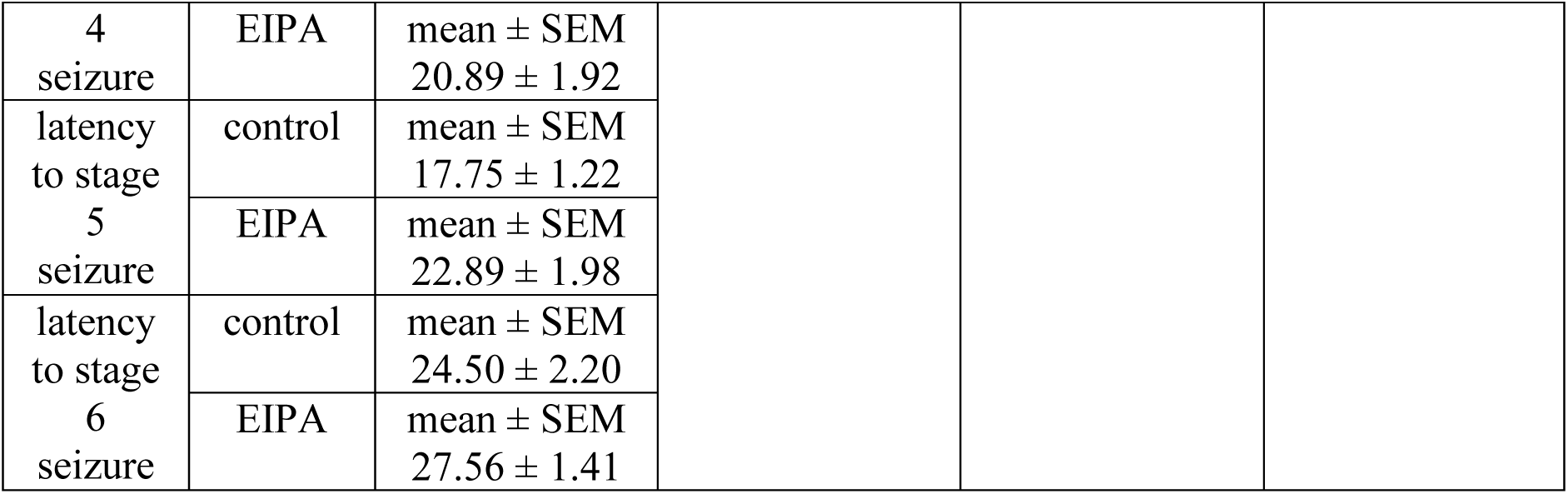
Numerical values, sample sizes and statistical analysis for main figures.

**Table S2.**
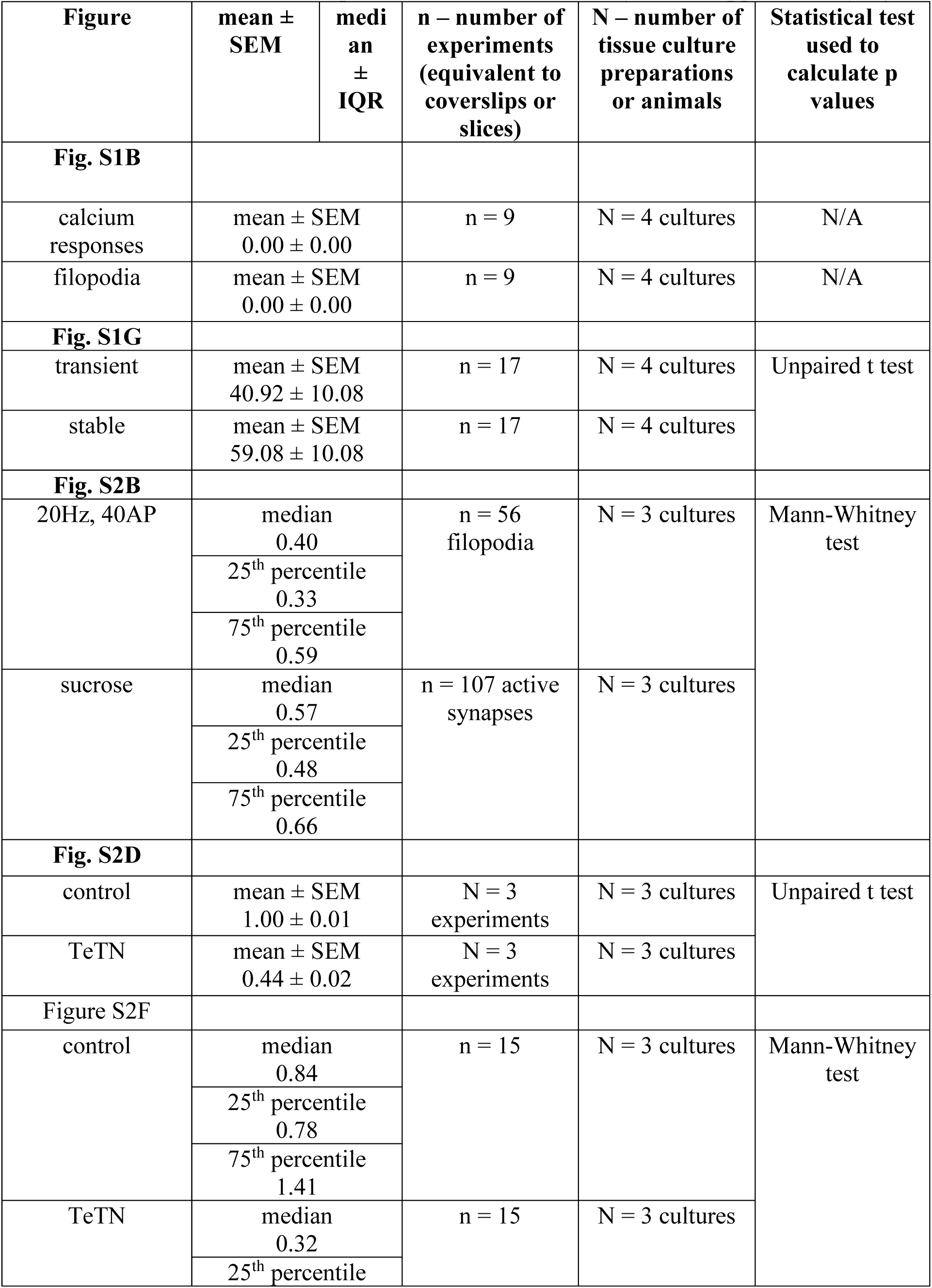
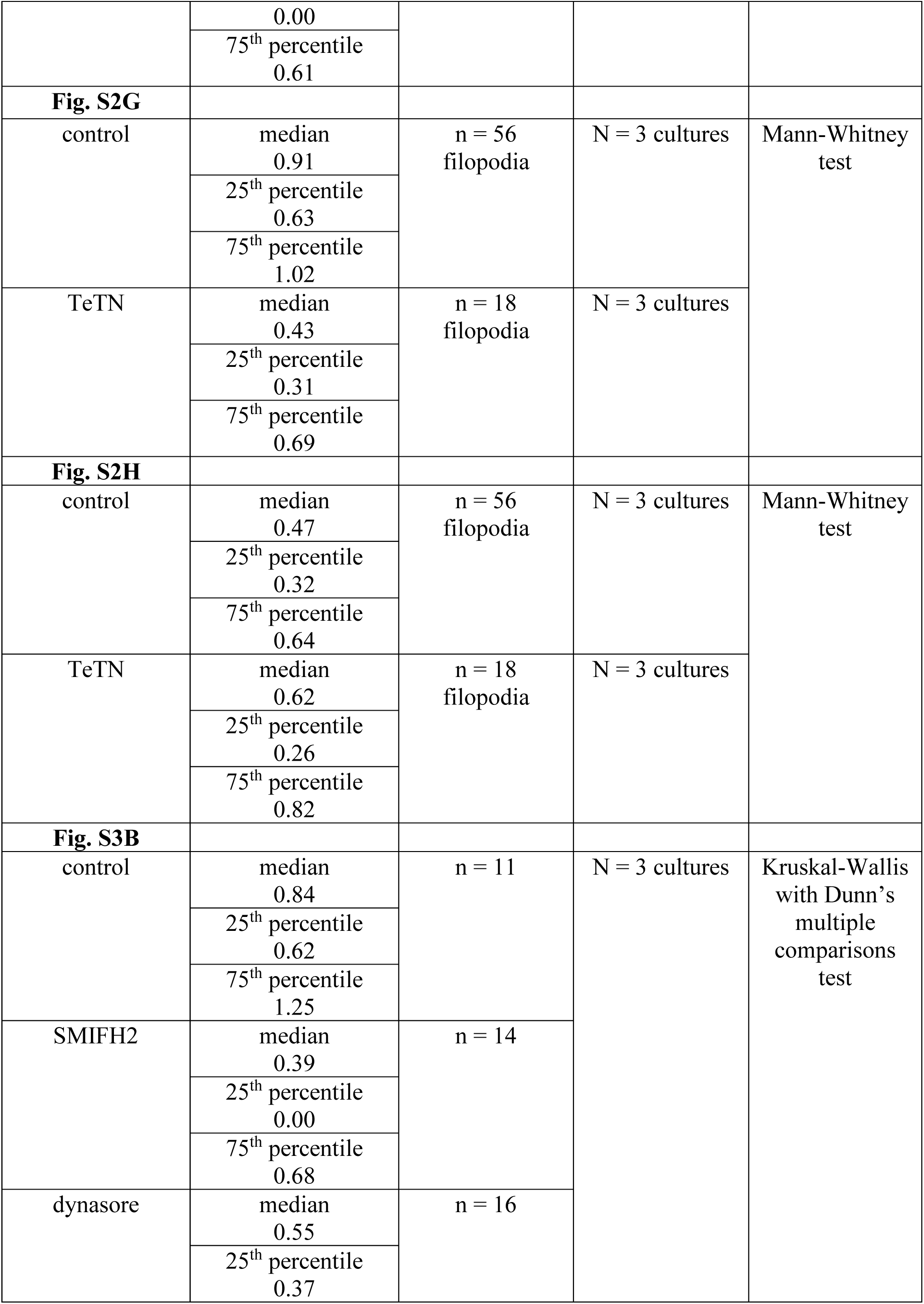
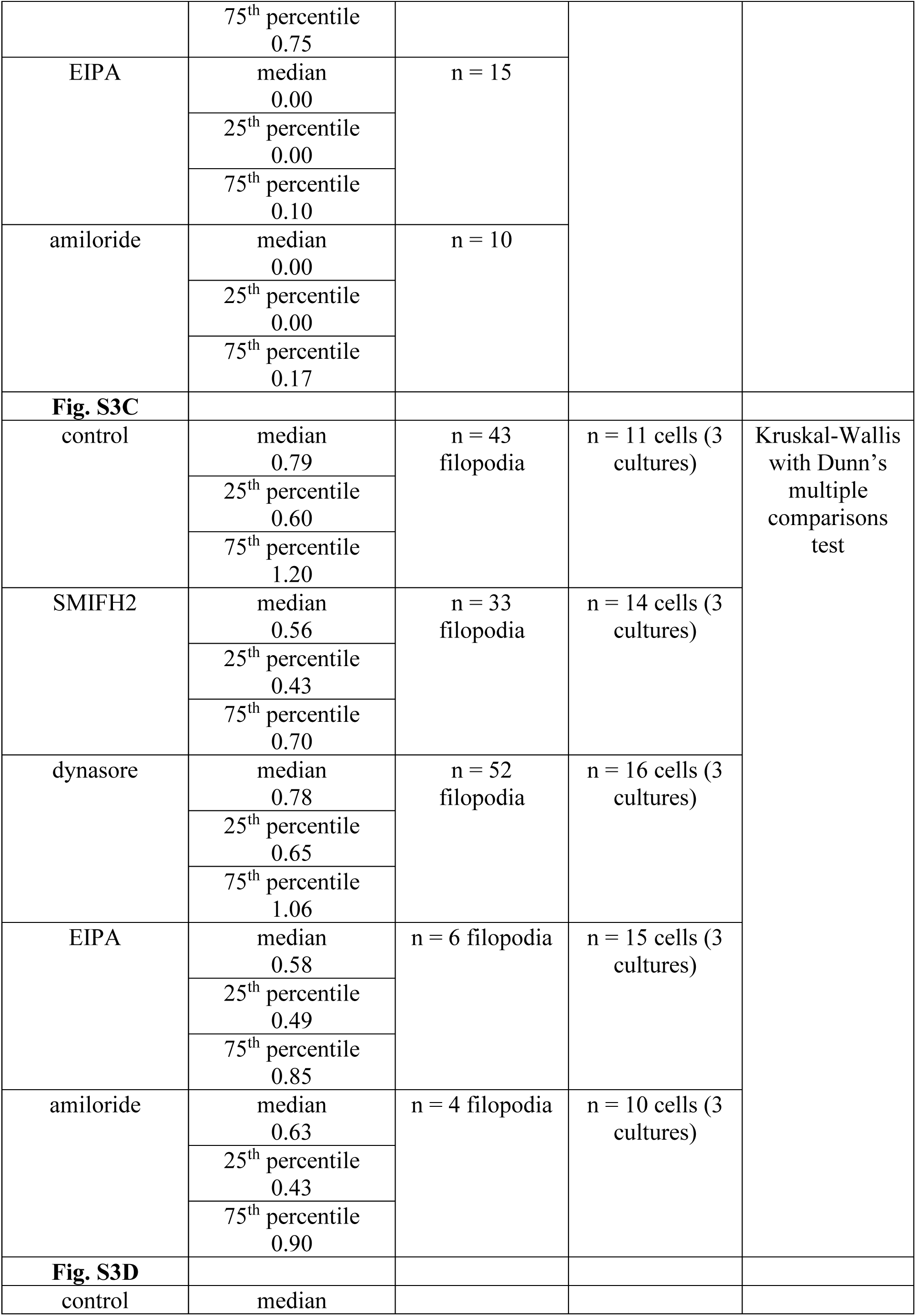
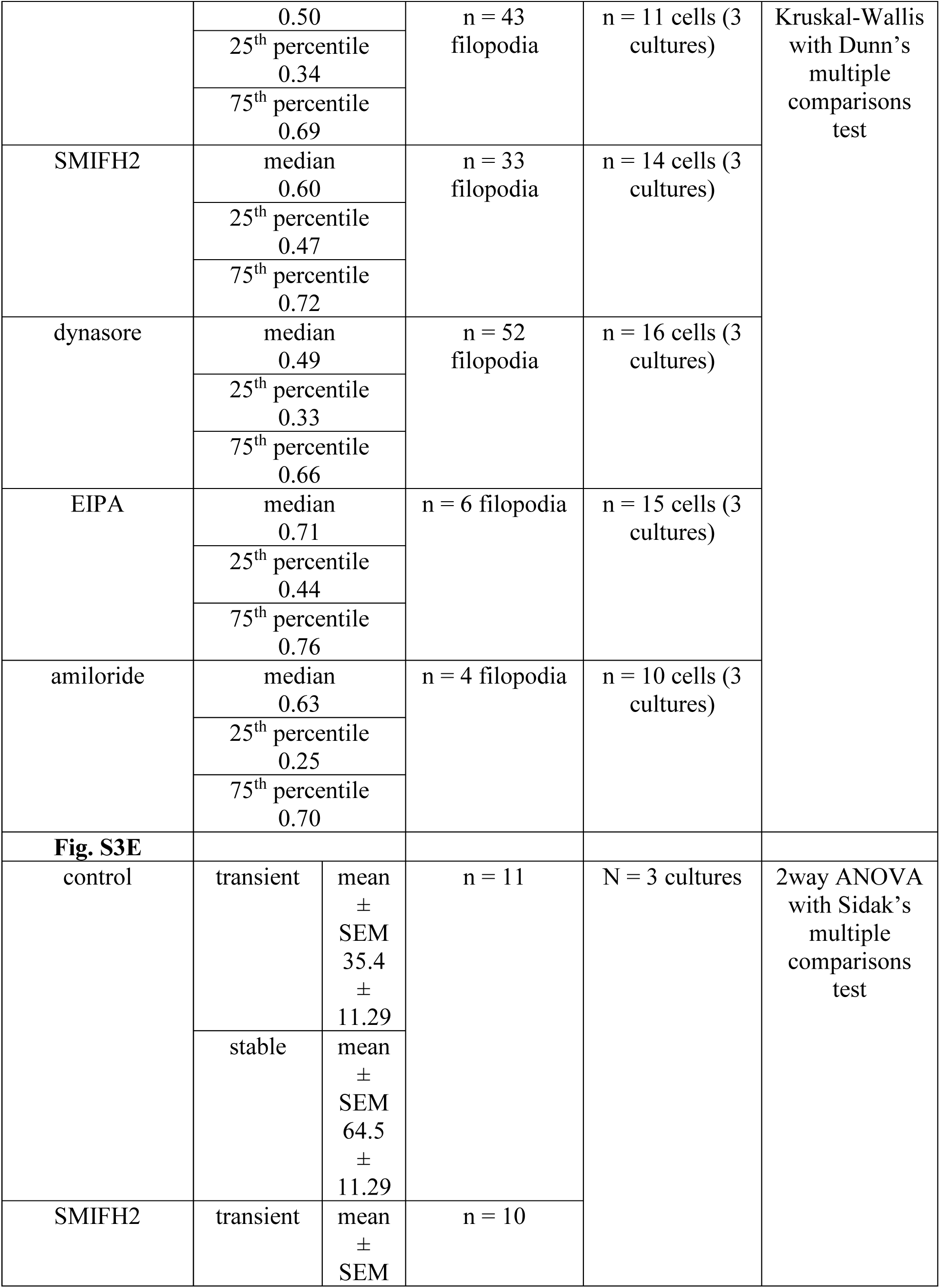
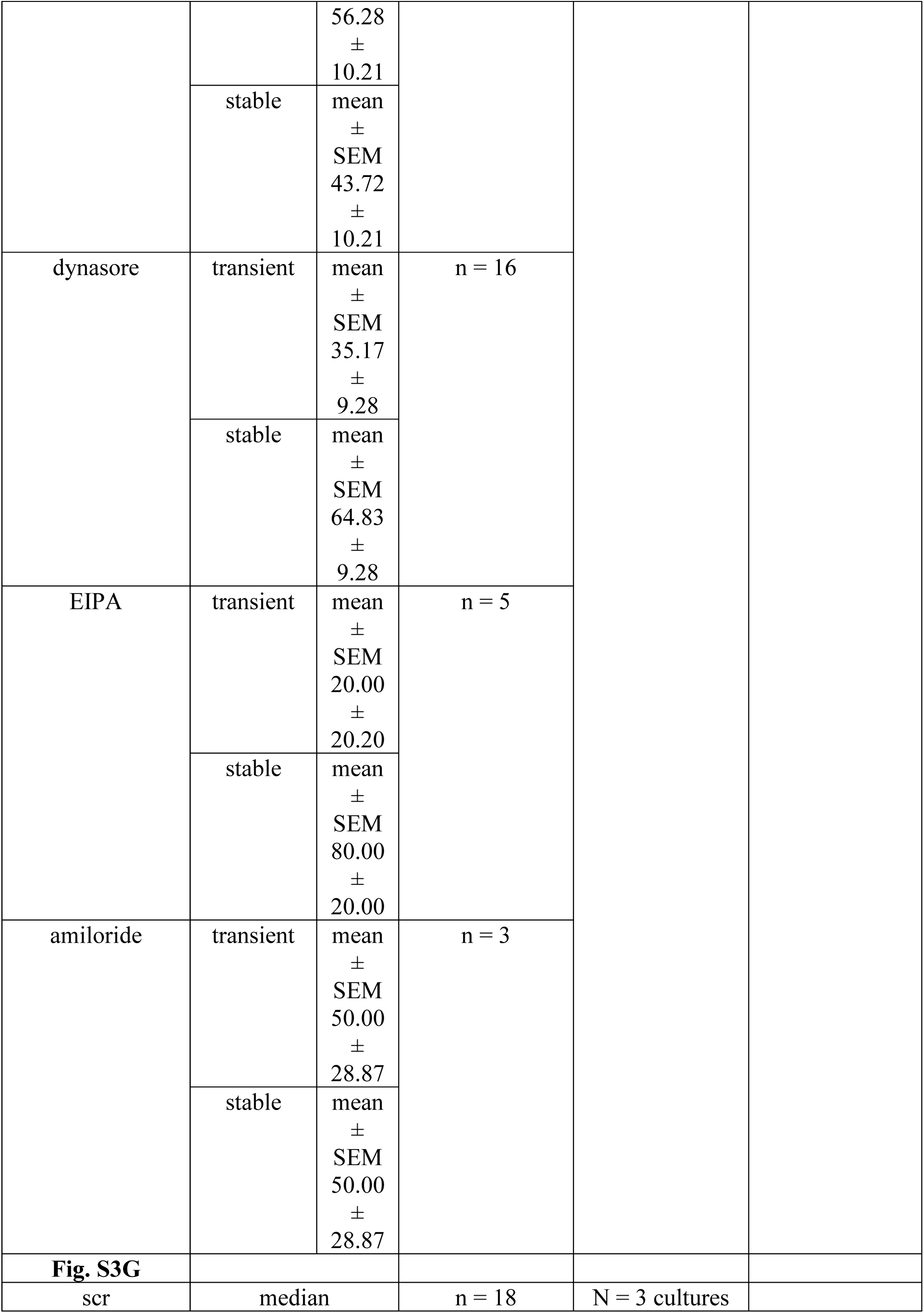
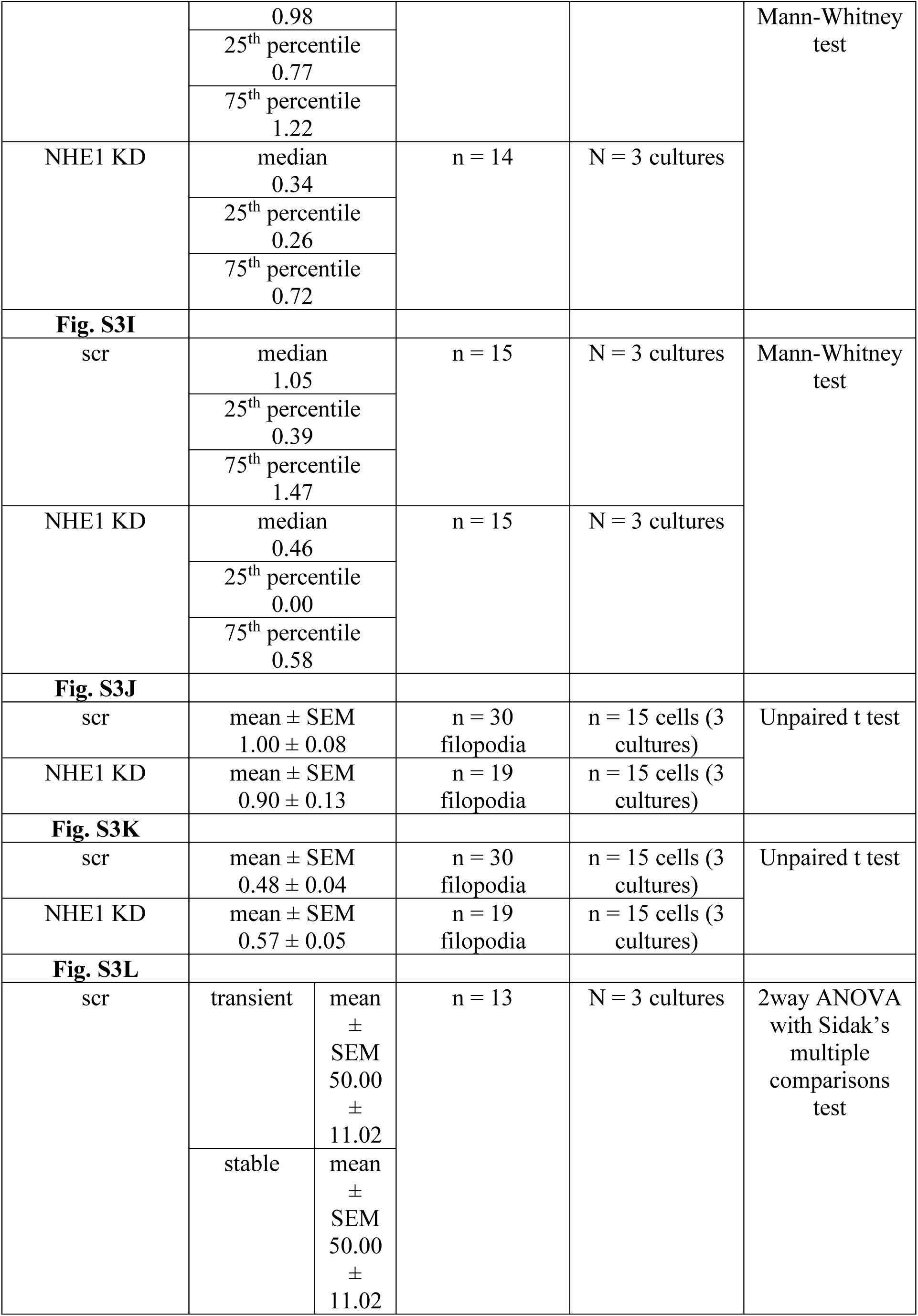
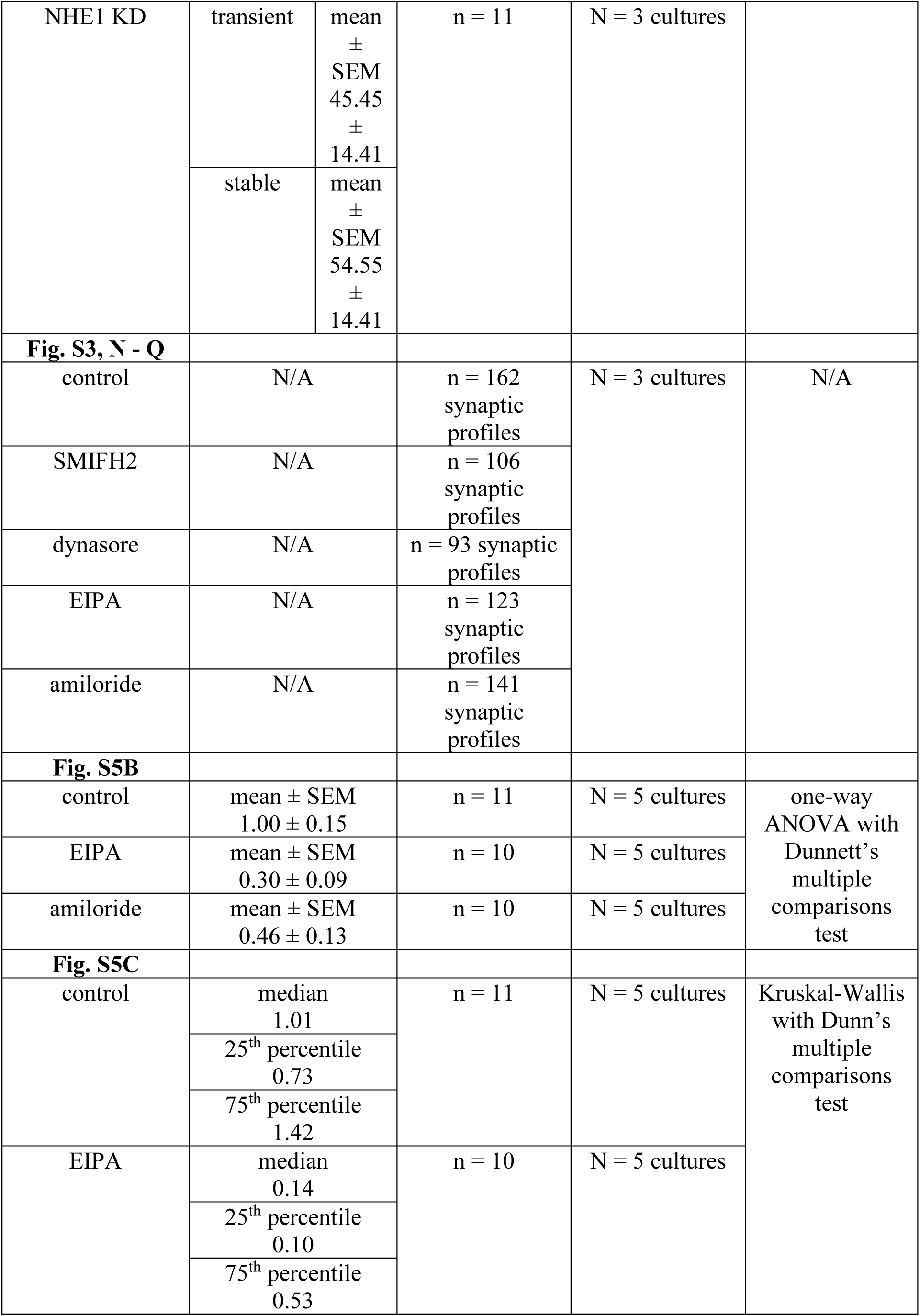
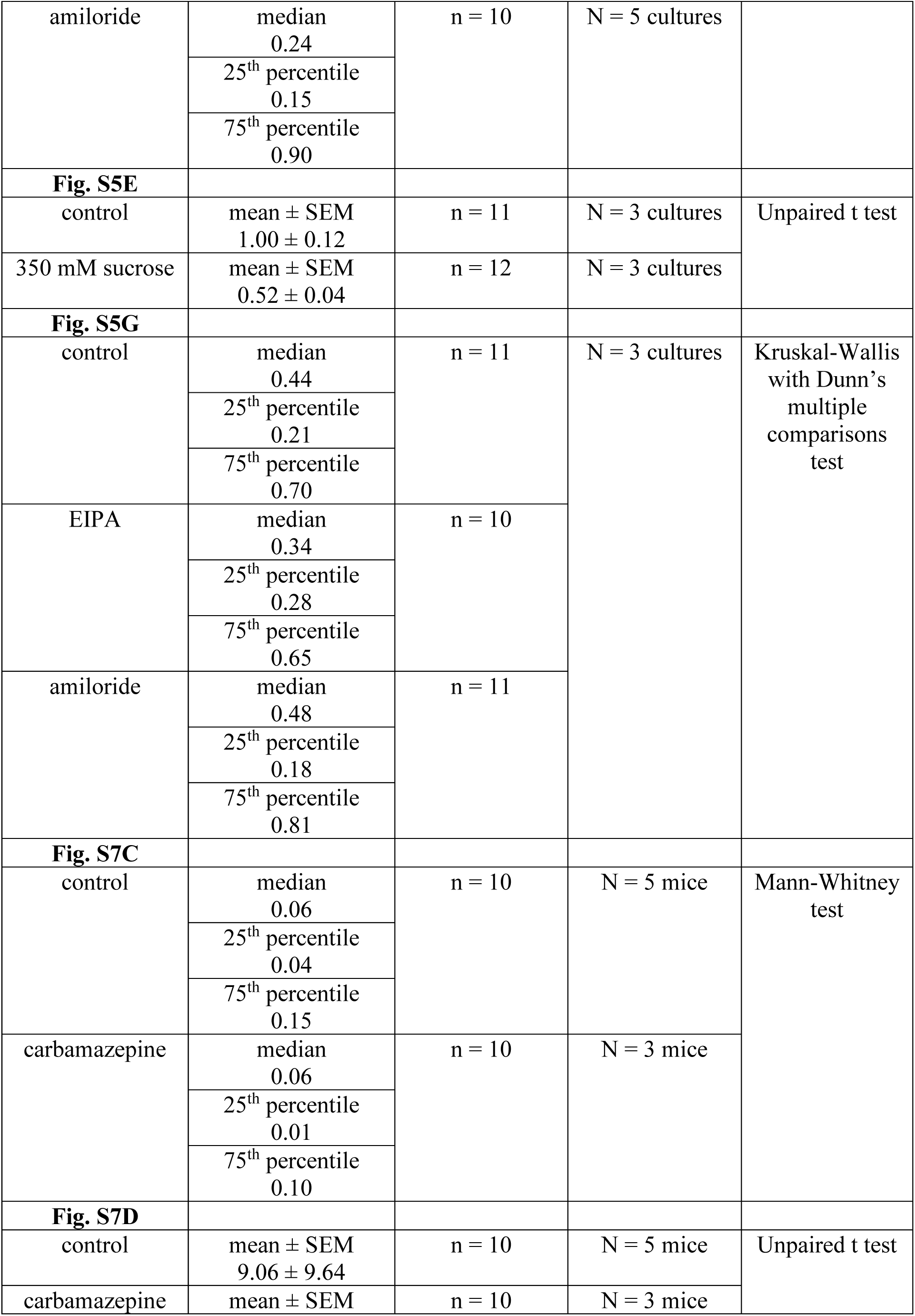
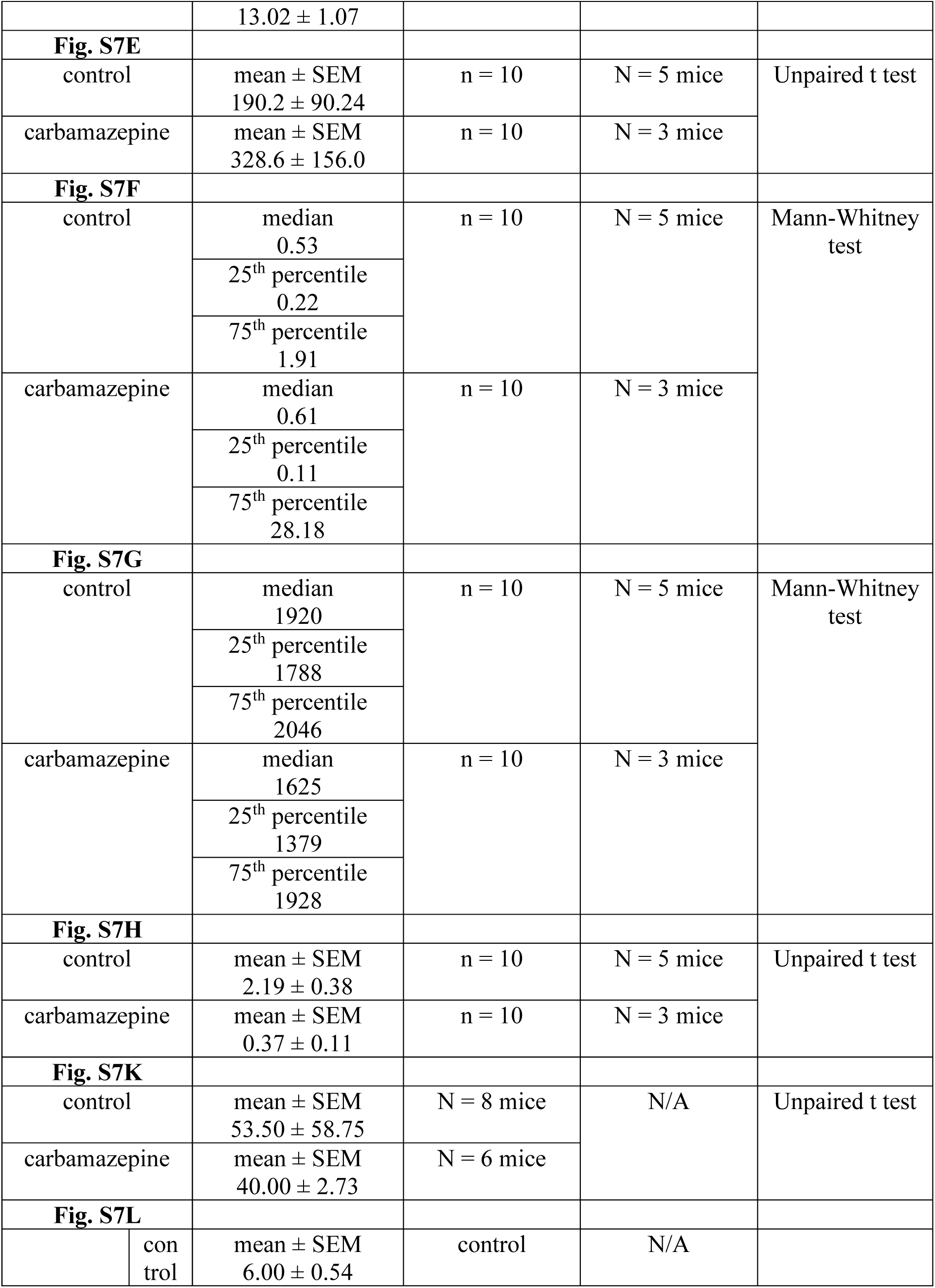
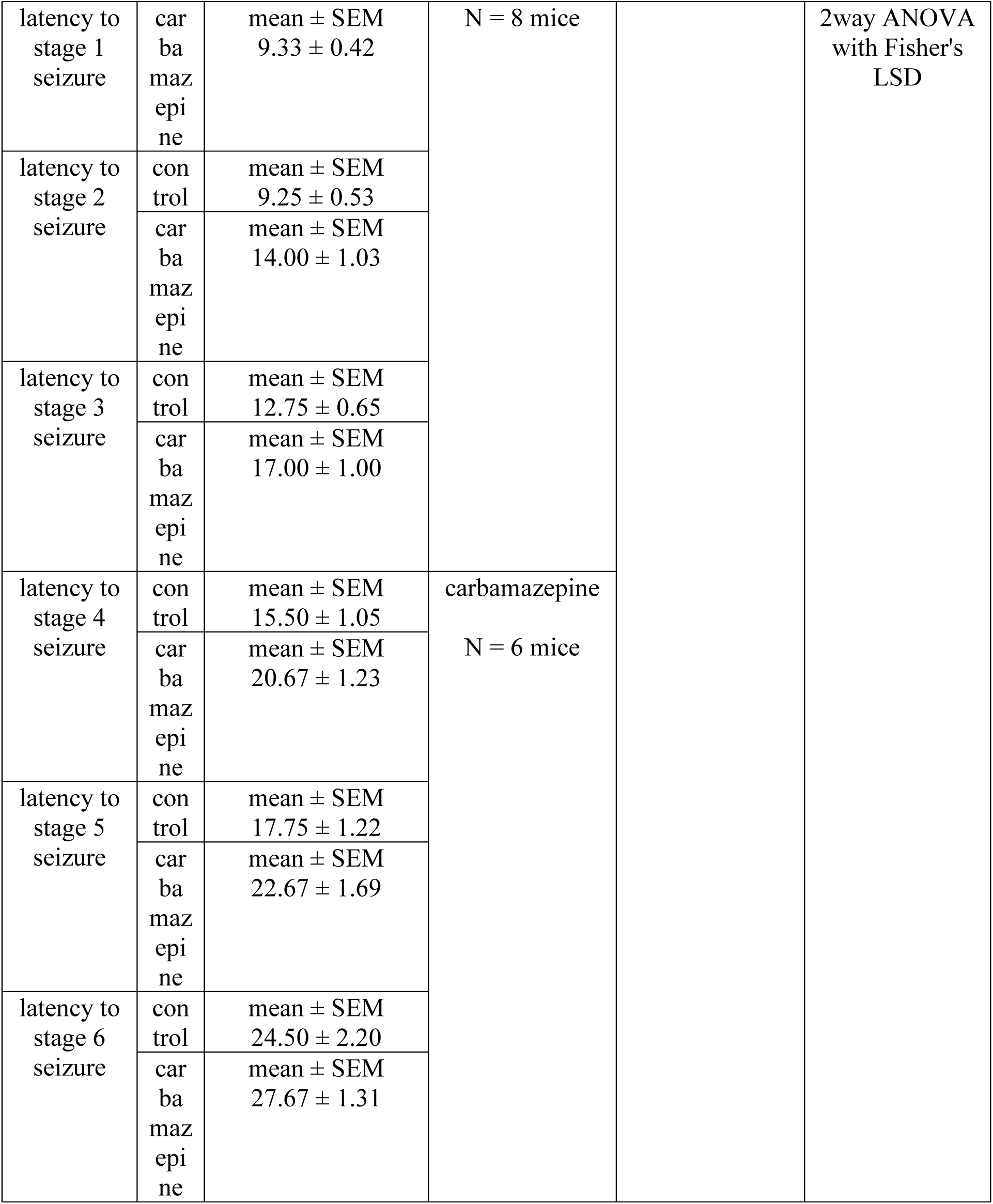
Numerical values, sample sizes and statistical analysis for supplementary figures.

